# MYC overrides HIF to regulate proliferating primary cell metabolism in hypoxia

**DOI:** 10.1101/2020.09.21.306464

**Authors:** Courtney A. Copeland, Benjamin A. Olenchock, David R. Ziehr, Sarah McGarrity, Kevin Leahy, Jamey D. Young, Joseph Loscalzo, William M. Oldham

## Abstract

Hypoxia requires metabolic adaptations to sustain energetically demanding cellular activities. While the metabolic consequences of hypoxia have been studied extensively in cancer cell models, comparatively little is known about the metabolic response of primary cells to hypoxia. We performed metabolic flux analyses of proliferating human lung fibroblasts and pulmonary artery smooth muscle cells in hypoxia. Unexpectedly, hypoxia decreased glycolytic flux despite activation of hypoxia-inducible factor (HIF) and increased glycolytic enzyme expression. Pharmacologic activation of HIF with the prolyl hydroxylase (PHD) inhibitor molidustat in normoxia did increase glycolytic flux, but hypoxia abrogated this effect. Multi-omic profiling revealed distinct molecular responses to hypoxia and pharmacologic PHD inhibition and suggested a critical role for MYC in modulating the HIF response in hypoxia. MYC knockdown in hypoxia increased lactate efflux, while MYC overexpression in normoxia blunted the effects of molidustat treatment. Together, these data suggest that other factors, notably MYC, supersede the anticipated effects of HIF-dependent up-regulation of glycolytic gene expression on glycolytic flux in hypoxic proliferating primary cells.

## INTRODUCTION

Cellular responses to hypoxia propel many physiologic and pathologic processes from wound healing and angiogenesis to vascular remodeling and fibrosis (Semenza, 2012; Lee *et al*, 2019a). These activities require cells to continue energetically demanding tasks, such as macromolecular biosynthesis and proliferation, despite limited oxygen availability. Since respiration is the most efficient way for cells to produce energy, cell metabolism must adapt to meet energetic needs when oxygen supply is limiting. Understanding how these metabolic adaptations sustain critical cellular processes in hypoxia is fundamentally important to our understanding of human health and disease.

Cells typically respond to hypoxia by shifting energy production away from respiration and toward glycolysis. This response is mediated primarily by stabilization of the hypoxia-inducible transcription factor 1α (HIF-1α). HIF-1α activates the transcription of glucose transporters, glycolytic enzymes, lactate dehydrogenase, and pyruvate dehydrogenase kinase, while decreasing the expression of tricarboxylic acid (TCA) cycle and electron transport chain enzymes (Semenza, 2012; Lee *et al*, 2020). Although HIF-1α is constitutively expressed, it is hydroxylated by prolyl hydroxylase enzymes (PHDs) in normoxia and targeted for proteasomal degradation. PHDs are the principal oxygen sensors in metazoan cells (Kaelin & Ratcliffe, 2008). PHDs are α-ketoglutarate-dependent dioxygenase enzymes that require molecular oxygen for their enzymatic activity. When oxygen tension falls, PHD activity decreases, leading to HIF-1α stabilization and activation of its associated transcriptional program. Overall, this transcriptional program should increase glycolytic capacity and divert glucose-derived pyruvate from oxidative phosphorylation toward lactate fermentation to maintain glycolytic ATP production.

In addition to metabolic changes designed to maintain energy supply, hypoxic cells also reduce energy demand through down-regulation of Na^+^/K^+^-ATPase, slowing protein translation, and decreasing cell proliferation (Wheaton & Chandel, 2011; Hubbi & Semenza, 2015). In particular, HIF-1α decreases cell proliferation by activating cyclin-dependent kinase inhibitor expression, inhibiting cell-cycle checkpoint progression (Gardner *et al*, 2001), and antagonizing pro-proliferative MYC signaling (Koshiji *et al*, 2004). Despite these canonical effects of HIF-1α activation, there are many examples where cells continue to proliferate despite hypoxic stress, including cancer cells, stem cells, and lung vascular cells (Hubbi & Semenza, 2015). How these cells meet the metabolic needs of sustained proliferation in hypoxia and how these adaptations are regulated are active areas of investigation (Jain *et al*, 2020; Oldham *et al*, 2015; Lee *et al*, 2020). Since hypoxia is a prominent feature of cancer biology as tumor growth outstrips blood supply, most detailed metabolic studies of hypoxic cell metabolism have used tumor cell models, yielding important insights into the metabolic pathobiology of cancer (Wise *et al*, 2011; Metallo *et al*, 2011; Melendez-Rodriguez *et al*, 2019; Garcia-Bermudez *et al*, 2018; Lee *et al*, 2019b; Jiang *et al*, 2016). For example, stable isotope tracing and metabolic flux analyses identified a critical role for the reductive carboxylation of glutamine-derived α-ketoglutarate for lipid biosynthesis in supporting tumor growth (Metallo *et al*, 2011; Gameiro *et al*, 2013b; Scott *et al*, 2011; Wise *et al*, 2011), and metabolomic studies identified aspartate as a limiting metabolite for cancer cell proliferation under hypoxia (Garcia-Bermudez *et al*, 2018). By contrast, comparatively little is known about metabolic adaptations of primary cells to hypoxia. Indeed, the importance of reductive carboxylation or aspartate biosynthesis remains to be elucidated in these cells. A more complete understanding of primary cell metabolic adaptations to hypoxia would provide an important context for understanding how metabolic reprogramming supports normal cellular responses to hypoxia, how these responses may be (mal)adaptive in a variety of disease contexts, and how the hypoxia metabolic program in primary cells differs from that observed in cancer cells.

To address these questions, we have developed models of bioenergetic carbon flux in human lung fibroblasts (LFs) and pulmonary artery smooth muscle cells (PASMCs) cultured in 21% or 0.5% oxygen. These cells may be exposed to a wide range of oxygen concentrations *in vivo*, continue to proliferate despite hypoxic culture conditions *in vitro*, and play important roles in the pathology of non-cancerous diseases in which tissue hypoxia is a prominent feature, including pulmonary hypertension and pulmonary fibrosis. We found that hypoxia fails to increase glycolysis in these primary cells despite robust up-regulation of the HIF-1α transcriptional program. In normoxia, HIF-1α stabilization by the PHD inhibitor molidustat (BAY-85-3934, “BAY”) (Flamme *et al*, 2014) did increase glycolysis and lactate efflux; however, hypoxia blocked this response. These findings suggested that important hypoxia-dependent regulatory mechanisms override the metabolic consequences of HIF-1α-dependent glycolytic gene expression. Transcriptomic profiling identified a critical role for the transcription factor MYC in the adaptive response to hypoxia. Using knockdown and overexpression approaches, we demonstrated that MYC attenuates HIF-driven glycolysis in hypoxia and following HIF stabilization in normoxia.

## RESULTS

### Hypoxia uncouples HIF-dependent glycolytic gene expression from glycolytic metabolic flux

The goal of this study was to characterize hypoxia-induced metabolic changes in proliferating primary LFs and PASMCs. To accomplish this goal, we adopted a metabolic flux analysis technique that enabled us to link intracellular metabolic fluxes to cell proliferation rates. Metabolic flux analysis fits cell proliferation rate, extracellular flux measurements, and ^13^C intracellular isotope labeling patterns to a computational model of cell metabolism (Antoniewicz, 2018). This analysis reconstructs comprehensive flux maps that depict the flow of carbon from extracellular substrates, through intracellular metabolic pathways, and into cell biomass and metabolic by-products (Young, 2014). These models assume that cells are at a metabolic pseudo-steady state over the experimental time course (Buescher *et al*, 2015). Exponential growth phase is thought to reflect metabolic pseudo-steady state as cells in culture steadily divide at their maximal condition-specific rate, provided nutrient supply does not become limiting (Buescher *et al*, 2015; Ahn & Antoniewicz, 2011). Thus, we first set out to define experimental conditions to capture exponential growth phase in normoxic and hypoxic cultures.

Cells were seeded and placed into hypoxia for 24 h prior to sample collection to provide adequate time for activation of the hypoxia-dependent transcriptional program (**Fig 1A**). We selected 0.5% oxygen for hypoxia as this level yielded the most reproducible phenotypic differences compared to 21% oxygen culture while being physiologically relevant and above the K_M_ of cytochrome *c* oxidase (electron transport chain complex IV) for oxygen (Lee *et al*, 2020; Wenger *et al*, 2015). From this starting point, we identified the optimal cell seeding density and time course to capture exponential cell growth (**Fig 1B**). LFs cultured in 0.5% oxygen grew more slowly than LFs cultured in 21% oxygen (**Fig 1C**), but slower growth was not associated with decreased cell viability (**Fig S1A**). As anticipated, hypoxic cells demonstrated robust stabilization of HIF-1α protein associated with up-regulation of downstream targets, such as glucose transporter 1 (GLUT1) and lactate dehydrogenase A (LDHA) (**Figs 1D-H**). These changes persisted for the duration of the experimental time course.

**Figure 1:**
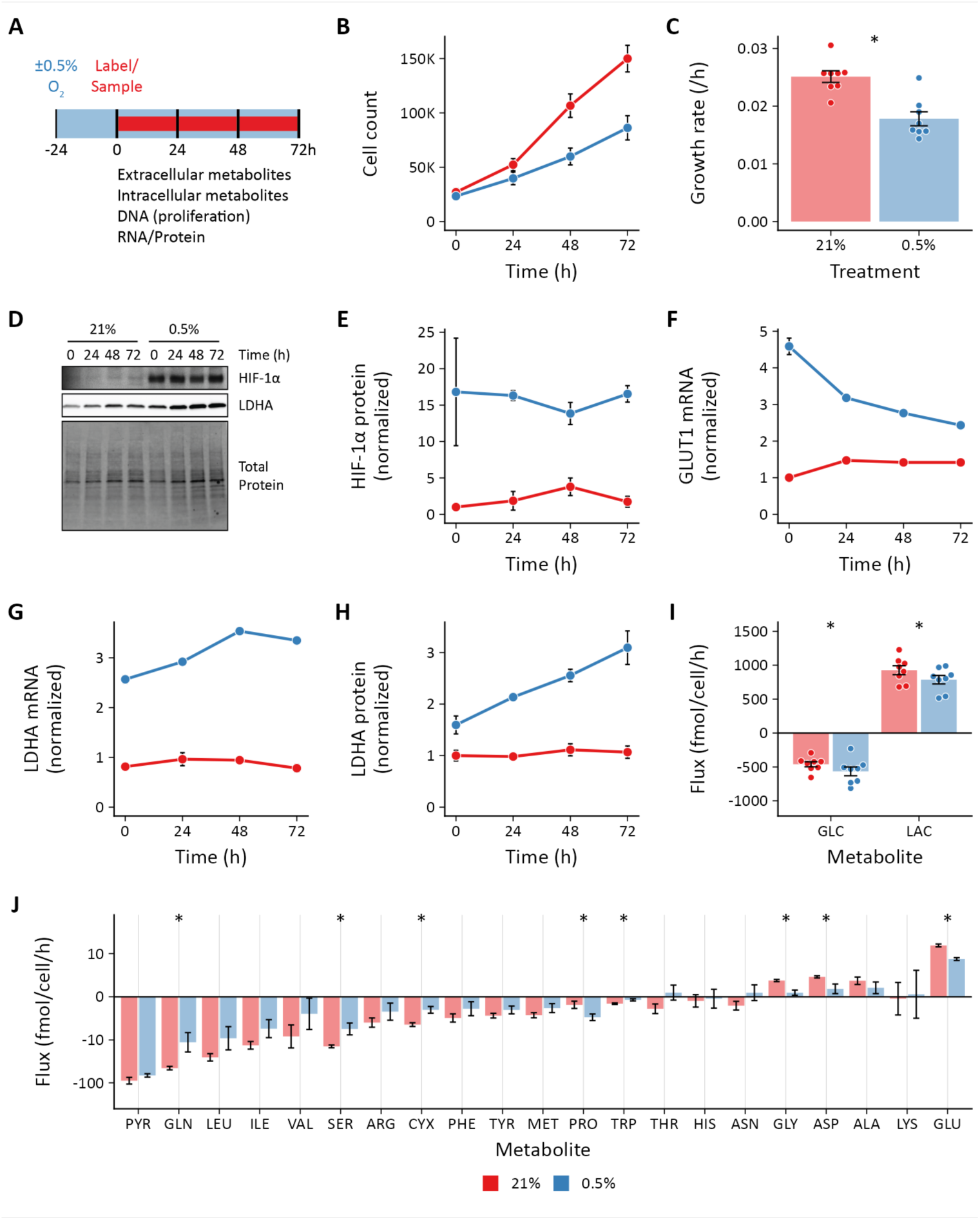
Effects of hypoxia on extracellular metabolite fluxes in lung fibroblasts. (**A**) Lung fibroblasts (LFs) were cultured in 21% or 0.5% oxygen beginning 24 h prior to time 0. Samples were collected every 24 h for 72 h. (**B**) Growth curves of LFs in each experimental condition (n = 8). (**C**) Growth rates from (B) were determined by robust linear modeling of log-transformed growth curves. (**D**) Representative immunoblot of LF protein lysates cultured as in (A). (**E**) Relative change in HIF-1α protein levels from (D) normalized to 21% oxygen at time 0 (n = 4). (**F**) Relative change in GLUT1 mRNA levels normalized to 21% oxygen treatment at time 0 (n = 4). (**G**) Relative change in LDHA mRNA levels as in (F). (**H**) Relative change in LDHA protein levels as in (E). (**I**) Extracellular fluxes of glucose (GLC) and lactate (LAC) (n = 8). By convention, negative fluxes indicate metabolite consumption. (**J**) Extracellular fluxes of pyruvate (PYR) and amino acids. Data are mean ± SEM (* p < 0.05).

Having identified experimental conditions for exponential growth, we next determined the extracellular fluxes of glucose (GLC), lactate (LAC), pyruvate (PYR), and amino acids (**Figs 1I-J**). Flux calculations incorporated changes in cell number, extracellular metabolite concentrations, metabolite degradation rates, and medium evaporation over time (Murphy & Young, 2013) (**Figure S1**). Interestingly, while we observed a modest increase in glucose uptake, we found that hypoxia actually decreased lactate efflux (**Figs 1I**). This decrease in lactate efflux occurred despite activation of the HIF-1α transcriptional program as reflected by increased expression of GLUT1 and LDHA. To test if more severe hypoxia would augment glycolysis, we cultured cells in 0.2% ambient oxygen (**Fig S2**). Under these conditions, we observed no change in glucose or lactate fluxes, similar to 0.5% oxygen culture. To test if this unexpected response was unique to LFs, we next studied PASMCs under 0.5% oxygen conditions (**Fig S3**). PASMCs grew faster than LFs, and so samples were collected every 12 h for 48 h for these cells. Again, similar to LFs at 0.5% and 0.2% oxygen, we observed no change in glucose uptake and reduced lactate efflux in PASMCs regardless of HIF-1α stabilization. Together, these data suggest that hypoxia uncouples HIF target gene expression and glycolytic flux in proliferating primary cells.

Given that hypoxia exposure did not increase glycolysis in LFs, we next wanted to determine how these cells responded to HIF-1α stabilization in normoxia. To accomplish this, LFs were treated with the PHD inhibitor molidustat (BAY, 10 μM) using a similar time course as our hypoxia experiments. Cells were treated with BAY for 24 h to activate the HIF transcriptional program prior to sample collection (**Fig 2A**). As with hypoxia, BAY decreased cell growth rate (**Figs 2B-C**) and activated the HIF-1α transcriptional program (**Figs 2D-H**). Compared to hypoxia, BAY treatment resulted in a similar activation of HIF-1α target gene transcription and protein expression. Unlike hypoxia, however, HIF-1α stabilization in normoxia markedly increased glucose uptake and lactate efflux (**Fig 2I**). This finding was consistent with the increased expression of glycolytic genes that we observed. Interestingly, although hypoxia and BAY treatments resulted in similar increases in HIF-1α, GLUT1, and LDHA, the glycolytic response was markedly different.

**Figure 2:**
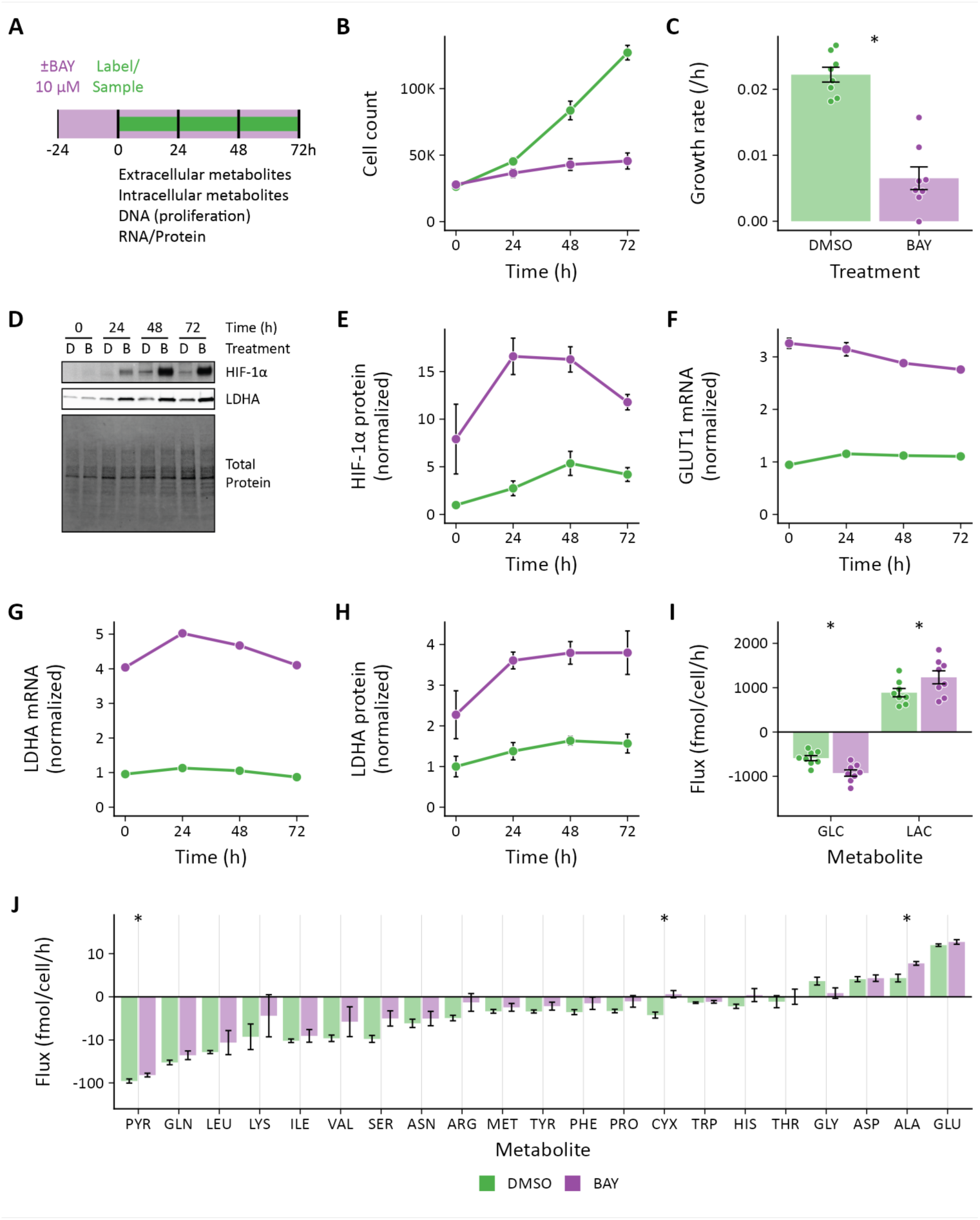
Effects of pharmacologic prolyl hydroxylase inhibition on extracellular metabolite fluxes in lung fibroblasts. (**A**) Lung fibroblasts (LFs) were treated with the prolyl hydroxlyase inhibitor molidustat (BAY, 10 μM) or DMSO beginning 24 h prior to time 0. Samples were collected every 24 h for 72 h. (**B**) Growth curves of LFs in each experimental condition (n = 8). (**C**) Growth rates from (B). (**D**) Representative immunoblot of LF protein lysates cultured as in (A). (**E**) Relative change in HIF-1α protein levels from (D) normalized to DMSO at time 0 (n = 4). (**F**) Relative change in GLUT1 mRNA levels normalized to DMSO at time 0 (n = 4). (**G**) Relative change in LDHA mRNA levels as in (F). (**H**) Relative change in LDHA protein levels as in (E). (**I**) Extracellular fluxes of glucose (GLC) and lactate (LAC) (n = 8). By convention, negative fluxes indicate metabolite consumption. (**J**) Extracellular fluxes of pyruvate (PYR) and amino acids. Data are mean ± SEM (* p < 0.05).

### Extracellular fluxes are treatment and cell-type dependent

In addition to glucose and lactate, we also determined the extracellular fluxes of pyruvate and amino acids (**Figs 1J, 2J, S2J, S3J**). To our knowledge, this is the first comprehensive extracellular flux profiling of key metabolic substrates in these primary cells. In LFs, overall, changes in these fluxes were modest, with hypoxia generally decreasing the fluxes of all measured metabolites. These findings were similar with 0.2% oxygen exposure (**Fig S2J**).

Notably, we observed a significant decrease in glutamine consumption in hypoxic LFs. This finding contrasts with previous studies of cancer cell metabolism demonstrating increased glutamine uptake as a key feature of the metabolic response to hypoxia (Metallo *et al*, 2011; Wise *et al*, 2011; Gameiro *et al*, 2013a). In these systems, glutamine-derived α-ketoglutarate was reductively carboxylated by isocitrate dehydrogenase enzymes to generate citrate for lipogenesis. In addition, glutamine has been shown to support TCA cycling in hypoxia in a Burkitt lymphoma model (Le *et al*, 2012). Unlike LFs, PASMCs did exhibit a trend toward increased glutamine uptake (**Figure S3J**), suggesting a greater reliance on these metabolic pathways in their adaptive response to hypoxia.

In LFs, among all of the measured amino acid fluxes, proline consumption uniquely increased (**Fig 1J**). Hypoxia increases collagen expression in these cells (Liu *et al*, 2013) and proline constitutes ∼ 10% of the total amino acid content of collagens. Together, these data suggest an important contribution of extracellular proline to collagen production in hypoxic LFs as has been observed in other fibroblast cell lineages (Szoka *et al*, 2017).

In PASMCs, we observed increased consumption of the branched-chain amino acids (BCAAs) leucine and valine as well as arginine (**Figure S3J**), which was not observed in LFs. BCAAs are transaminated by branch chain amino transferase enzymes to branched chain α-keto acids (BCKAs). BCKAs are further metabolized to yield acyl-CoA derivatives for lipogenesis or oxidation (Mann *et al*, 2021; Crown *et al*, 2015). Previous studies have shown that hypoxia up-regulates arginase expression in hypoxic PASMCs (Chen *et al*, 2009; Xue *et al*, 2017) to support polyamine and proline synthesis required for cell proliferation (Li *et al*, 2001). Interestingly, activation of these metabolic pathways in hypoxia was not observed in LFs and suggests distinct metabolic vulnerabilities of these different cell types.

Compared to hypoxia treatment, BAY demonstrated more modest effects on amino acid fluxes generally (**Figure 2J**). In particular, glutamate efflux was not affected by BAY treatment, while it was reduced by hypoxia. Alanine efflux was increased by BAY treatment, but decreased by hypoxia. In addition to the glucose and lactate fluxes noted above, these findings further highlight fundamental differences in the metabolic consequences of HIF activation in normoxia and hypoxia.

### Isotope tracing reveals altered substrate utilization in hypoxia

To investigate intracellular metabolic reprogramming in hypoxic cells, we performed ^13^C stable isotope tracing with [U-^13^C_6_]-glucose, [1,2-^13^C_2_]-glucose, and [U-^13^C_5_]-glutamine. Isotopic enrichment of downstream metabolites in glycolysis and the TCA cycle were determined by LC-MS (**Figs S3, S4**). Overall, relatively small changes in the patterns of isotope incorporation were observed following hypoxia or BAY treatment. The most substantial differences were observed in pyruvate (PYR), the terminal product of glycolysis, and citrate (CIT), a central metabolic node in TCA and fatty acid metabolism (**Figs 3A-C**). Both hypoxia and BAY treatments decreased incorporation of glucose-derived carbon into pyruvate (**Fig 3A**) (*i.e.,* the unlabeled, or M0, fraction was greater). This suggests a greater contribution from an unlabeled carbon source, such as extracellular pyruvate, lactate, or alanine, than from glucose to the intracellular pyruvate pool following PHD inhibition.

**Figure 3:**
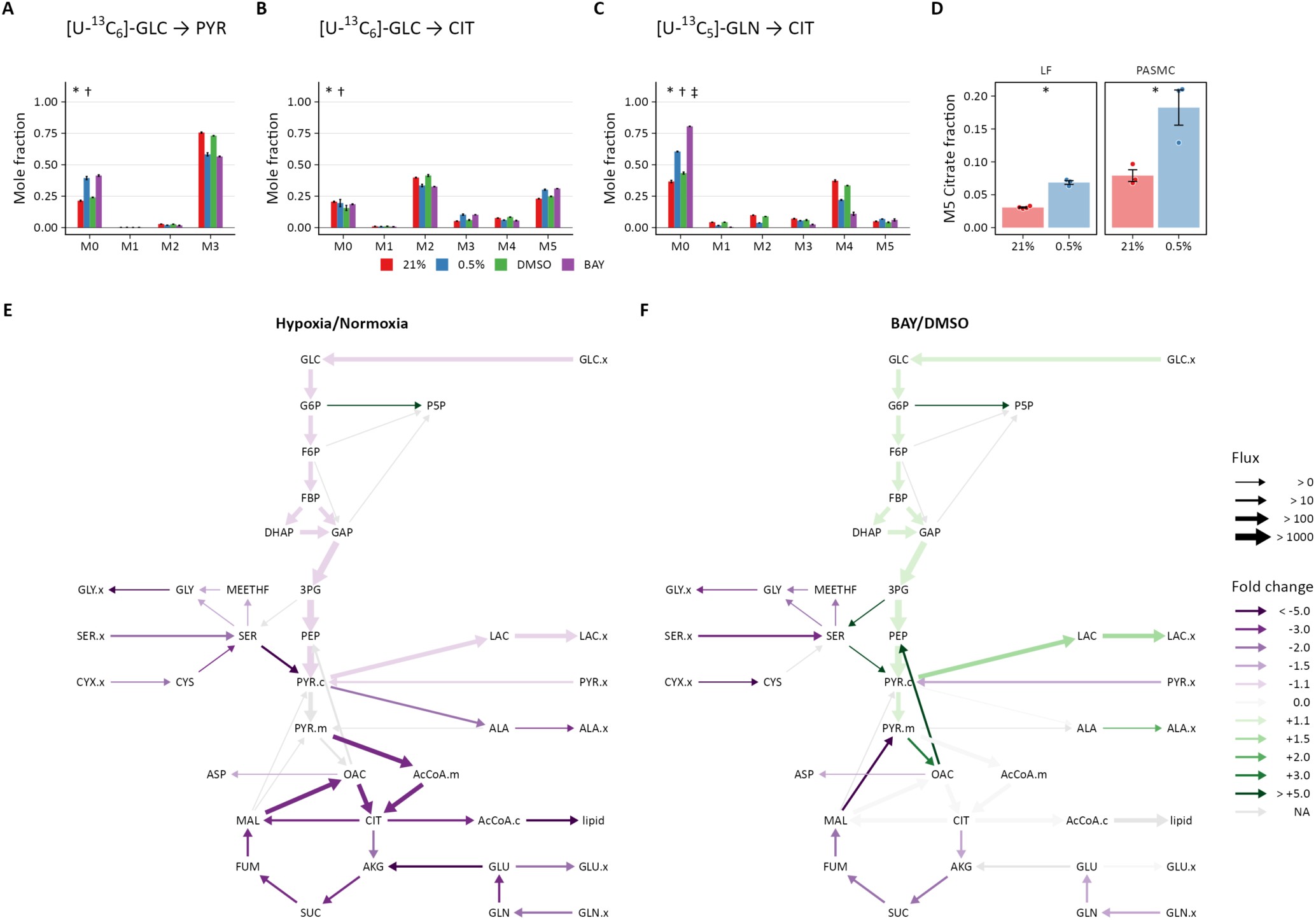
Metabolic flux analysis of lung fibroblasts following hypoxic and pharmacologic PHD inhibition. (**A**) Mass isotopomer distribution (MID) of pyruvate (PYR) following 72 h labeling with [U-^13^C_6_]-glucose (GLC). (**B**) MID of citrate after 72 h labeling with [U-^13^C_6_]- GLC (**C**) MID of citrate after 72 h labeling with [U-^13^C_5_]-glutamine (GLN). Data are mean ± SEM (n = 4, p < 0.05 indicated as * 0.5% *v.* 21% oxygen, † BAY *v.* DMSO, ‡ Δoxygen *v.* ΔBAY). (**D**) Fraction of M5 citrate indicating reductive carboxylation after labeling with [U-^13^C_5_]-GLN in LFs and PASMCs (n = 3-4, * p < 0.05). (**E**) Ratio of modeled metabolic fluxes in 0.5% oxygen compared to 21% oxygen. Fluxes with non-overlapping confidence intervals are highlighted with arrows colored according to the magnitude of the fold change. Arrow thickness corresponds to the absolute flux measured in hypoxia. (**F**) Ratio of metabolic fluxes in BAY-treated cells compared to DMSO-treated control. Arrows are colored as in (E) and arrow weights correspond to the absolute flux as measured in BAY-treated cells.

Total citrate labeling from [U-^13^C_6_]-glucose was unchanged across the treatment conditions (**Fig 3B**). As expected, we observed decreased M2 and M4 citrate isotopes, consistent with decreased pyruvate dehydrogenase activity in hypoxia. Interestingly, we observed increased M3 and M5 citrate isotopes. Pyruvate carboxylase catalyzes the carboxylation of pyruvate to oxaloacetate after which all three pyruvate carbons are incorporated into citrate by citrate synthase. Thus, this labeling pattern suggests a more prominent contribution of pyruvate carboxylase to sustain TCA cycle anaplerosis despite pyruvate dehydrogenase inhibition following HIF-1α activation. By contrast to glucose labeling, much less citrate was labeled by glutamine with hypoxia or BAY with a more pronounced effect of BAY treatment (**Fig 3C**), suggesting a less important contribution of glutamine to TCA anaplerosis under these conditions. In addition, the overall fraction of M5 citrate resulting from reductive carboxylation of glutamine-derived α-ketoglutarate was low (< 7%) (**Fig 3D**). Although a hypoxia-mediated increase in M5 citrate was observed, the overall fraction was much less than the 10-20% levels previously reported in cancer cells (Wise *et al*, 2011; Metallo *et al*, 2011).

The stable isotope labeling patterns in PASMCs were generally similar to LFs (**Figure S5**). The most notable differences between LF and PASMC labeling were observed in citrate. Compared with LFs, a much lower fraction of total citrate was labeled by glucose in PASMCs. Less activity of pyruvate carboxylase in these cells was suggested by decreased M3 and M5 citrate isotopes after glucose labeling. Interestingly, the M5 citrate fraction in PASMCs was more consistent with previous reports from the cancer literature (**Fig 3D**), suggesting a more important role for glutamine metabolism for biomass synthesis in these cells.

### Glycolytic flux in hypoxia is closely coupled to cell growth rate

The mass isotopomer distribution for a given metabolite is determined by the complex relationship among the rate of isotope incorporation into the metabolic network, the contributions of unlabeled substrates, and fluxes through related pathways. To clarify how these labeling patterns reflect changes in intracellular metabolite fluxes, we next generated metabolic flux models incorporating the extracellular flux measurements and stable isotope tracing data described above. Preliminary labeling time courses indicated that, even after 72 h of labeling, intracellular metabolites did not reach isotopic steady state (**Fig S6**). Thus, we performed isotopically non-stationary metabolic flux analysis as implemented by Isotopomer Network Compartment Analysis (INCA) (Murphy & Young, 2013; Young *et al*, 2014; Jazmin & Young, 2013) (**Figs 3E-F, S7, Tables S1-S3**).

Overall, LF and PASMC metabolic fluxes were dominated by high rates of glucose uptake and glycolysis (**Figs S7A-B**). Approximately 10% of cytoplasmic pyruvate enters the TCA cycle with the balance converted to lactate. Consistent with extracellular flux measurements and isotope labeling patterns described above, significant reductions in glycolysis, the TCA cycle, and amino acid metabolism were observed in the metabolic flux models of LFs cultured in hypoxia (**Fig 3E**). A significant increase in pentose phosphate pathway flux was also observed, although the absolute flux through this pathway is low. By contrast, HIF-1α activation by BAY in 21% oxygen increased glycolysis and lactate fermentation by nearly 50% (**Figure 3F**), but had a similar effect on decreasing serine and glutamine uptake as hypoxia. Metabolite fluxes in DMSO-treated cells were similar to 21% oxygen controls (**Table S1-S2**).

In normoxia, the magnitude of intracellular metabolite fluxes was generally similar in LFs and PASMCs (**Figs S7A-C, Tables S1, S3**). Compared to LFs, PASMCs had slower rates of glycolysis and faster rates of TCA metabolism driven, in part, by increased glutamine uptake. In hypoxia, PASMCs exhibited similar decreases in glycolytic flux as LFs but also a marked, and unexpected, increase in TCA flux (**Figure S7D**). The increased TCA flux in PASMCs was driven by increased glutamine consumption. This finding is similar to a prior report of glutamine-driven oxidative phosphorylation in hypoxic cancer cells (Fan *et al*, 2013), where oxidative phosphorylation continued to provide the majority of cellular ATP even at 1% oxygen.

Given the global decrease in bioenergetic metabolic flux in hypoxic LFs, we hypothesized that these differences may be a consequence of decreased growth rate. After normalizing metabolite fluxes in normoxia and hypoxia to the cell growth rate, a modest increase (∼10%) in glycolytic flux was observed (**Fig S7E**). This finding suggests that, while glycolysis increases relative to growth rate in hypoxic cells, regulators of cell proliferation rate override the consequences of the HIF-1α transcriptional program. Indeed, even after adjusting for cell growth rate, the relative increase in glycolytic flux is modest compared to the marked up-regulation of glycolytic protein levels and the glycolytic potential of these cells demonstrated by BAY treatment in normoxia. BAY treatment decreased cell proliferation rate (**Fig 2B**), indicating that, unlike hypoxia, BAY treatment in normoxia uncouples cell proliferation and metabolic flux.

### Hypoxia and BAY treatment increase lactate oxidation

Although the metabolite exchange fluxes for bidirectional reactions tend to be poorly resolved by metabolic flux analysis (Wiechert, 2007), two observations are worth highlighting (**Tables S1-S3**). First, consistent with the stable isotope tracing results, the modeled rate of reductive carboxylation through reverse flux by isocitrate dehydrogenase in LFs is low (∼4 fmol/cell/h), unchanged by hypoxia, and modestly increased by BAY treatment. By contrast, the rate of reductive carboxylation increases 6-fold in PASMCs in hypoxia, highlighting an important role for this pathway in the metabolic response of PASMCs to decreased oxygen availability (**Fig 4A**).

**Figure 4:**
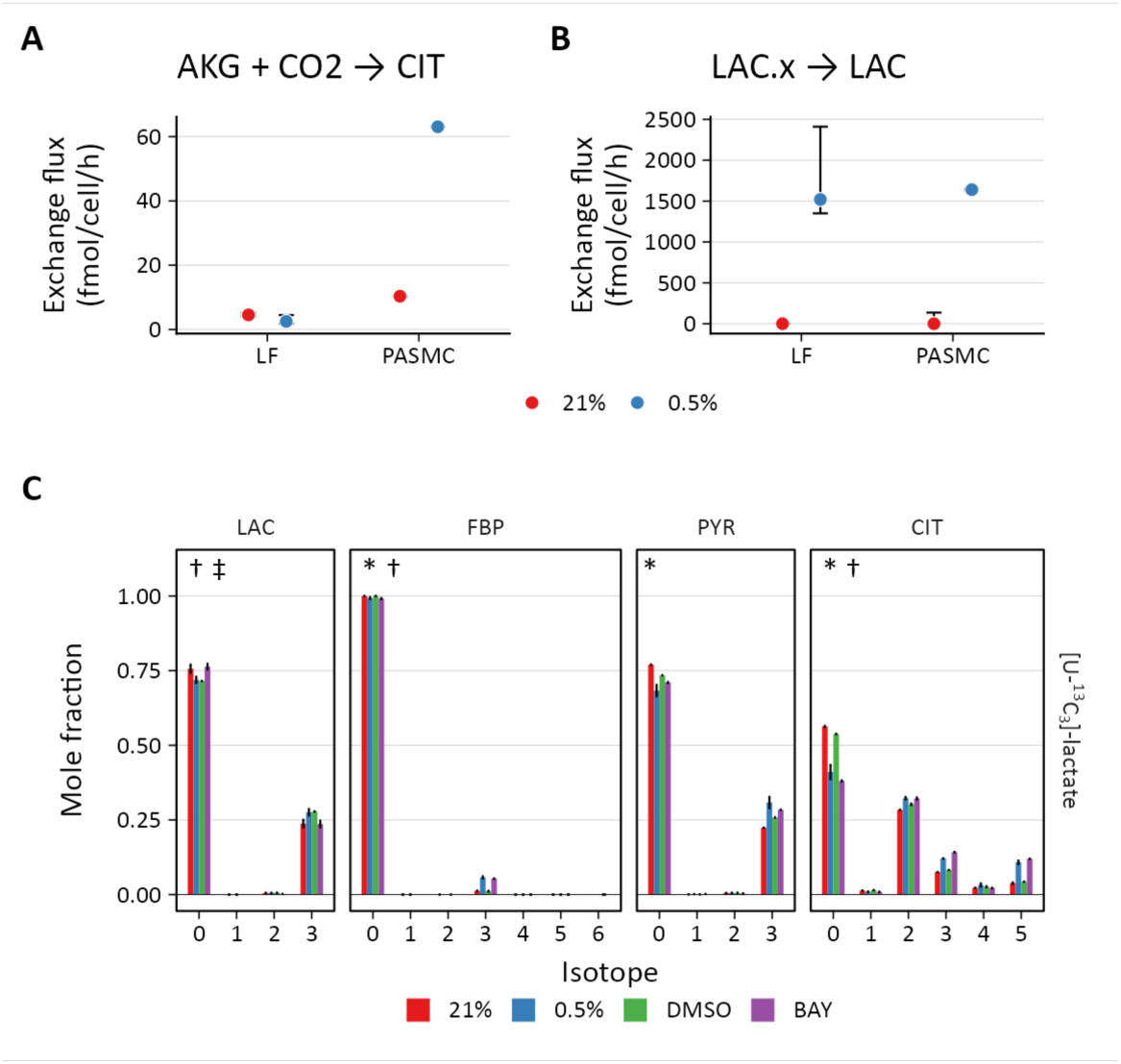
PHD inhibition increases lactate uptake and oxidation. (**A**) Exchange flux estimates for reductive carboxylation of α-ketoglutarate (AKG) to citrate (CIT). Data show the estimate with upper and lower bounds for LFs and PASMCs. (**B**) Exchange flux estimates for lactate import presented as in (A). (**C**) Mass isotopomer distributions of lactate (LAC), fructose-bisphosphate (FBP), pyruvate (PYR), and citrate (CIT) in LFs following 72 h labeling with [U-^13^C_3_] lactate. Data are mean ± SEM (n = 4, p < 0.05 indicated as * 0.5% *v.* 21% oxygen, † BAY *v.* DMSO, ‡ Δoxygen *v.* ΔBAY).

Second, PHD inhibition is associated with a marked increase in the lactate transport exchange flux in LFs from ∼ 0 to 1,500 and 700 fmol/cell/h in 0.5% oxygen and BAY treatment conditions, respectively, with similar results in PASMCs (**Fig 4B**). Since the net lactate transport flux is secretion, this observation suggests increased lactate uptake with hypoxia or BAY treatment, a finding that may be consistent with the HIF-driven increased expression of the reversible lactate transporter MCT4 (Contreras-Baeza *et al*, 2019). To investigate this hypothesis, LFs and PASMCs were treated with [U-^13^C_3_] lactate (2 mM) and ^13^C incorporation into intracellular metabolites was analyzed by LC-MS (**Figs 4, S4, S5**). Lactate labeled ∼50% of citrate and ∼20% of downstream TCA cycle metabolites in both LFs and PASMCs, indicating that lactate may be an important respiratory fuel source in these cells even though lactate efflux is high. Although lactate has been used less commonly than glucose and glutamine in stable isotope tracing studies, Faubert and colleagues (2017) demonstrated lactate incorporation in human lung adenocarcinoma *in vivo*. In this study, lactate incorporation corresponded to regions of high glucose uptake as determined by [¹⁸F]-fluorodeoxyglucose positron emission tomography, suggesting that lactate consumption can occur even in areas of high glucose utilization. Subsequently, investigators have demonstrated the importance of lactate as a metabolic fuel *in vivo* (Hui *et al*, 2017; Hui *et al*, 2020). As predicated from our metabolic flux analysis, with hypoxia or BAY treatment, we observed increased labeling of the TCA metabolites citrate (CIT), α-ketoglutarate (AKG), malate (MAL), and aspartate (ASP) in LFs. Interestingly, although increased labeling of pyruvate was observed in hypoxic PASMCs, the label was not incorporated into the TCA cycle as observed in LFs (**Fig S5**).

In addition to downstream metabolites, we also observed hypoxia- and BAY-dependent increases in lactate incorporation in fructose bisphosphate (FBP) and 3-phosphoglycerate (3PG). This observation is consistent with prior reports describing hypoxia-mediated increases in gluconeogenesis and glycogen synthesis (Owczarek *et al*, 2020; Pelletier *et al*, 2012; Favaro *et al*, 2012). These data suggest that lactate also makes a small (∼5% carbon) contribution to glycogen precursors. Together, these findings from stable isotope tracing of lactate reveal its important contribution to primary cell metabolism under standard culture conditions, but also reveal increased utilization of this substrate in hypoxia.

### Hypoxia abrogates the effects of BAY on increasing glycolysis

To reconcile the differential effects of prolyl hydroxylase inhibition by hypoxia and BAY, we next addressed whether hypoxia could suppress the effects of BAY on glucose and lactate fluxes (**Figs 5A-C**). LFs cultured in standard growth medium were treated with BAY and placed in either 21% or 0.5% oxygen. Similar to previous experiments, BAY treatment decreased cell growth rate, increased glucose uptake, and increased lactate efflux in 21% oxygen. However, when combined with 0.5% oxygen, BAY treatment was unable to enhance lactate efflux. These data clearly demonstrate that hypoxia antagonizes the effects of HIF-1α activation on glycolytic flux in these cells.

**Figure 5:**
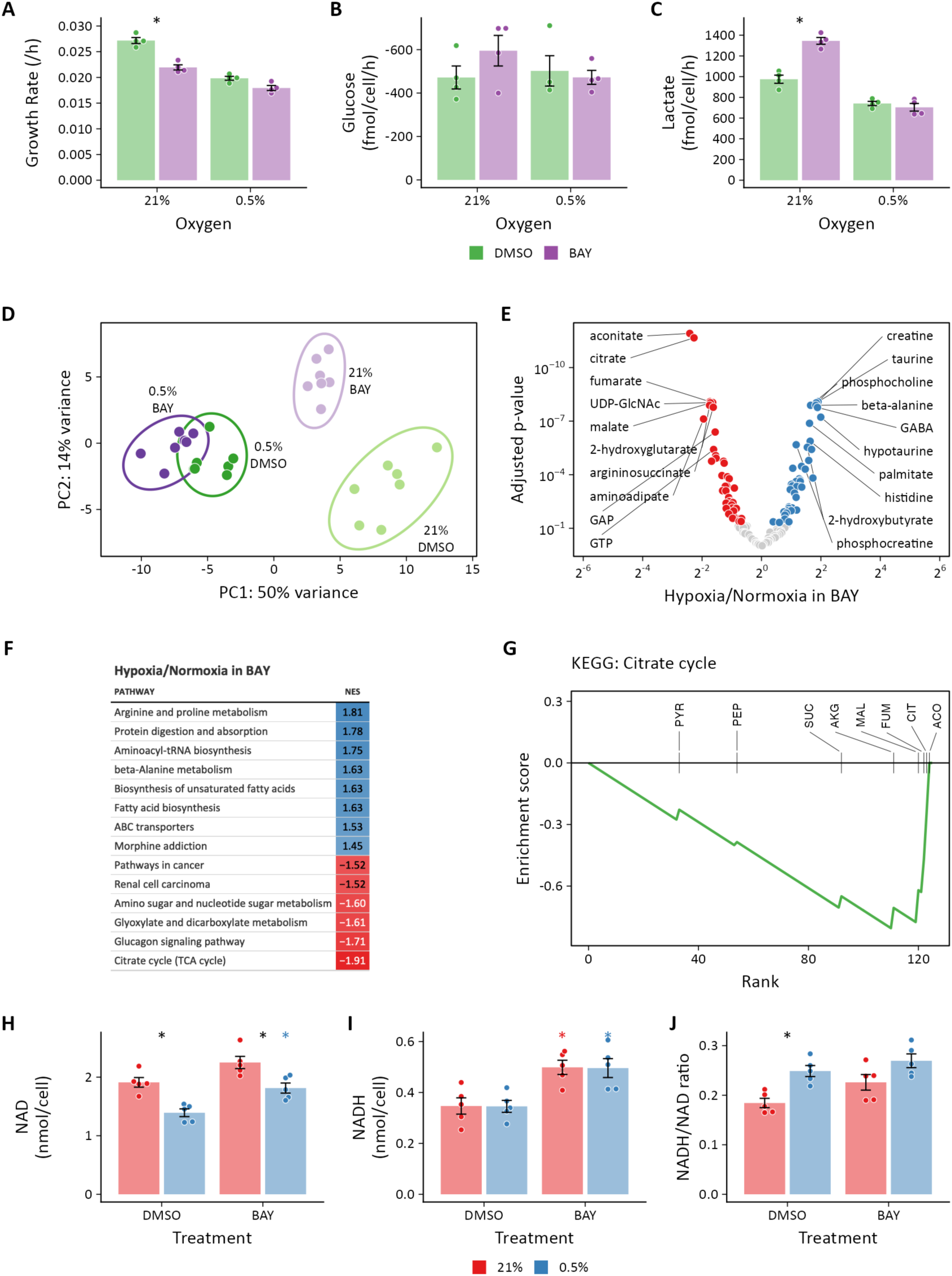
Metabolomic analysis of molidustat treatment in normoxia and hypoxia. (**A**) LFs were cultured in standard growth medium and treated with molidustat (BAY, 10 μM) or vehicle (DMSO, 0.1%) in 21% or 0.5% oxygen conditions for 72 h. Growth rates were determined by linear fitting of log-transformed growth curves. (**B**) Glucose uptake in LFs treated with BAY cultured in hypoxia (note reversed *y*-axis). (**C**) Lactate efflux in LFs treated with BAY cultured in hypoxia. Data are mean ± SEM (* p < 0.05 BAY *v.* DMSO within a given oxygen exposure). (**D**) Principal component analysis of intracellular metabolites following 72 h of treatment as in (A) suggests a dominant effect of hypoxia over pharmacologic PHD inhibition on the metabolome (n = 7). (**E**) Volcano plot of intracellular metabolites of BAY-treated cells cultured in 21% or 0.5% oxygen. Circles indicate significantly increased (*blue*) and decreased (*red*) levels in hypoxia. (**F**) Metabolite set enrichment analysis of metabolites from (E). All KEGG pathways with p-values < 0.05 are shown. (**G**) Leading edge analysis of the TCA cycle metabolite set. Negative values indicate the relative enrichment associated with BAY treatment compared to hypoxia treatment. Abbreviations: PYR, pyruvate; SUC, succinate; PEP, phosphoenolpyruvate; CIT, citrate; AKG, α-ketoglutarate; MAL, malate; ACO, aconitate; FUM, fumarate. (**H**) NAD^+^ from LFs cultured as in (A). (**I**) NADH from LFs cultured as in (A). (**J**) NADH/NAD^+^ ratio from (H) and (I). NAD and NADH were determined by enzymatic assay. Data are mean ± SEM (n = 5, p < 0.05 with *black* * indicating a significant effect of treatment within a given oxygen tension and *colored* * indicating a significant effect of treatment within a given oxygen tension denoted by the color).

To investigate these metabolic differences further, we performed metabolomic profiling of LFs treated for 72 h with hypoxia or BAY separately or in combination. Both 0.5% oxygen and BAY treatment induced marked changes in intracellular metabolite levels (**Fig S8**). Of 133 total metabolites, 99 were differentially regulated by hypoxia and 54 were differentially regulated by BAY. Of the differentially regulated metabolites, 44 were affected by both 0.5% oxygen and BAY treatment. Metabolite set enrichment analysis of KEGG biochemical pathways identified increased enrichment of arginine and proline metabolism and fatty acid biosynthesis pathways with hypoxia (**Fig S8D**). By contrast, BAY treatment was enriched for metabolites involved in pentose/glucuronate interconversions and glycolysis (**Fig S8E**). Aspartate was the most significantly decreased metabolite with both treatments, consistent with prior reports demonstrating an important role for HIF-1 regulation of aspartate biosynthesis in cancer cells (Garcia-Bermudez *et al*, 2018; Melendez-Rodriguez *et al*, 2019).

Principal component analysis revealed greater similarity among both treatment groups cultured in 0.5% oxygen than among the BAY-treatment groups (**Fig 5D**). Moreover, these hypoxia-treated cells were well-segregated from cells treated with BAY alone. These observations are, again, consistent with the results of the metabolic flux models demonstrating an overriding effect of hypoxia *per se* on cell metabolism and highlighting important differences between hypoxic and pharmacologic PHD inhibition. To identify the metabolic changes that depend on hypoxia rather than BAY inhibition, we identified differentially regulated metabolites in BAY treated cells cultured in normoxia and hypoxia (**Fig 5E**). Of 133 metabolites, 83 were significantly differentially regulated by hypoxia in BAY-treated cells. An enrichment analysis of these differentially regulated metabolites demonstrated up-regulation of arginine and proline metabolism and down-regulation of the TCA cycle as the most impacted by hypoxia in BAY treated cells (**Fig 5F**). Indeed, leading edge analysis highlights negative enrichment scores associated with all of the TCA metabolites detected by our platform (**Fig 5G**). This result indicates better preservation of TCA cycle flux in normoxic BAY-treated cells than in hypoxic cells, as suggested by our metabolic flux models where hypoxia resulted in a 1.5-2-fold reduction of TCA flux compared to a 1.1-1.5-fold reduction with BAY treatment (**Fig 3**).

In addition to these differential effects on polar metabolite levels, we reasoned that another critical difference between hypoxia and BAY treatment is the impact of hypoxia on cellular redox state. As oxygen deprivation causes reductive stress (Xiao & Loscalzo, 2020), we next measured the impact of these treatments on intracellular NAD(H) (**Figs 5H-J**). As expected, hypoxia increased the NADH/NAD^+^ ratio, driven primarily by a decrease in intracellular NAD^+^. Interestingly, while BAY treatment increased the levels of NADH, a concomitant increase in NAD^+^ resulted in preservation of the NADH/NAD^+^ ratio. As NADH accumulation is a putative inhibitor of glycolytic flux (Tilton *et al*, 1991), this may be one mechanism by which glycolytic flux is decreased in hypoxia but not following BAY treatment.

### Transcriptomic analysis identifies regulators of metabolism in hypoxia

To identify the upstream regulators of the observed metabolic changes, we next performed RNA-seq transcriptomic analysis of LFs treated with hypoxia or BAY, separately or together (**Fig 6, S9**). As anticipated, both hypoxia and BAY treatment induced substantial changes in gene expression (**Fig S9A, S9B**). Of the 9,923 differentially expressed genes across both conditions, 869 (9%) were unique to BAY treatment in normoxia, 4,002 (40%) were shared by BAY and hypoxia, while 5,052 (51%) were unique to 0.5% hypoxia culture (**Fig S9C**). This distribution of transcriptional changes was nearly identical to that observed for the metabolite changes (**Fig S8C**), where half of the hypoxia-mediated changes overlapped with BAY treatment and half were unique to hypoxia. Gene set enrichment analysis of these differentially regulated metabolites was performed using Molecular Signatures Database “Hallmark” gene sets (Liberzon *et al*, 2015) (**Figs S9D-F**). As expected, both treatments were associated with enrichment of the “hypoxia” and “glycolysis” gene sets.

**Figure 6:**
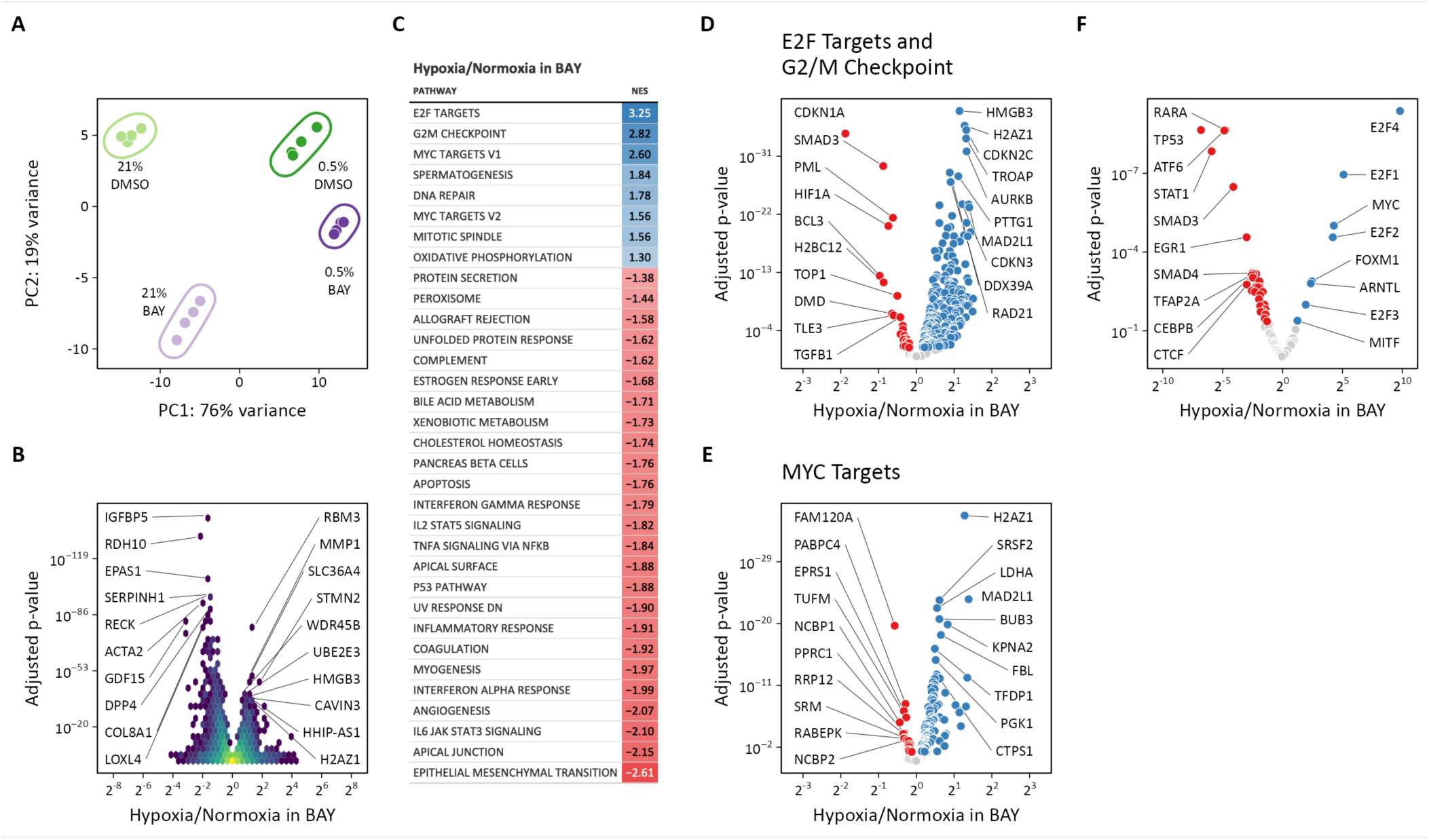
Transcriptomic analysis of molidustat treatment in normoxia and hypoxia. (**A**) Principal component analysis of transcriptional changes in lung fibroblasts following 72 h of treatment with 0.5% oxygen or molidustat (BAY), separately or together (n = 4). (**B**) Volcano plot illustrating the effects of hypoxia in BAY-treated cells on gene expression. (**C**) Gene set enrichment analysis of transcripts from (B). (**D**) Volcano plot of those transcripts comprising the E2F Targets and G2/M Checkpoint Hallmark gene sets. (**E**) Volcano plot of those transcripts comprising the MYC Targets V1 and V2 Hallmark gene sets. (**F**) Volcano plot illustrating the results of a transcription factor enrichment analysis suggests mechanisms for differential regulation of gene expression following hypoxia or BAY treatment.

Given the disparate effects of hypoxic and pharmacologic PHD inhibition on cellular metabolism described above, we focused our transcriptomics analyses on the differences between hypoxia and BAY treatments. Principal component analysis again demonstrated clear separation among the four treatment groups (**Fig 6A**). The first and second principal components correspond to 0.5% oxygen and BAY treatments, respectively. Consistent with our prior observations, the combination of 0.5% oxygen plus BAY was more similar to 0.5% oxygen alone with decreased distance between both hypoxia-treated groups along the axis of the second principal component, again, consistent with the hypothesis that hypoxia overrides the effects of BAY treatment. To identify the transcripts driving these differences, we identified genes differentially expressed following hypoxia in BAY-treated cells (**Fig 6B**). Interestingly, an enrichment analysis of these differentially expressed transcripts identified pro-proliferative gene sets like “E2F targets”, “G2/M checkpoint”, and “MYC targets” associated with hypoxia (**Fig 6C-E**). These findings were further supported by a transcription factor enrichment analysis identifying enrichment of MYC transcription factor activity associated with hypoxia, but not BAY treatment (**Fig 6F**). Classically, hypoxia and HIF activation are thought to inhibit cell proliferation by inhibiting pro-proliferative MYC signaling (Koshiji *et al*, 2004). These results indicate that hypoxia-induced MYC activation may be sustaining proliferation in these LFs. We reasoned that these pro-proliferative signals may also account for the unexpected effects of hypoxia on glycolysis that we observed.

### MYC antagonizes HIF-dependent glycolytic fluxes

To test the role of hypoxia-induced MYC activation in the metabolic response to hypoxia in proliferating primary cells, we first measured MYC protein levels by immunoblot. Consistent with our bioinformatic results, we observed increased MYC protein levels in hypoxia-treated cells, but not with BAY-treatment alone, where MYC was decreased (**Fig 7A-B**). Together, these data suggest a model whereby hypoxia activates MYC and inhibits HIF-driven increases in glycolysis. Inhibiting MYC in hypoxia, therefore, should increase glycolysis. Conversely, HIF-dependent glycolysis after BAY treatment in normoxia should be sensitive to inhibition by MYC overexpression (**Fig 7C**). To test the hypothesis that hypoxia-induced MYC expression inhibits glycolysis in primary cells, we first combined MYC knockdown with hypoxia treatment (**Fig 7D-F**). As expected, MYC-deficient cells proliferated more slowly in normoxia and MYC was essential for sustaining cell proliferation in hypoxia (**Fig 7E**). Consistent with our hypothesis, MYC-knockdown cells demonstrated increased lactate efflux with hypoxia culture, unlike control siRNA-treated cells (**Fig 7F**). We next performed the complementary experiment to determine whether MYC overexpression could attenuate the increase in glycolysis observed with BAY treatment (**Fig 7G-I**). MYC increased the proliferation rate of DMSO-treated cells, although it did not augment the proliferation rate of BAY-treated cells (**Fig 7H**). As expected, MYC overexpression blocked the BAY-stimulated increase in lactate efflux. Together, these data suggest that hypoxia-induced MYC expression may be one factor that uncouples the HIF transcriptional program from glycolytic flux in proliferating primary cells.

**Figure 7:**
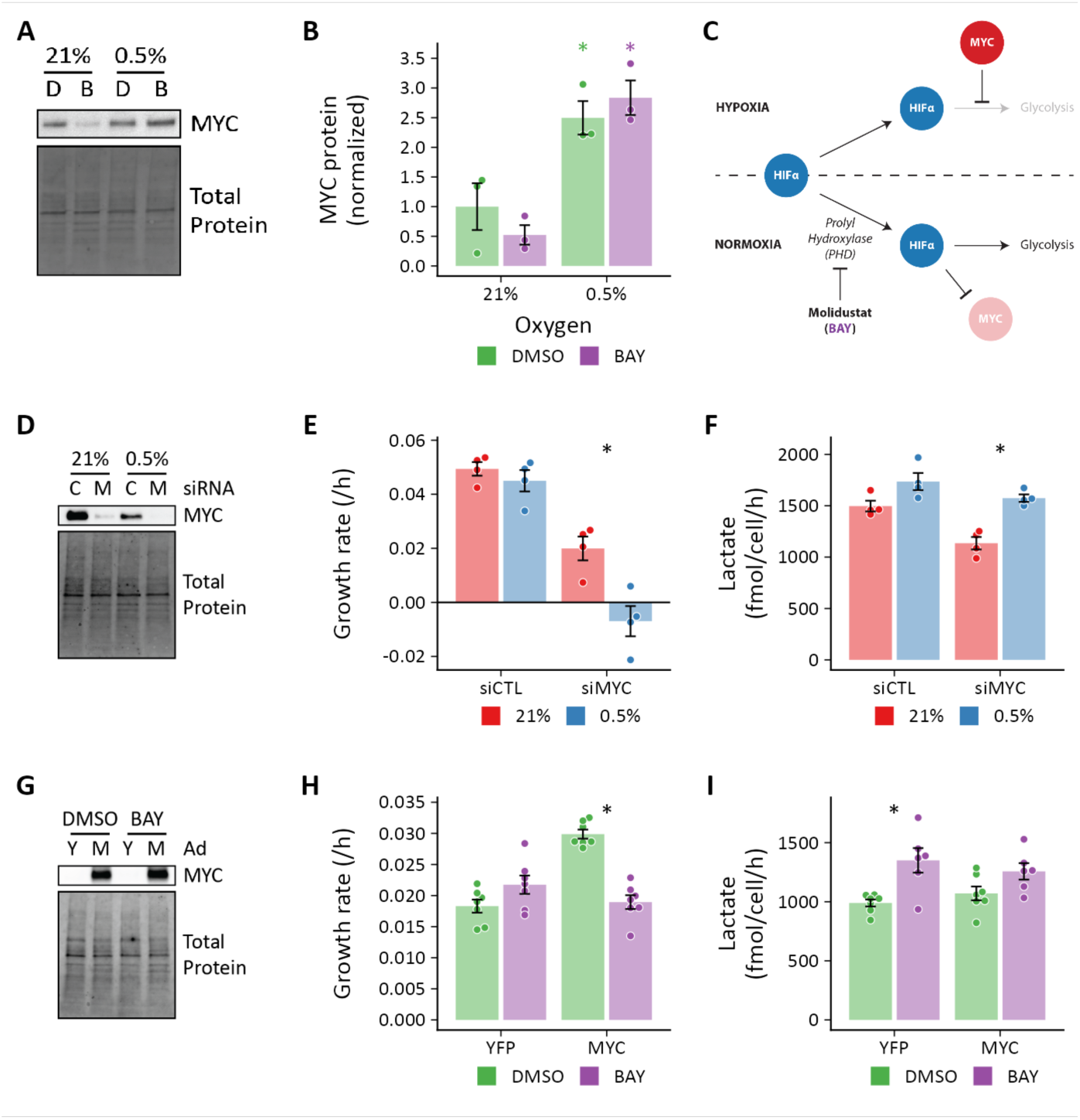
MYC regulates HIF-dependent glycolytic flux. (**A**) Representative immunoblot of MYC protein expression in lung fibroblasts (LFs) following 72 h of treatment with 0.5% oxygen or molidustat (BAY, B). (**B**) Quantification of band densities from (A). (**C**) Working model illustrating that hypoxia-induced MYC antagonizes HIF-dependent metabolic and transcriptional events. (**D**) Representative immunoblot of LFs treated with siRNA targeting MYC (M) demonstrating adequate protein knockdown. (**E**) Growth rates of MYC-knockdown cells cultured in hypoxia. (**F**) Lactate efflux rates of MYC-knockdown cells cultured in hypoxia. (**G**) Representative immunoblot of LFs treated with MYC adenovirus. (**H**) Growth rates of MYC overexpressing cells cultured with BAY. (**I**) Lactate efflux rates of MYC overexpressing cells cultured with BAY. Data are mean ± SEM (n = 3-7, p < 0.05 as indicated by black * for differences within groups and colored * for differences between groups defined by the *x*-axis).

## DISCUSSION

In this work, we used ^13^C metabolic flux analysis to identify hypoxia-mediated metabolic changes in proliferating human primary cells. Our principal finding was that hypoxia reduced, rather than increased, carbon flux through glycolysis and lactate fermentation pathways despite robust activation of the HIF transcriptional program and up-regulation of glycolytic genes. Certainly, the LFs studied here are capable of augmenting glycolysis in response to HIF stabilization, as demonstrated by experiments with the PHD inhibitor BAY; however, these effects are completely attenuated when BAY-treated cells are cultured in hypoxia. Together, these findings suggest that changes in enzyme levels alone are insufficient to alter metabolic flux in hypoxia and point to the importance of regulatory mechanisms that supersede the effects of HIF-dependent gene transcription in primary cells.

Our data indicate that hypoxia-induced MYC expression is one such regulatory mechanism. MYC is a transcription factor that regulates the expression of numerous genes involved in many biological processes, including metabolism, proliferation, apoptosis, and differentiation (Dang *et al*, 2006; Li *et al*, 2020; Stine *et al*, 2015). As deregulated MYC activity has been associated with the majority of human cancers (Vita & Henriksson, 2006), much of our understanding of MYC regulation comes from studies using cancer cell models (Li *et al*, 2020; Stine *et al*, 2015; Dang, 2012; Madden *et al*, 2021), while the role of MYC in the biology of untransformed cells is less well understood. The literature describes a complex and reciprocal relationship between HIF and MYC that depends on both environmental (*e.g.*, hypoxia) and cellular context (Li *et al*, 2020). Generally, HIF-1 has been observed to inhibit MYC through multiple mechanisms (Koshiji *et al*, 2004, 2005; Gordan *et al*, 2007; Zhang *et al*, 2007), and this previous work is consistent with our present observations that HIF stabilization following BAY treatment in normoxia decreased MYC protein and target gene expression. Conversely, MYC has been implicated in increased HIF activity through transcriptional and post-transcriptional mechanisms (Zhang *et al*, 2009; Chen *et al*, 2013; Doe *et al*, 2012), primary in the context of malignant transformation. The observation that MYC may antagonize the transcriptional effects of HIF to sustain primarily cell proliferation and metabolism in hypoxia suggests a substantially different regulatory relationship than has been previously described.

Understanding how MYC transcriptional activity affects hypoxic primary cell metabolism is imperative to our understanding of cellular adaptation to hypoxia. MYC stimulates the expression of nuclear-encoded mitochondrial genes and promotes mitochondrial biogenesis, both directly and through activation of mitochondrial transcriptional factor A (TFAM) (Li *et al*, 2005). Indeed, we found that the oxidative phosphorylation gene set was relatively enriched with hypoxia culture of BAY-treated cells (**Fig 6C**). In this way, hypoxic MYC activation may sustain energy production by oxidative phosphorylation, thereby decreasing the energetic demands that would otherwise drive increased glycolytic flux. Beyond oxidative phosphorylation, MYC regulates genes involved in many other intermediary metabolic pathways, including amino acids, nucleotides, and lipids (Stine *et al*, 2015), that may also impact the central pathways of carbon metabolism studied in this work.

Beyond MYC, the identification of other HIF-independent mechanisms regulating primary cell adaption to hypoxia is of critical importance. Cells express several oxygen-dependent enzymes in addition to PHD whose activities may be altered in hypoxia but not by PHD inhibition. For example, PHD is one of many α-ketoglutarate-dependent dioxygenase enzymes that rely on molecular oxygen for their catalytic activity (Islam *et al*, 2018). Jumonji-C (JmjC) domain-containing histone demethylases are other prominent members of this family whose inhibition by hypoxia has been shown to cause rapid and HIF-independent induction of histone methylation (Batie *et al*, 2019). Similarly, a recently described cysteamine dioxygenase has been shown to mediate the oxygen-dependent degradation of Regulators of G protein Signaling 4 and 5 and IL-32 (Masson *et al*, 2019). In addition to dioxygenase enzymes, electron transport chain dysfunction resulting from impaired Complex IV activity leads to increased ROS production in hypoxia (Chandel *et al*, 1998). Mitochondrial ROS increase the half-lives of several mRNAs in hypoxia, including MYC, independent of HIF stabilization (Guzy *et al*, 2005). Finally, hypoxia imposes a reductive stress on cells associated with an increase in the NADH/NAD^+^ ratio secondary to impaired electron transport (**Fig 5K**)(Chance & Williams, 1955; Garofalo *et al*, 1988). NADH accumulation may slow glycolysis *via* feedback inhibition of GAPDH (Tilton *et al*, 1991). Any or all of these molecular mechanisms may also contribute to uncoupling glycolytic enzyme expression from glycolytic flux as observed in the experiments described here.

In addition to its effects on cellular metabolism, another canonical role of HIF-1 activation is slowing of cellular proliferation rate in the face of limited oxygen availability (Hubbi & Semenza, 2015), which we found here (**Figs 1B, 2B, 5A**). The effects of HIF-1 on cell proliferation rate are mediated, in part, by increased expression of cyclin-dependent kinase inhibitor p21 (*CDKN1A*), inhibition of E2F targets (Gardner *et al*, 2001), and inhibition of pro-proliferative MYC signaling (Koshiji *et al*, 2004). These transcriptional effects are precisely what we observed in BAY treated LFs in normoxia. By contrast, hypoxia culture was associated with decreased expression of p21, consistent with a previous report (Mizuno *et al*, 2009), as well as increased expression of MYC protein and enrichment of MYC target genes. Indeed, the most marked differences between hypoxia and BAY treatment on LF gene transcription were the up-regulation of pro-proliferative gene sets containing E2F targets and G2/M checkpoint proteins. Much of this transcriptional response may be mediated by hypoxia-induced up-regulation of MYC, which is known to stimulate cell cycle progression through its effects on the expression and activity of cyclins, cyclin-dependent kinases, and cyclin-dependent kinase inhibitors (Hydbring *et al*, 2017). Clarifying the complex interactions among HIFs, MYC, and cell proliferation will be important for understanding the cellular response of these mesenchymal cells to tissue injury.

Taken together, these findings raise important questions regarding the cell-autonomous role of HIFs in the hypoxia response. On an organismal level, HIFs drive expression of angiogenic and erythropoietic factors to increase oxygen delivery to hypoxic tissues. Within individual cells, HIF-1α seems to be important for mitigating the adverse effects of ROS formation by dysfunctional electron transport in the mitochondria. Indeed, hypoxia increased oxygen consumption and ROS production in HIF-1α-null mouse embryonic fibroblasts (MEFs), which was associated with increased cell death (Zhang *et al*, 2008). Interestingly, these cells also had increased ATP levels compared to wild type, suggesting that mitochondrial function was adequate under 1% oxygen culture conditions to support oxidative phosphorylation and to meet the energy needs of the cells. Given the prominence of HIFs in mediating the transcriptional response to hypoxia, it is somewhat surprising that neither PHD, HIFs, nor their downstream targets were found to be selectively essential as a function of oxygen tension in a genome-wide CRISPR growth screen of K562 human lymphoblasts cultured in normoxia or hypoxia (Jain *et al*, 2020). Similarly, knockout of HIF signaling did not affect growth, internal metabolite concentrations, glucose consumption, or lactate production under hypoxia by human acute myeloid leukemia cells (Wierenga *et al*, 2019). Together with our results, these studies highlight the need for additional research linking hypoxia-induced metabolic changes to their transcriptional and post-transcriptional regulatory mechanisms, particularly in primary cells.

This work also highlights two specific metabolic features that appear to be important in the metabolic response of primary cells to hypoxia. First, both LFs and PASMCs demonstrated notable incorporation of lactate-derived carbon into intracellular metabolic pathways that increased with hypoxia and BAY treatments (**Figs 4, S4, S5**). This finding is consistent with increasing evidence suggesting an important role for lactate as a metabolic fuel in several organ systems (Faubert *et al*, 2013; Hui *et al*, 2017). Although typically considered a metabolic waste product (Rabinowitz & Enerback, 2020), an important contribution of lactate *import* in supporting metabolic homeostasis in the face of an ischemic insult, which is associated with increased extracellular lactate, is an evolutionarily attractive hypothesis that merits further investigation. Second, PASMCs, but not LFs, demonstrated significant rates of reductive carboxylation that increased in 0.5% oxygen (**Fig S5**). Reductive carboxylation was first identified in hypoxic tumor cells where stable isotope tracing revealed ^13^C incorporation from labeled glutamine into lipids (Metallo *et al*, 2011; Gameiro *et al*, 2013a; Scott *et al*, 2011; Wise *et al*, 2011). Hypoxia drives PASMC proliferation *in vivo* contributing to the development pulmonary hypertension in humans and animal models. Isocitrate dehydrogenase has previously been implicated in the pathobiology of this disease (Fessel *et al*, 2012), and our findings suggest that reductive carboxylation catalyzed by isocitrate dehydrogenase may be a metabolic vulnerability of hypoxic PASMCs associated with pulmonary vascular disease.

Our finding that hypoxia was associated with decreased glycolysis and lactate fermentation was unexpected. Several aspects of our experimental design may have contributed to this finding. First, our goal was to understand how metabolic reprogramming may support cell proliferation in hypoxia. Thus, we measured metabolite fluxes in cells during the exponential growth phase accounting for cell growth rate, metabolite degradation rates, and medium evaporation with multiple measurements over a 72 h time course. Often, cells are studied near confluence, where metabolic contributions to biomass production are less and the rate of glycolysis in hypoxia may be higher. Second, we began our experimental treatments 24 h prior to collecting samples to ensure that the hypoxia metabolic program was established prior to labeling. Similar studies (Metallo *et al*, 2011; Grassian *et al*, 2014) typically placed cells into hypoxia at the time of labeling. Third, and perhaps most importantly, these flux determinations were performed in human primary cell cultures rather than immortalized cell lines. Although both cell types used in this study were derived from lung, we anticipate that many of our findings will be generalizable to primary cells from different tissues.

In summary, in this metabolic flux analysis of proliferating human primary cells *in vitro*, we have demonstrated that MYC uncouples an increase in HIF-dependent glycolytic gene transcription from glycolytic flux in hypoxia. Indeed, the degree of metabolic reprogramming in hypoxia was modest and suggests close coupling between proliferation and metabolism. In light of our findings, additional studies are warranted to clarify the role of HIFs in mediating the metabolic response to hypoxia, to determine how MYC activity is regulated by hypoxia, and to identify other key regulators of hypoxic metabolic reprogramming in primary cells. Moreover, these data strongly caution investigators against drawing conclusions about metabolite flux from measures of gene transcription alone.

## MATERIALS AND METHODS

### Cell culture

Primary normal human lung fibroblasts (LFs) were purchased from Lonza (CC-2512) and cultured in FGM-2 (Lonza CC-3132). Cells from two donors were used in these studies: #33652 (56 y.o., male) and #29132 (19 y.o., female). Primary human pulmonary artery smooth muscle cells were purchased from Lonza (CC-2581) and cultured in SmGM-2 (Lonza CC-3182). Cells from three donors were used in these studies: #30020 (64 y.o., male), #27662 (35 y.o., male), #26698 (51 y.o., male), and #19828 (51 y.o., male). Cell authentication was performed by the vendor. Cells were maintained in a standard tissue culture incubator in 5% CO_2_ at 37 °C.

### Metabolic flux protocol

For extracellular flux measurements, cells were seeded in either standard growth medium or MCDB131 medium lacking glucose, glutamine, and phenol red (genDEPOT) which was supplemented with 2% dialyzed fetal bovine serum (Mediatech) and naturally labeled glucose and glutamine (“light” labeling medium). For LFs, glucose was supplemented at 8 mM and glutamine was supplemented at 1 mM. For PASMCs, glucose was supplemented at 5.55 mM and glutamine was supplemented at 10 mM. These concentrations match the concentrations of these substrates determined in standard growth medium. Preliminary experiments were performed to identify the optimal cell seeding density, exponential growth phase, and labeling duration consistent with metabolic and isotopic steady state. On Day -1, 25,000 cells were seeded in a 35 mm dish in “light” labeling medium. Hypoxia-treated cells were transferred to a tissue culture glovebox set at 0.5% oxygen and 5% CO_2_ (Coy Lab Products). Medium was supplemented with DMSO 0.1% or BAY (10 μM) for these treatment conditions. On Day 0, cells were washed with PBS and the medium was changed to either “light” labeling medium for flux measurements or “heavy” labeling medium containing [1,2-^13^C_2_]-glucose, [U-^13^C_6_]-glucose, [U-^13^C_5_]-glutamine, or [U-^13^C_2_]-lactate for tracer experiments. For LFs, samples were collected on Day 0 and every 24 h for 72 h. For PASMCs, samples were collected on Day 0 and every 12 h for 48 h. Medium and cell lysates were collected at each time point for intra- and extracellular metabolite measurements and total DNA quantification. Dishes without cells were weighed daily to correct for evaporative medium losses and to empirically determine degradation and accumulation rates of metabolites. Medium samples and cell lysates for DNA measurement were stored at −80 °C until analysis. Each individual experiment included triplicate wells for each treatment and time point, and each experiment was repeated 4-8 times.

### Cell count

Direct cell counts of trypsinized cell suspensions in PBS were obtained following staining with propidium iodide and acridine orange using a LUNA-FL fluorescence cell counter (Logos Biosystems). Indirect cell counts for flux measurements were interpolated from total DNA quantified using the Quant-iT PicoGreen dsDNA Assay Kit (Thermo). Cells were washed once with two volumes of PBS, lysed with Tris-EDTA buffer containing 2% Triton X-100, and collected by scraping. Total DNA in 10 μL of lysate was determined by adding 100 μL of 1X PicoGreen dye in Tris-EDTA buffer and interpolating the fluorescence intensity with a standard curve generated using the λ DNA standard. Cell counts were interpolated from a standard curve of DNA obtained from known cell numbers seeded in basal medium (**Fig S1B-C**). No difference in total cellular DNA was identified between normoxia and hypoxia cultures (**Fig S1D**).

### Immunoblots

Cells were washed with one volume of PBS and collected by scraping in PBS. Cell suspensions were centrifuged at 5,000 ×*g* for 5 min at 4 °C. Pellets were lysed in buffer containing Tris 10 mM, pH 7.4, NaCl 150 mM, EDTA 1 mM, EGTA 1 mM, Triton X-100 1% v/v, NP-40 0.5% v/v, and Halt Protease Inhibitor Cocktail (Thermo). Protein concentrations were determined by BCA Protein Assay (Thermo). Lysates were normalized for protein concentration and subjected to SDS-PAGE separation on stain-free tris-glycine gels (Bio-Rad), cross-linked and imaged with the Chemidoc system (Bio-Rad), transferred to PVDF membranes with the Trans-Blot Turbo transfer system (Bio-Rad), imaged, blocked in 5% blocking buffer (Bio-Rad), blotted in primary and secondary antibodies, and developed using WesternBright ECL (Advansta). Band signal intensity was normalized to total protein per lane as determined from the stain-free gel or membrane images.

### RT-qPCR

Total RNA was isolated from cells with the RNeasy Mini Kit (Qiagen). cDNA was synthesized from 0.25-1.00 ng RNA with the High Capacity cDNA Reverse Transcription Kit (Applied Biosystems). RT-qPCR analysis was performed with an Applied Biosystems 7500 Fast Real Time PCR System with TaqMan Universal PCR Master Mix and pre-designed TaqMan gene expression assays (Life Technologies). Relative expression levels were calculated using the comparative cycle threshold method referenced to *ACTB*.

### Glucose assay

Medium samples were diluted 10-fold in PBS. Glucose concentration was determined using the Glucose Colorimetric Assay Kit (Cayman) according to the manufacturer’s protocol. Standards were prepared in PBS.

### Lactate assay

Medium samples were diluted 10-fold in PBS. Glucose concentration was determined using the ʟ-Lactate Assay Kit (Cayman). Medium samples did not require deproteinization, otherwise the samples were analyzed according to the manufacturer’s protocol. Standards were prepared in PBS.

### Pyruvate assay

Pyruvate was measured using either an enzymatic assay (most samples) or an HPLC-based assay (medium from 0.2% oxygen experiments). For the enzymatic assay, medium samples were diluted 20-fold in PBS. Pyruvate concentration was determined using the Pyruvate Assay Kit (Cayman). Medium samples did not require deproteinization, otherwise the samples were analyzed according to the manufacturer’s protocol. Standards were prepared in PBS. For the HPLC assay, 2-oxovaleric acid was added to medium samples as an internal standard. Samples were subsequently deproteinized with 2 volumes of ice-cold acetone. Supernatants were evaporated to < 50% of the starting volume at 43 °C in a SpeedVac concentrator (Thermo Savant) and reconstituted to the starting volume with HPLC-grade water prior to derivatization. Samples were derivatized 1:1 by volume with *o*-phenylenediamine (25 mM in 2 M HCl) for 30 min at 80 °C. Derivatized pyruvate was separated with a Poroshell HPH C-18 column (2.1 × 100 mm, 2.7 μm) on an Infinity II high-performance liquid chromatography system with fluorescence detection of OPD-derivatized α-keto acids as described previously (Guarino *et al*, 2019).

### Amino acid assay

Medium amino acid concentrations were determined following the addition of norvaline and sarcosine internal standards and deproteinization with 2 volumes of ice-cold acetone. Supernatants were evaporated to < 50% of the starting volume at 43 °C in a SpeedVac concentrator (Thermo Savant) and reconstituted to the starting volume with HPLC-grade water prior to analysis. Amino acids in deproteinized medium were derivatized with *o*-phthalaldehyde (OPA) and 9-fluorenylmethylchloroformate (FMOC) immediately prior to separation with a Poroshell HPH-C18 column (4.6 × 100 mm, 2.7 μm) on an Infinity II high-performance liquid chromatography system with ultraviolet and fluorescence detection of OPA- and FMOC-derivatized amino acids, respectively, according to the manufacturer’s protocol (Agilent) (Long, 2017).

### MYC knockdown

Approximately 1.25 M LFs were reverse transfected in 6-cm dishes with 40 pmol siMYC or non-targeting siCTL pools (Dharmacon) and 20 μL RNAiMAX (Thermo) in 500 μL OptiMEM (Thermo). After 24 h, cells were collected by trypsinization and re-seeded as described in *Metabolic flux protocol* above for growth rate and lactate efflux measurements.

### MYC overexpression

LFs were seeded at 25,000 cells per 35 mm dish on Day -2. On Day -1, cells were transduced with adenovirus for MYC (Vector Biolabs) or YFP overexpression (Oldham *et al*, 2015). After 24 h, the medium was changed and samples were collected as described in *Metabolic flux protocol* above.

### Flux calculations

The growth rate (𝜇) and flux (*v*) for each measured metabolite were defined as follows (Murphy & Young, 2013):

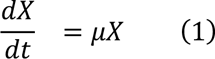

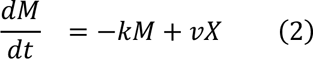

where *X* is the cell density, *k* is the first-order degradation or accumulation rate, and *M* is the mass of the metabolite. These equations are solved as follows:

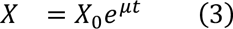

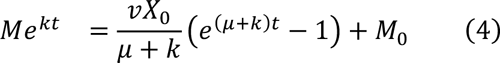

Growth rate (𝜇) and cell count at time 0 (𝑋_0_) were determined by robust linear modeling of the logarithm of cell count as a function of time (𝑡). Metabolite mass was calculated from the measured metabolite concentrations and predicted well volume accounting for evaporative losses (**Fig S1E**). First-order degradation and accumulation rates were obtained from robust linear modeling of metabolite mass *v*. time in unconditioned culture medium. Rates that significantly differed from 0 by Student’s *t*-test were incorporated into the flux calculations. Final fluxes were obtained by robust linear modeling of 𝑀𝑒^𝑘𝑡^ *versus* (𝑒^(𝜇+𝑘)𝑡^ − 1) to determine the slope from which 𝑣 was calculated using equation (4).

### Metabolomics

#### Metabolite extraction

Intracellular metabolites were obtained after washing cells with 2 volumes of ice-cold PBS and floating on liquid nitrogen. Plates were stored at −80 °C until extraction. Metabolites were extracted with 1 mL 80% MeOH pre-cooled to −80 °C containing 10 nmol [D_8_]-valine as an internal standard (Cambridge Isotope Labs). Insoluble material was removed by centrifugation at 21,000 ×*g* for 15 min at 4 °C. The supernatant was evaporated to dryness at 42 °C using a SpeedVac concentrator (Thermo Savant). Samples were resuspended in 35 μL LC-MS-grade water prior to analysis.

#### Acquisition parameters

LC-MS analysis was performed on a Vanquish ultra-high-performance liquid chromatography system coupled to a Q Exactive orbitrap mass spectrometer by a HESI-II electrospray ionization probe (Thermo). External mass calibration was performed weekly. Metabolite samples (2.5 μL) were separated using a ZIC-pHILIC stationary phase (2.1 × 150 mm, 5 μm) (Merck). The autosampler temperature was 4 °C and the column compartment was maintained at 25 °C. Mobile phase A was 20 mM ammonium carbonate and 0.1% ammonium hydroxide. Mobile phase B was acetonitrile. The flow rate was 0.1 mL/min. Solvent was introduced to the mass spectrometer *via* electrospray ionization with the following source parameters: sheath gas 40, auxiliary gas 15, sweep gas 1, spray voltage +3.0 kV for positive mode and −3.1 kV for negative mode, capillary temperature 275 °C, S-lens RF level 40, and probe temperature 350 °C. Data were acquired and peaks integrated using TraceFinder 4.1 (Thermo).

#### Stable isotope quantification

All metabolites except fructose 2,6-bisphosphate (FBP) and 3-phosphoglycerate (3PG) were measured using the following mobile phase gradient: 0 min, 80% B; 5 min, 80% B; 30 min, 20% B; 31 min, 80% B; 42 min, 80% B. The mass spectrometer was operated in selected ion monitoring mode with an m/*z* window width of 9.0 centered 1.003355-times half the number of carbon atoms in the target metabolite. The resolution was set at 70,000 and AGC target was 1×10^5^ ions. Peak areas were corrected for quadrupole bias as in Kim *et al*. (2015). Mass isotope distributions for FBP and 3PG were calculated from full scan chromatograms as described below. Raw mass isotopomer distributions were corrected for natural isotope abundance using a custom R package (mzrtools, https://github.com/oldhamlab/mzrtools) employing the method of Fernandez, *et al*. (1996).

#### Metabolomic profiling

For metabolomic profiling and quantification of isotopic enrichment for FBP and 3PG, the following mobile phase gradient was used: 0 min, 80% B; 20 min, 20% B; 20.5 min, 80% B; 28 min, 80% B; 42 min, 80% B. The mass spectrometer was operated in polarity switching full scan mode from 70-1000 m/*z*. Resolution was set to 70,000 and the AGC target was 1×10^6^ ions. Peak identifications were based on an in-house library of authentic metabolite standards previously analyzed utilizing this method. For metabolomics studies, pooled quality control (QC) samples were injected at the beginning, end, and between every four samples of the run. Raw peak areas for each metabolite were corrected for instrument drift using a cubic spline model of QC peak areas. Low quality features were removed on the basis of a relative standard deviation greater than 0.2 in the QC samples and a dispersion ratio greater than 0.4 (Broadhurst *et al*, 2018). Missing values were imputed using random forest. Samples peak areas were normalized using probabilistic quotient normalization (Dieterle *et al*, 2006). Differentially regulated metabolites were identified using limma (Ritchie *et al*, 2015). Metabolite set enrichment analysis was performed using the fgsea package (Korotkevich *et al*, 2021) with KEGG metabolite pathways (Kanehisa & Goto, 2000).

### Biomass determination

The dry weight of LFs was determined to be 493 pg/cell. The dry weight of PASMCs was determined to be 396 pg/cell. These values were estimated by washing 3 × 10^6^ cells twice in PBS and thrice in ice-cold acetone prior to drying overnight in a SpeedVac. The composition of the dry cell mass was estimated from the literature (Quek *et al*, 2010; Sheikh *et al*, 2005), and stoichiometric coefficients were determined as described (Murphy *et al*, 2013; Zamorano *et al*, 2010).

### Metabolic flux analysis

Metabolic flux analysis was performed using the elementary metabolite unit-based software package INCA (Young, 2014). Inputs to the model include the chemical reactions and atom transitions of central carbon metabolism, extracellular fluxes, the identity and composition of ^13^C-labeled tracers, and the MIDs of labeled intracellular metabolites. The metabolic network was adapted from previously published networks (Murphy *et al*, 2013; Vacanti *et al*, 2014) and comprises 48 reactions representing glycolysis, the pentose phosphate pathway, the tricarboxylic acid cycle, anaplerotic pathways, serine metabolism, and biomass synthesis. The network includes seven extracellular substrates (aspartate, cystine, glucose, glutamine, glycine, pyruvate, serine) and five metabolic products (alanine, biomass, glutamate, lactate, lipid). Models were fit using three ^13^C-labeled tracers, [1,2-^13^C_2_] glucose, [U-^13^C_6_] glucose, and [U-^13^C_5_] glutamine. The MIDs of twelve metabolites (2-oxoglutarate, 3-phosphoglycerate, alanine, aspartate, citrate, fructose bisphosphate, glutamate, glutamine, lactate, malate, pyruvate, serine) were used to constrain intracellular fluxes. The following assumptions were made:

1. Metabolism was at steady state.
2. Labeled CO_2_ produced during decarboxylation reactions left the system and did not re-incorporate during carboxylation reactions.
3. Protein turnover occurred at a negligible rate compared to glucose and glutamine consumption.
4. Acetyl-CoA and pyruvate existed in cytosolic and mitochondrial pools. Aspartate, fumarate, oxaloacetate, and malate were allowed to exchange freely between the compartments.
5. The per cell biomass requirements of LFs and PASMCs were similar to published estimates from other cell types (Quek *et al*, 2010; Sheikh *et al*, 2005).
6. Succinate and fumarate are symmetric molecules that have interchangeable orientations when metabolized by TCA cycle enzymes.

Flux estimation was repeated a minimum of 50 times from random initial values. Results were subjected to a χ^2^ statistical test to assess goodness-of-fit. Accurate 95% confidence intervals were computed for estimated parameters by evaluating the sensitivity of the sum-of-square residuals to parameter variations (Antoniewicz *et al*, 2006; Murphy *et al*, 2013).

### NAD(H) assay

Cellular NAD^+^ and NADH were measured using an enzymatic fluorometric cycling assay based on the reduction of NAD^+^ to NADH by alcohol dehydrogenase (ADH) and subsequent electron transfer to generate the fluorescent molecule resorufin (Oldham *et al*, 2015). Briefly, cells were washed twice with one volume PBS. Pyridine nucleotides were extracted on ice with buffer containing 50% by volume PBS and 50% lysis buffer (100 mM sodium carbonate, 20 mM sodium bicarbonate, 10 mM nicotinamide, 0.05% by volume Triton-X-100, 1% by mass dodecyltrimethylammonium bromide) and collected by scraping. Extracts were divided equally and 0.5 volume of 0.4 N HCl was added to one sample. Both extracts were heated at 65 °C for 15 min to degrade selectively either the oxidized (buffer) or reduced (HCl) nucleotides. The reaction was cooled on ice and quenched by adding 0.5 M Tris-OH to the acid-treated samples or 0.2 N HCl plus 0.25 M Tris-OH to the buffer samples. Samples were then diluted in reaction buffer (50 mM EDTA and 10 mM Tris, pH 7.06). Cell debris was pelleted by centrifugation, and 50 μL was incubated for 2 h with 100 μL reaction buffer containing 0.6 M EtOH, 0.5 mM phenazine methosulfate, 0.05 mM resazurin, and 0.1 mg/mL ADH. Fluorescence intensities were measured with a Spectramax Gemini XPS (Molecular Devices) with excitation 540 nm, emission 588 nm, and 550 nm excitation cut-off filter. Sample intensities were compared to a standard curve generated from known concentrations of NADH. The ratio of fluorescence in buffer-extracted to acid-extracted samples corresponds to the NADH/NAD^+^ ratio. Absolute NADH and NAD^+^ were normalized to estimated cell counts from total DNA quantification as described above.

### RNA-seq

RNA was collected from LFs treated for three days ± hypoxia ± BAY as described above. Four biological replicates were analyzed. Library construction and sequencing was performed by BGI Genomics using 100 bp paired end analysis and a read depth of 50 M reads per sample. Sequences were deposited in the NIH SRA (PRJNA721596). Sequences were mapped to the human GRCh38 primary assembly and counts summarized using Rsubread (Liao *et al*, 2019). This data is available from the Oldham Lab GitHub repository (https://github.com/oldhamlab/rnaseq.lf.hypoxia.molidustat). Differentially expressed transcripts were identified using DESeq2 (Love *et al*, 2014). Gene set enrichment and transcription factor enrichment was performed using the fgsea and DoRothEA R packages, respectively (Korotkevich *et al*, 2021; Garcia-Alonso *et al*, 2019).

### Quantification and Statistical Analysis

The raw data and annotated analysis code necessary to reproduce this manuscript are contained in an R package research compendium available from the Oldham Lab GitHub repository (https://github.com/oldhamlab/Copeland.2022.hypoxia.flux). Data analysis, statistical comparisons, and visualization were performed in R (R Core Team, 2022) using the packages referenced in the data supplement. Experiments included technical and biological replicates as noted above. The number of biological replicates (n) is indicated in the figure legends. Summary data show the mean ± SEM. Outliers were identified using twice the median absolute deviation as a cutoff threshold. Comparisons were performed using linear mixed-effects models with oxygen, treatment, and their interaction as fixed effects and biological replicate as a random effect. Significant differences in estimated marginal means were identified by comparisons to the multivariate *t* distribution. Metabolomics and RNA-seq data were analyzed as described above. Probability values less than 0.05 were considered statistically significant.

### Key Resources Table

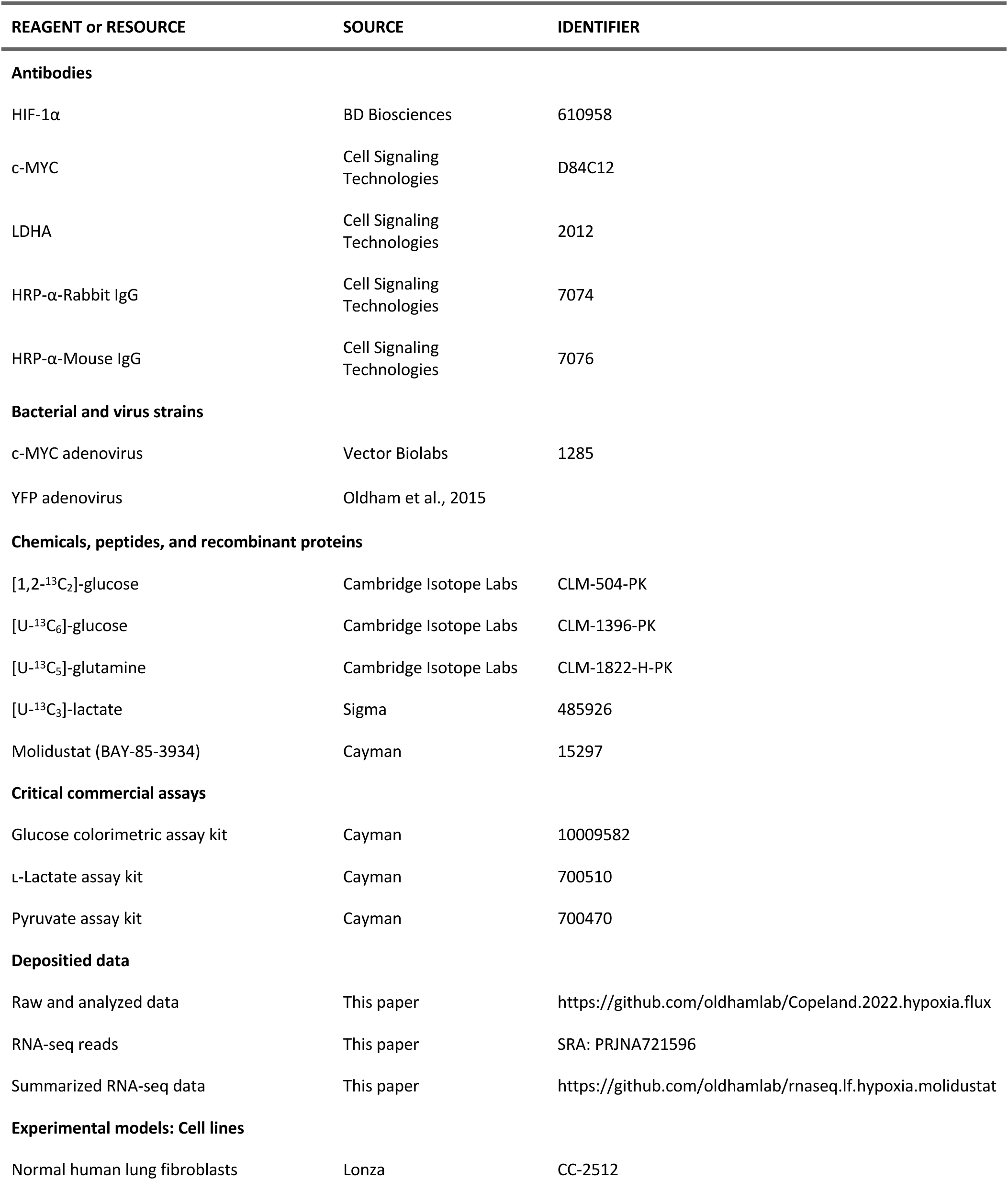
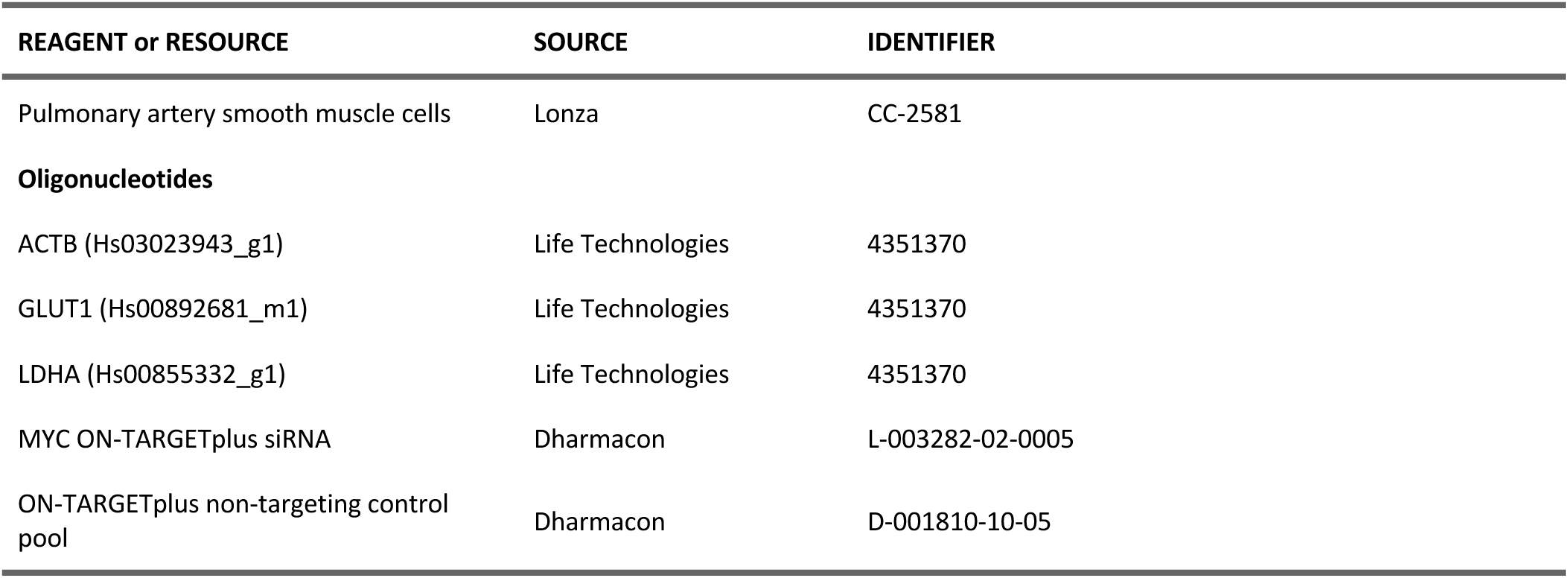

## ACKNOWLEDGEMENTS

This work was supported by grants from the NIH (K08HL128802), American Lung Association, Pulmonary Hypertension Association, and the American Thoracic Society Foundation to W.M.O and from the NIH (U01HG007690, U54HL119145, R01HL155107, R01HL155096) and the American Heart Association (D700382, CV-19) to J.L.

## AUTHOR CONTRIBUTIONS

W.M.O. conceived and designed the analysis. C.A.C., B.A.O., D.R.Z., S.M., K.L., and W.M.O. collected the data. J.D.Y. and W.M.O. contributed data or analysis tools. W.M.O. performed the analysis. W.M.O. drafted the manuscript. All authors participated in interpreting the results and revising the manuscript. All authors approve the final submission.

## CONFLICT OF INTEREST

The authors declare that they have no conflicts of interest.

## Notes

https://github.com/oldhamlab/Copeland.2022.hypoxia.flux

### Competing Interest Statement

The authors have declared no competing interest.

### Summary of Updates

Revised figure presentation. Revised results text.

## REFERENCES

Ahn WS & Antoniewicz MR (2011) Metabolic flux analysis of CHO cells at growth and non-growth phases using isotopic tracers and mass spectrometry. Metab Eng 13: 598–609

Antoniewicz MR (2018) A guide to 13C metabolic flux analysis for the cancer biologist. Exp Mol Med 50: 1–13

Antoniewicz MR, Kelleher JK & Stephanopoulos G (2006) Determination of confidence intervals of metabolic fluxes estimated from stable isotope measurements. Metab Eng 8: 324–37

Batie M, Frost J, Frost M, Wilson JW, Schofield P & Rocha S (2019) Hypoxia induces rapid changes to histone methylation and reprograms chromatin. Science 363: 1222–1226

Broadhurst D, Goodacre R, Reinke SN, Kuligowski J, Wilson ID, Lewis MR & Dunn WB (2018) Guidelines and considerations for the use of system suitability and quality control samples in mass spectrometry assays applied in untargeted clinical metabolomic studies. Metabolomics 14: 72

Buescher JM, Antoniewicz MR, Boros LG, Burgess SC, Brunengraber H, Clish CB, DeBerardinis RJ, Feron O, Frezza C, Ghesquiere B, et al (2015) A roadmap for interpreting (13)C metabolite labeling patterns from cells. Curr Opin Biotechnol 34: 189–201

Chance B & Williams GR (1955) Respiratory enzymes in oxidative phosphorylation. III. The steady state. J Biol Chem 217: 409–27

Chandel NS, Maltepe E, Goldwasser E, Mathieu CE, Simon MC & Schumacker PT (1998) Mitochondrial reactive oxygen species trigger hypoxia-induced transcription. Proc Natl Acad Sci U S A 95: 11715–20

Chen B, Calvert AE, Cui H & Nelin LD (2009) Hypoxia promotes human pulmonary artery smooth muscle cell proliferation through induction of arginase. American Journal of Physiology-Lung Cellular and Molecular Physiology 297: L1151–L1159

Chen C, Cai S, Wang G, Cao X, Yang X, Luo X, Feng Y & Hu J (2013) c-Myc enhances colon cancer cell-mediated angiogenesis through the regulation of HIF-1α. Biochem Biophys Res Commun 430: 505–511

Contreras-Baeza Y, Sandoval PY, Alarcon R, Galaz A, Cortes-Molina F, Alegria K, Baeza-Lehnert F, Arce-Molina R, Guequen A, Flores CA, et al (2019) Monocarboxylate transporter 4 (MCT4) is a high affinity transporter capable of exporting lactate in high-lactate microenvironments. J Biol Chem 294: 20135–20147

Crown SB, Marze N & Antoniewicz MR (2015) Catabolism of Branched Chain Amino Acids Contributes Significantly to Synthesis of Odd-Chain and Even-Chain Fatty Acids in 3T3-L1 Adipocytes. PLoS One 10: e0145850

Dang CV (2012) MYC on the path to cancer. Cell 149: 22–35

Dang CV, O’Donnell KA, Zeller KI, Nguyen T, Osthus RC & Li F (2006) The c-Myc target gene network. Semin Cancer Biol 16: 253–264

Dieterle F, Ross A, Schlotterbeck G & Senn H (2006) Probabilistic quotient normalization as robust method to account for dilution of complex biological mixtures. Application in 1H NMR metabonomics. Anal Chem 78: 4281–90

Doe MR, Ascano JM, Kaur M & Cole MD (2012) Myc posttranscriptionally induces HIF1 protein and target gene expression in normal and cancer cells. Cancer Res 72: 949–957

Fan J, Kamphorst JJ, Mathew R, Chung MK, White E, Shlomi T & Rabinowitz JD (2013) Glutamine-driven oxidative phosphorylation is a major ATP source in transformed mammalian cells in both normoxia and hypoxia. Mol Syst Biol 9: 712

Faubert B, Boily G, Izreig S, Griss T, Samborska B, Dong Z, Dupuy F, Chambers C, Fuerth BJ, Viollet B, et al (2013) AMPK is a negative regulator of the Warburg effect and suppresses tumor growth in vivo. Cell Metab 17: 113–24

Faubert B, Li KY, Cai L, Hensley CT, Kim J, Zacharias LG, Yang C, Do QN, Doucette S, Burguete D, et al (2017) Lactate Metabolism in Human Lung Tumors. Cell 171: 358–371 e9

Favaro E, Bensaad K, Chong MG, Tennant DA, Ferguson DJ, Snell C, Steers G, Turley H, Li JL, Gunther UL, et al (2012) Glucose utilization via glycogen phosphorylase sustains proliferation and prevents premature senescence in cancer cells. Cell Metab 16: 751–64

Fernandez CA, Des Rosiers C, Previs SF, David F & Brunengraber H (1996) Correction of 13C mass isotopomer distributions for natural stable isotope abundance. J Mass Spectrom 31: 255–62

Fessel JP, Hamid R, Wittmann BM, Robinson LJ, Blackwell T, Tada Y, Tanabe N, Tatsumi K, Hemnes AR & West JD (2012) Metabolomic analysis of bone morphogenetic protein receptor type 2 mutations in human pulmonary endothelium reveals widespread metabolic reprogramming. Pulm Circ 2: 201–13

Flamme I, Oehme F, Ellinghaus P, Jeske M, Keldenich J & Thuss U (2014) Mimicking hypoxia to treat anemia: HIF-stabilizer BAY 85-3934 (Molidustat) stimulates erythropoietin production without hypertensive effects. PLoS One 9: e111838

Gameiro PA, Laviolette LA, Kelleher JK, Iliopoulos O & Stephanopoulos G (2013a) Cofactor balance by nicotinamide nucleotide transhydrogenase (NNT) coordinates reductive carboxylation and glucose catabolism in the tricarboxylic acid (TCA) cycle. J Biol Chem 288: 12967–77

Gameiro PA, Yang J, Metelo AM, Perez-Carro R, Baker R, Wang Z, Arreola A, Rathmell WK, Olumi A, Lopez-Larrubia P, et al (2013b) In vivo HIF-mediated reductive carboxylation is regulated by citrate levels and sensitizes VHL-deficient cells to glutamine deprivation. Cell Metab 17: 372–85

Garcia-Alonso L, Holland CH, Ibrahim MM, Turei D & Saez-Rodriguez J (2019) Benchmark and integration of resources for the estimation of human transcription factor activities. Genome Res 29: 1363–1375

Garcia-Bermudez J, Baudrier L, La K, Zhu XG, Fidelin J, Sviderskiy VO, Papagiannakopoulos T, Molina H, Snuderl M, Lewis CA, et al (2018) Aspartate is a limiting metabolite for cancer cell proliferation under hypoxia and in tumours. Nat Cell Biol 20: 775–781

Gardner LB, Li Q, Park MS, Flanagan WM, Semenza GL & Dang CV (2001) Hypoxia inhibits G1/S transition through regulation of p27 expression. J Biol Chem 276: 7919–7926

Garofalo O, Cox DW & Bachelard HS (1988) Brain levels of NADH and NAD+ under hypoxic and hypoglycaemic conditions in vitro. J Neurochem 51: 172–6

Gordan JD, Bertout JA, Hu CJ, Diehl JA & Simon MC (2007) HIF-2alpha promotes hypoxic cell proliferation by enhancing c-myc transcriptional activity. Cancer Cell 11: 335–47

Grassian AR, Parker SJ, Davidson SM, Divakaruni AS, Green CR, Zhang X, Slocum KL, Pu M, Lin F, Vickers C, et al (2014) IDH1 mutations alter citric acid cycle metabolism and increase dependence on oxidative mitochondrial metabolism. Cancer Res 74: 3317–31

Guarino VA, Oldham WM, Loscalzo J & Zhang YY (2019) Reaction rate of pyruvate and hydrogen peroxide: assessing antioxidant capacity of pyruvate under biological conditions. Sci Rep 9: 19568

Guzy RD, Hoyos B, Robin E, Chen H, Liu L, Mansfield KD, Simon MC, Hammerling U & Schumacker PT (2005) Mitochondrial complex III is required for hypoxia-induced ROS production and cellular oxygen sensing. Cell Metab 1: 401–8

Hubbi ME & Semenza GL (2015) Regulation of cell proliferation by hypoxia-inducible factors. Am J Physiol Cell Physiol 309: C775–82

Hui S, Cowan AJ, Zeng X, Yang L, TeSlaa T, Li X, Bartman C, Zhang Z, Jang C, Wang L, et al (2020) Quantitative Fluxomics of Circulating Metabolites. Cell Metab

Hui S, Ghergurovich JM, Morscher RJ, Jang C, Teng X, Lu W, Esparza LA, Reya T, Le Z, Yanxiang Guo J, et al (2017) Glucose feeds the TCA cycle via circulating lactate. Nature 551: 115–118

Hydbring P, Castell A & Larsson L-G (2017) MYC Modulation around the CDK2/p27/SKP2 Axis. Genes (Basel) 8: 174

Islam MS, Leissing TM, Chowdhury R, Hopkinson RJ & Schofield CJ (2018) 2-Oxoglutarate-Dependent Oxygenases. Annu Rev Biochem 87: 585–620

Jain IH, Calvo SE, Markhard AL, Skinner OS, To TL, Ast T & Mootha VK (2020) Genetic Screen for Cell Fitness in High or Low Oxygen Highlights Mitochondrial and Lipid Metabolism. Cell 181: 716–727 e11

Jazmin LJ & Young JD (2013) Isotopically nonstationary 13C metabolic flux analysis. Methods Mol Biol 985: 367– 90

Jiang L, Shestov AA, Swain P, Yang C, Parker SJ, Wang QA, Terada LS, Adams ND, McCabe MT, Pietrak B, et al (2016) Reductive carboxylation supports redox homeostasis during anchorage-independent growth. Nature 532: 255–8

Kaelin WG & Ratcliffe PJ (2008) Oxygen sensing by metazoans: the central role of the HIF hydroxylase pathway. Mol Cell 30: 393–402

Kanehisa M & Goto S (2000) KEGG: kyoto encyclopedia of genes and genomes. Nucleic Acids Res 28: 27–30

Kim D, Fiske BP, Birsoy K, Freinkman E, Kami K, Possemato RL, Chudnovsky Y, Pacold ME, Chen WW, Cantor JR, et al (2015) SHMT2 drives glioma cell survival in ischaemia but imposes a dependence on glycine clearance. Nature 520: 363–7

Korotkevich G, Sukhov V, Budin N, Shpak B, Artyomov MN & Sergushichev A (2021) Fast gene set enrichment analysis. 060012 doi:10.1101/060012 [PREPRINT]

Koshiji M, Kageyama Y, Pete EA, Horikawa I, Barrett JC & Huang LE (2004) HIF-1alpha induces cell cycle arrest by functionally counteracting Myc. EMBO J 23: 1949–1956

Koshiji M, To KK-W, Hammer S, Kumamoto K, Harris AL, Modrich P & Huang LE (2005) HIF-1alpha induces genetic instability by transcriptionally downregulating MutSalpha expression. Mol Cell 17: 793–803

Le A, Lane AN, Hamaker M, Bose S, Gouw A, Barbi J, Tsukamoto T, Rojas CJ, Slusher BS, Zhang H, et al (2012) Glucose-independent glutamine metabolism via TCA cycling for proliferation and survival in B cells. Cell Metab 15: 110–21

Lee JW, Ko J, Ju C & Eltzschig HK (2019a) Hypoxia signaling in human diseases and therapeutic targets. Exp Mol Med 51: 1–13

Lee P, Chandel NS & Simon MC (2020) Cellular adaptation to hypoxia through hypoxia inducible factors and beyond. Nat Rev Mol Cell Biol 21: 268–283

Lee WD, Mukha D, Aizenshtein E & Shlomi T (2019b) Spatial-fluxomics provides a subcellular-compartmentalized view of reductive glutamine metabolism in cancer cells. Nat Commun 10: 1351

Li F, Wang Y, Zeller KI, Potter JJ, Wonsey DR, O’Donnell KA, Kim J-W, Yustein JT, Lee LA & Dang CV (2005) Myc stimulates nuclearly encoded mitochondrial genes and mitochondrial biogenesis. Mol Cell Biol 25: 6225–6234

Li H, Meininger CJ, Hawker JR, Haynes TE, Kepka-Lenhart D, Mistry SK, Morris SM & Wu G (2001) Regulatory role of arginase I and II in nitric oxide, polyamine, and proline syntheses in endothelial cells. American Journal of Physiology-Endocrinology and Metabolism 280: E75–E82

Li Y, Sun X-X, Qian DZ & Dai M-S (2020) Molecular Crosstalk Between MYC and HIF in Cancer. Front Cell Dev Biol 8: 590576

Liao Y, Smyth GK & Shi W (2019) The R package Rsubread is easier, faster, cheaper and better for alignment and quantification of RNA sequencing reads. Nucleic Acids Res 47: e47

Liberzon A, Birger C, Thorvaldsdottir H, Ghandi M, Mesirov JP & Tamayo P (2015) The Molecular Signatures Database (MSigDB) hallmark gene set collection. Cell Syst 1: 417–425

Liu S-S, Wang H-Y, Tang J-M & Zhou X-M (2013) Hypoxia-induced collagen synthesis of human lung fibroblasts by activating the angiotensin system. Int J Mol Sci 14: 24029–24045

Long W (2017) Automated amino acid analysis using an Agilent Poroshell HPH-C18 column. *Application Note, Agilent Technologies, Inc* Publication Number 5991-5571EN: 1–10

Love MI, Huber W & Anders S (2014) Moderated estimation of fold change and dispersion for RNA-seq data with DESeq2. Genome Biol 15: 550

Madden SK, de Araujo AD, Gerhardt M, Fairlie DP & Mason JM (2021) Taking the Myc out of cancer: toward therapeutic strategies to directly inhibit c-Myc. Molecular Cancer 20: 3

Mann G, Mora S, Madu G & Adegoke OAJ (2021) Branched-chain Amino Acids: Catabolism in Skeletal Muscle and Implications for Muscle and Whole-body Metabolism. Front Physiol 12: 702826

Masson N, Keeley TP, Giuntoli B, White MD, Puerta ML, Perata P, Hopkinson RJ, Flashman E, Licausi F & Ratcliffe PJ (2019) Conserved N-terminal cysteine dioxygenases transduce responses to hypoxia in animals and plants. Science 365: 65–69

Melendez-Rodriguez F, Urrutia AA, Lorendeau D, Rinaldi G, Roche O, Bogurcu-Seidel N, Ortega Muelas M, Mesa-Ciller C, Turiel G, Bouthelier A, et al (2019) HIF1alpha Suppresses Tumor Cell Proliferation through Inhibition of Aspartate Biosynthesis. Cell Rep 26: 2257–2265 e4

Metallo CM, Gameiro PA, Bell EL, Mattaini KR, Yang J, Hiller K, Jewell CM, Johnson ZR, Irvine DJ, Guarente L, et al (2011) Reductive glutamine metabolism by IDH1 mediates lipogenesis under hypoxia. Nature 481: 380–4

Mizuno S, Bogaard HJ, Voelkel NF, Umeda Y, Kadowaki M, Ameshima S, Miyamori I & Ishizaki T (2009) Hypoxia regulates human lung fibroblast proliferation via p53-dependent and -independent pathways. Respir Res 10: 17

Murphy TA, Dang CV & Young JD (2013) Isotopically nonstationary 13C flux analysis of Myc-induced metabolic reprogramming in B-cells. Metab Eng 15: 206–17

Murphy TA & Young JD (2013) ETA: robust software for determination of cell specific rates from extracellular time courses. Biotechnol Bioeng 110: 1748–58

Oldham WM, Clish CB, Yang Y & Loscalzo J (2015) Hypoxia-Mediated Increases in L-2-hydroxyglutarate Coordinate the Metabolic Response to Reductive Stress. Cell Metab 22: 291–303

Owczarek A, Gieczewska K, Jarzyna R, Jagielski AK, Kiersztan A, Gruza A & Winiarska K (2020) Hypoxia increases the rate of renal gluconeogenesis via hypoxia-inducible factor-1-dependent activation of phosphoenolpyruvate carboxykinase expression. Biochimie 171–172: 31–37

Pelletier J, Bellot G, Gounon P, Lacas-Gervais S, Pouyssegur J & Mazure NM (2012) Glycogen Synthesis is Induced in Hypoxia by the Hypoxia-Inducible Factor and Promotes Cancer Cell Survival. Front Oncol 2: 18

Quek LE, Dietmair S, Kromer JO & Nielsen LK (2010) Metabolic flux analysis in mammalian cell culture. Metab Eng 12: 161–71

R Core Team (2022) R: A language and environment for statistical computing Vienna, Austria: R Foundation for Statistical Computing

Rabinowitz JD & Enerback S (2020) Lactate: the ugly duckling of energy metabolism. Nat Metab 2: 566–571

Ritchie ME, Phipson B, Wu D, Hu Y, Law CW, Shi W & Smyth GK (2015) limma powers differential expression analyses for RNA-sequencing and microarray studies. Nucleic Acids Res 43: e47

Scott DA, Richardson AD, Filipp FV, Knutzen CA, Chiang GG, Ronai ZA, Osterman AL & Smith JW (2011) Comparative metabolic flux profiling of melanoma cell lines: beyond the Warburg effect. J Biol Chem 286: 42626–34

Semenza GL (2012) Hypoxia-inducible factors in physiology and medicine. Cell 148: 399–408

Sheikh K, Forster J & Nielsen LK (2005) Modeling hybridoma cell metabolism using a generic genome-scale metabolic model of Mus musculus. Biotechnol Prog 21: 112–21

Stine ZE, Walton ZE, Altman BJ, Hsieh AL & Dang CV (2015) MYC, Metabolism, and Cancer. Cancer Discov 5: 1024–1039

Szoka L, Karna E, Hlebowicz-Sarat K, Karaszewski J & Palka JA (2017) Exogenous proline stimulates type I collagen and HIF-1α expression and the process is attenuated by glutamine in human skin fibroblasts. Mol Cell Biochem 435: 197–206

Tilton WM, Seaman C, Carriero D & Piomelli S (1991) Regulation of glycolysis in the erythrocyte: role of the lactate/pyruvate and NAD/NADH ratios. J Lab Clin Med 118: 146–52

Vacanti NM, Divakaruni AS, Green CR, Parker SJ, Henry RR, Ciaraldi TP, Murphy AN & Metallo CM (2014) Regulation of substrate utilization by the mitochondrial pyruvate carrier. Mol Cell 56: 425–35

Vita M & Henriksson M (2006) The Myc oncoprotein as a therapeutic target for human cancer. Semin Cancer Biol 16: 318–330

Wenger RH, Kurtcuoglu V, Scholz CC, Marti HH & Hoogewijs D (2015) Frequently asked questions in hypoxia research. Hypoxia (Auckl) 3: 35–43

Wheaton WW & Chandel NS (2011) Hypoxia. 2. Hypoxia regulates cellular metabolism. Am J Physiol Cell Physiol 300: C385–93

Wiechert W (2007) The Thermodynamic Meaning of Metabolic Exchange Fluxes. Biophysical Journal 93: 2255– 2264

Wierenga ATJ, Cunningham A, Erdem A, Lopera NV, Brouwers-Vos AZ, Pruis M, Mulder AB, Gunther UL, Martens JHA, Vellenga E, et al (2019) HIF1/2-exerted control over glycolytic gene expression is not functionally relevant for glycolysis in human leukemic stem/progenitor cells. Cancer Metab 7: 11

Wise DR, Ward PS, Shay JE, Cross JR, Gruber JJ, Sachdeva UM, Platt JM, DeMatteo RG, Simon MC & Thompson CB (2011) Hypoxia promotes isocitrate dehydrogenase-dependent carboxylation of alpha-ketoglutarate to citrate to support cell growth and viability. Proc Natl Acad Sci U S A 108: 19611–6

Xiao W & Loscalzo J (2020) Metabolic Responses to Reductive Stress. Antioxid Redox Signal 32: 1330–1347

Xue J, Nelin LD & Chen B (2017) Hypoxia induces arginase II expression and increases viable human pulmonary artery smooth muscle cell numbers via AMPKα1 signaling. American Journal of Physiology-Lung Cellular and Molecular Physiology 312: L568–L578

Young JD (2014) INCA: a computational platform for isotopically non-stationary metabolic flux analysis. Bioinformatics 30: 1333–5

Young JD, Allen DK & Morgan JA (2014) Isotopomer measurement techniques in metabolic flux analysis II: mass spectrometry. Methods Mol Biol 1083: 85–108

Zamorano F, Wouwer AV & Bastin G (2010) A detailed metabolic flux analysis of an underdetermined network of CHO cells. J Biotechnol 150: 497–508

Zhang H, Bosch-Marce M, Shimoda LA, Tan YS, Baek JH, Wesley JB, Gonzalez FJ & Semenza GL (2008) Mitochondrial autophagy is an HIF-1-dependent adaptive metabolic response to hypoxia. J Biol Chem 283: 10892–903

Zhang H, Gao P, Fukuda R, Kumar G, Krishnamachary B, Zeller KI, Dang CV & Semenza GL (2007) HIF-1 Inhibits Mitochondrial Biogenesis and Cellular Respiration in VHL-Deficient Renal Cell Carcinoma by Repression of C-MYC Activity. Cancer Cell 11: 407–420

Zhang J, Sattler M, Tonon G, Grabher C, Lababidi S, Zimmerhackl A, Raab MS, Vallet S, Zhou Y, Cartron M-A, et al (2009) Targeting angiogenesis via a c-Myc/hypoxia-inducible factor-1alpha-dependent pathway in multiple myeloma. Cancer Res 69: 5082–5090

