## Supplemental Data for "MYC overrides HIF to regulate proliferating primary cell metabolism in hypoxia"

### TABLES

#### Table 1: Lung fibroblast fluxes in 21% and 0.5% oxygen

| | | 21%[a] | | | 0.5%[b] | | | |
|---|---|---|---|---|---|---|---|---|
| **ID** | **Reaction** | **Flux** | **LB** | **UB** | **Flux** | **LB** | **UB** | **Ratio** |
| **NET** | | | | | | | | |
| **Transport** | | | | | | | | |
| GLUT | GLC.x -> GLC | 5.14e+02 | 5.11e+02 | 5.21e+02 | 4.41e+02 | 4.26e+02 | 4.58e+02 | 0.86 |
| PYRR | PYR.x -> PYR.c | 7.56e+01 | 7.31e+01 | 7.96e+01 | 6.21e+01 | 5.83e+01 | 6.60e+01 | 0.82 |
| MCT | LAC <-> LAC.x | 9.99e+02 | 9.98e+02 | 1.02e+03 | 8.91e+02 | 8.62e+02 | 9.25e+02 | 0.89 |
| ALAR | ALA -> ALA.x | 2.25e+00 | 1.95e+00 | 2.49e+00 | 5.84e-01 | 1.10e-03 | 1.16e+00 | 0.26 |
| GLNR | GLN.x -> GLN | 4.15e+01 | 4.06e+01 | 4.16e+01 | 1.43e+01 | 1.26e+01 | 1.94e+01 | 0.34 |
| GLUR | GLU <-> GLU.x | 1.62e+01 | 1.58e+01 | 1.68e+01 | 7.55e+00 | 6.88e+00 | 8.15e+00 | 0.47 |
| ASPR | ASP -> ASP.x | 2.57e+00 | 2.53e+00 | 2.68e+00 | 1.08e+00 | 4.17e-01 | 1.69e+00 | 0.42 |
| SERR | SER.x -> SER | 1.42e+01 | 1.35e+01 | 1.49e+01 | 5.49e+00 | 4.99e+00 | 6.06e+00 | 0.39 |
| CYSR | CYX.x -> CYS + CYS | 4.41e+00 | 4.23e+00 | 4.58e+00 | 1.65e+00 | 1.32e+00 | 2.08e+00 | 0.37 |
| GLYR | GLY -> GLY.x | 2.05e+00 | 1.90e+00 | 2.15e+00 | 2.60e-01 | 2.00e-02 | 4.92e-01 | 0.13 |
| **Glycolysis** | | | | | | | | |
| HK | GLC -> G6P | 5.14e+02 | 5.11e+02 | 5.21e+02 | 4.41e+02 | 4.26e+02 | 4.58e+02 | 0.86 |
| PGI | G6P <-> F6P | 5.11e+02 | 4.99e+02 | 5.24e+02 | 4.23e+02 | 4.04e+02 | 4.40e+02 | 0.83 |
| PFK | F6P -> FBP | 5.09e+02 | 5.00e+02 | 5.12e+02 | 4.32e+02 | 4.17e+02 | 4.49e+02 | 0.85 |
| ALDO | FBP <-> DHAP + GAP | 5.09e+02 | 5.00e+02 | 5.12e+02 | 4.32e+02 | 4.17e+02 | 4.49e+02 | 0.85 |
| TPI | DHAP <-> GAP | 5.08e+02 | 5.06e+02 | 5.08e+02 | 4.31e+02 | 4.15e+02 | 4.48e+02 | 0.85 |
| GAPDH | GAP <-> 3PG | 1.02e+03 | 9.96e+02 | 1.04e+03 | 8.69e+02 | 8.35e+02 | 9.03e+02 | 0.85 |
| ENO | 3PG -> PEP | 1.01e+03 | 9.99e+02 | 1.03e+03 | 8.68e+02 | 8.36e+02 | 9.00e+02 | 0.86 |
| PK | PEP -> PYR.c | 1.04e+03 | 9.95e+02 | 1.04e+03 | 8.78e+02 | 8.36e+02 | 9.21e+02 | 0.84 |
| LDH | PYR.c <-> LAC | 9.99e+02 | 9.98e+02 | 1.02e+03 | 8.91e+02 | 8.62e+02 | 9.25e+02 | 0.89 |
| GPT1 | PYR.c <-> ALA | 1.19e+01 | 9.12e+00 | 1.19e+01 | 5.55e+00 | -9.08e+02 | 6.13e+00 | 0.47 |
| GPT2 | PYR.m <-> ALA | -2.58e+00 | -4.56e+00 | 2.87e+00 | -2.40e-03 | -3.22e+01 | 9.11e+02 | |
| **Pentose phosphate pathway** | | | | | | | | |
| G6PD | G6P -> P5P + CO2 | 1.26e-07 | 0.00e+00 | 3.91e-01 | 1.62e+01 | 4.41e+00 | 2.89e+01 | 128571428.57 |

| | | 21%[a] | | | 0.5%[b] | | | |
|---|---|---|---|---|---|---|---|---|
| ID | Reaction | Flux | LB | UB | Flux | LB | UB | Ratio |
| TK1 | P5P + P5P <-> S7P + GAP | -9.11e-01 | -9.29e-01 | -8.30e-01 | 4.76e+00 | -1.22e-01 | 9.62e+00 | -5.23 |
| TA | S7P + GAP <-> F6P + E4P | -9.11e-01 | -9.29e-01 | -8.30e-01 | 4.76e+00 | -1.22e-01 | 9.62e+00 | -5.23 |
| TK2 | P5P + E4P <-> F6P + GAP | -9.11e-01 | -9.29e-01 | -8.30e-01 | 4.76e+00 | -1.22e-01 | 9.62e+00 | -5.23 |
| **Anaplerosis** | | | | | | | | |
| PYRT | PYR.c -> PYR.m | 1.16e+02 | 1.16e+02 | 1.19e+02 | 4.42e+01 | 3.82e+01 | 9.58e+02 | |
| PC | PYR.m + CO2 -> OAC | 1.88e+01 | 1.74e+01 | 1.91e+01 | 1.37e+01 | 9.82e+00 | 2.69e+01 | |
| PEPCK | OAC -> PEP + CO2 | 2.56e+01 | 1.58e+01 | 2.57e+01 | 9.66e+00 | 0.00e+00 | 2.60e+01 | |
| ME2 | MAL -> PYR.m + CO2 | 2.05e+00 | 9.51e-02 | 2.68e+00 | 1.00e-07 | 0.00e+00 | 2.25e+01 | |
| ME1 | MAL -> PYR.c + CO2 | 2.78e-02 | 0.00e+00 | 2.63e+01 | 8.71e-05 | 0.00e+00 | 2.52e+01 | |
| FAO | FAO -> AcCoA.m | 1.00e-07 | 0.00e+00 | 2.13e+00 | 6.58e-06 | 0.00e+00 | 7.73e-01 | |
| GLDH | GLU <-> AKG | 1.71e+01 | 1.56e+01 | 1.84e+01 | 9.11e-01 | -6.16e-01 | 7.27e+00 | 0.05 |
| GLS | GLN <-> GLU | 3.78e+01 | 3.60e+01 | 3.86e+01 | 1.17e+01 | 1.01e+01 | 1.70e+01 | 0.31 |
| **Tricarboxylic acid cycle** | | | | | | | | |
| PDH | PYR.m -> AcCoA.m + CO2 | 1.02e+02 | 8.76e+01 | 1.15e+02 | 3.05e+01 | 2.86e+01 | 5.24e+01 | 0.30 |
| CS | AcCoA.m + OAC -> CIT | 1.02e+02 | 8.30e+01 | 1.11e+02 | 3.05e+01 | 2.88e+01 | 5.09e+01 | 0.30 |
| IDH | CIT <-> AKG + CO2 | 2.49e+01 | 2.42e+01 | 2.53e+01 | 1.01e+01 | 8.75e+00 | 1.41e+01 | 0.41 |
| OGDH | AKG -> SUC + CO2 | 4.19e+01 | 4.01e+01 | 4.25e+01 | 1.10e+01 | 7.87e+00 | 2.02e+01 | 0.26 |
| SDH | SUC <-> FUM | 4.19e+01 | 4.01e+01 | 4.25e+01 | 1.10e+01 | 7.87e+00 | 2.02e+01 | 0.26 |
| FH | FUM <-> MAL | 4.19e+01 | 4.01e+01 | 4.25e+01 | 1.10e+01 | 7.87e+00 | 2.02e+01 | 0.26 |
| MDH | MAL <-> OAC | 1.17e+02 | 1.08e+02 | 1.24e+02 | 3.14e+01 | 2.62e+01 | 5.70e+01 | 0.27 |
| GOT | OAC <-> ASP | 8.11e+00 | 8.06e+00 | 8.23e+00 | 4.98e+00 | 4.32e+00 | 5.64e+00 | 0.61 |
| **Amino acid metabolism** | | | | | | | | |
| PST | 3PG -> SER | 1.95e+00 | 1.63e+00 | 2.00e+00 | 2.42e-01 | 1.34e-01 | 3.57e+01 | |
| SHT | SER <-> GLY + MEETHF | 6.38e+00 | 6.22e+00 | 6.43e+00 | 3.91e+00 | 3.71e+00 | 4.10e+00 | 0.61 |
| CYST | SER <-> CYS | -7.12e+00 | -7.19e+00 | -6.81e+00 | -2.10e+00 | -2.97e+00 | -1.44e+00 | 0.30 |
| SD | SER -> PYR.c | 1.17e+01 | 1.04e+01 | 1.20e+01 | 2.82e-01 | 0.00e+00 | 1.47e+00 | 0.02 |

| | | 21%[a] | | | 0.5%[b] | | | |
|---|---|---|---|---|---|---|---|---|
| ID | Reaction | Flux | LB | UB | Flux | LB | UB | Ratio |
| GLYS | CO2 + MEETHF -> GLY | 3.39e+00 | 3.35e+00 | 3.49e+00 | 1.80e+00 | 1.66e+00 | 1.93e+00 | 0.53 |
| **Biomass** | | | | | | | | |
| BIOMASS | 1216*AcCoA.c + 295.6*ALA + 232.4*ASP + 114.7*CO2 + 71.43*CYS + 57.14*DHAP + 142.4*G6P + 158.6*GLN + 190.1*GLU + 324.2*GLY + 125.6*MEETHF + 114.7*P5P + 217.2*SER -> biomass | 2.38e-02 | 2.34e-02 | 2.39e-02 | 1.68e-02 | 1.61e-02 | 1.75e-02 | 0.71 |
| ACL | CIT -> AcCoA.c + MAL | 7.74e+01 | 6.29e+01 | 1.04e+02 | 2.04e+01 | 1.95e+01 | 3.71e+01 | 0.26 |
| LIPS | AcCoA.c -> lipid | 4.84e+01 | 4.55e+01 | 4.84e+01 | 1.00e-07 | 0.00e+00 | 1.68e+01 | 0.00 |
| **Mixing** | | | | | | | | |
| cPYR | 0*PYR.c -> PYR.ms | 1.00e+00 | 8.47e-01 | 1.00e+00 | 1.42e-01 | 0.00e+00 | 1.00e+00 | |
| mPYR | 0*PYR.m -> PYR.ms | 1.00e-07 | 0.00e+00 | 1.53e-01 | 8.58e-01 | 0.00e+00 | 1.00e+00 | |
| sPYR | PYR.ms -> PYR.fix | 1.00e+00 | 1.00e+00 | 1.00e+00 | 1.00e+00 | 1.00e+00 | 1.00e+00 | |
| **EXCH** | | | | | | | | |
| **Transport** | | | | | | | | |
| MCT | LAC <-> LAC.x | 1.00e-07 | 0.00e+00 | 1.05e-01 | 1.52e+03 | 1.35e+03 | 2.41e+03 | 15200000000.00 |
| GLUR | GLU <-> GLU.x | 5.10e+00 | 4.77e+00 | 5.23e+00 | 1.54e+00 | 1.11e+00 | 2.54e+00 | 0.30 |
| **Glycolysis** | | | | | | | | |
| PGI | G6P <-> F6P | 2.78e+05 | 1.77e+05 | Inf | 2.46e+05 | 0.00e+00 | Inf | |
| ALDO | FBP <-> DHAP + GAP | 1.43e+02 | 1.43e+02 | 1.43e+02 | 3.20e+02 | 2.79e+02 | 3.60e+02 | 2.24 |
| TPI | DHAP <-> GAP | 4.33e+03 | 4.33e+03 | 1.09e+04 | 1.70e+03 | 1.06e+03 | 3.06e+03 | 0.39 |
| GAPDH | GAP <-> 3PG | 4.42e+02 | 4.72e+00 | 4.50e+02 | 1.00e-07 | 0.00e+00 | 2.39e+02 | |
| LDH | PYR.c <-> LAC | 1.63e+03 | 1.62e+03 | 1.80e+03 | 4.80e+00 | 0.00e+00 | 3.51e+02 | 0.00 |
| GPT1 | PYR.c <-> ALA | 1.00e-07 | 0.00e+00 | 2.61e-01 | 8.32e+02 | 0.00e+00 | 9.06e+02 | |
| GPT2 | PYR.m <-> ALA | 4.21e-04 | 0.00e+00 | 2.92e+00 | 1.28e-04 | 0.00e+00 | | |
| **Pentose phosphate pathway** | | | | | | | | |
| TK1 | P5P + P5P <-> S7P + GAP | 9.97e+04 | 6.27e+03 | Inf | 1.47e+02 | 6.67e+01 | 2.60e+02 | 0.00 |
| TA | S7P + GAP <-> F6P + E4P | 5.93e+00 | 5.79e+00 | 6.97e+00 | 2.35e-04 | 0.00e+00 | 7.54e+00 | |
| TK2 | P5P + E4P <-> F6P + GAP | 1.00e+07 | -Inf | Inf | 9.05e+00 | 4.10e+00 | 1.43e+01 | |
| **Anaplerosis** | | | | | | | | |

| | | | 21%[a] | | | 0.5%[b] | | |
|---|---|---|---|---|---|---|---|---|
| **ID** | **Reaction** | **Flux** | **LB** | **UB** | **Flux** | **LB** | **UB** | **Ratio** |
| GLDH | GLU <-> AKG | 1.52e+03 | 1.52e+03 | 7.13e+03 | 3.78e+02 | 1.93e+02 | 1.94e+03 | |
| GLS | GLN <-> GLU | 3.99e-01 | 0.00e+00 | 8.04e-01 | 1.00e-07 | 0.00e+00 | 3.84e-01 | |
| **Tricarboxylic acid cycle** | | | | | | | | |
| IDH | CIT <-> AKG + CO2 | 4.55e+00 | 4.03e+00 | 5.19e+00 | 2.52e+00 | 1.80e+00 | 4.50e+00 | |
| SDH | SUC <-> FUM | 1.22e+03 | | Inf | 7.60e+01 | 2.57e+01 | Inf | |
| FH | FUM <-> MAL | 3.66e+05 | 1.95e+05 | Inf | 5.05e+05 | 3.06e+02 | Inf | |
| MDH | MAL <-> OAC | 1.11e+03 | 7.88e+02 | 2.38e+03 | 1.33e+02 | 7.22e+01 | 3.25e+02 | 0.12 |
| GOT | OAC <-> ASP | 1.00e+07 | -Inf | Inf | 4.42e+01 | 0.00e+00 | Inf | |
| **Amino acid metabolism** | | | | | | | | |
| SHT | SER <-> GLY + MEETHF | 5.10e+00 | 8.92e-01 | 5.25e+00 | 6.07e-07 | 0.00e+00 | 3.32e+02 | |
| CYST | SER <-> CYS | 1.52e-05 | 0.00e+00 | 2.55e-04 | 1.46e-02 | 0.00e+00 | Inf | |

[a] SSR 391.7 [311.2-416.6] (95% CI, 362 DOF)

[b] SSR 334.3 [311.2-416.6] (95% CI, 362 DOF)

#### Table 2: Lung fibroblast fluxes following DMSO and BAY treatment

| | | DMSO[a] | | | BAY[b] | | | |
|---|---|---|---|---|---|---|---|---|
| **ID** | **Reaction** | **Flux** | **LB** | **UB** | **Flux** | **LB** | **UB** | **Ratio** |
| **NET** | | | | | | | | |
| **Transport** | | | | | | | | |
| GLUT | GLC.x -> GLC | 6.12e+02 | 6.12e+02 | 6.12e+02 | 8.80e+02 | 8.80e+02 | 8.80e+02 | 1.44 |
| PYRR | PYR.x -> PYR.c | 9.98e+01 | 9.95e+01 | 1.01e+02 | 6.06e+01 | 6.06e+01 | 6.06e+01 | 0.61 |
| MCT | LAC <-> LAC.x | 8.19e+02 | 8.17e+02 | 8.20e+02 | 1.33e+03 | 1.33e+03 | 1.33e+03 | 1.62 |
| ALAR | ALA -> ALA.x | 2.67e+00 | 2.36e+00 | 3.29e+00 | 5.98e+00 | 5.88e+00 | 6.24e+00 | 2.24 |
| GLNR | GLN.x -> GLN | 3.78e+01 | 3.77e+01 | 3.79e+01 | 2.06e+01 | 2.06e+01 | 2.06e+01 | 0.54 |
| GLUR | GLU <-> GLU.x | 1.61e+01 | 1.56e+01 | 1.62e+01 | 1.68e+01 | 1.68e+01 | 1.68e+01 | 1.05 |
| ASPR | ASP -> ASP.x | 2.36e+00 | 2.32e+00 | 2.49e+00 | 1.80e+00 | 1.80e+00 | 1.81e+00 | 0.76 |
| SERR | SER.x -> SER | 1.03e+01 | 1.03e+01 | 1.06e+01 | 2.50e+00 | 2.50e+00 | 2.50e+00 | 0.24 |
| CYSR | CYX.x -> CYS + CYS | 2.79e+00 | 2.79e+00 | 2.95e+00 | 3.07e-01 | 3.06e-01 | 3.07e-01 | 0.11 |
| GLYR | GLY -> GLY.x | 2.52e+00 | 2.30e+00 | 2.73e+00 | 5.52e-01 | 4.30e-01 | 7.45e-01 | 0.22 |
| **Glycolysis** | | | | | | | | |
| HK | GLC -> G6P | 6.12e+02 | 6.12e+02 | 6.12e+02 | 8.80e+02 | 8.80e+02 | 8.80e+02 | 1.44 |
| PGI | G6P <-> F6P | 6.09e+02 | 6.08e+02 | 6.09e+02 | 8.42e+02 | 8.42e+02 | 8.42e+02 | 1.38 |
| PFK | F6P -> FBP | 6.07e+02 | 6.07e+02 | 6.07e+02 | 8.65e+02 | 8.65e+02 | 8.65e+02 | 1.43 |
| ALDO | FBP <-> DHAP + GAP | 6.07e+02 | 6.07e+02 | 6.07e+02 | 8.65e+02 | 8.65e+02 | 8.65e+02 | 1.43 |
| TPI | DHAP <-> GAP | 6.06e+02 | 6.06e+02 | 6.06e+02 | 8.65e+02 | 8.65e+02 | 8.65e+02 | 1.43 |
| GAPDH | GAP <-> 3PG | 1.21e+03 | 1.21e+03 | 1.21e+03 | 1.74e+03 | 1.74e+03 | 1.74e+03 | 1.44 |
| ENO | 3PG -> PEP | 1.21e+03 | 1.21e+03 | 1.21e+03 | 1.57e+03 | 1.57e+03 | 1.57e+03 | 1.30 |
| PK | PEP -> PYR.c | 1.23e+03 | 1.19e+03 | 1.23e+03 | 1.65e+03 | 1.65e+03 | 1.65e+03 | 1.34 |
| LDH | PYR.c <-> LAC | 8.19e+02 | 8.17e+02 | 8.20e+02 | 1.33e+03 | 1.33e+03 | 1.33e+03 | 1.62 |
| GPT1 | PYR.c <-> ALA | 9.62e+00 | 9.44e+00 | 9.62e+00 | 9.36e+00 | 9.32e+00 | 9.42e+00 | 0.97 |
| GPT2 | PYR.m <-> ALA | 1.14e-01 | | | 2.28e-07 | -1.22e-05 | 6.41e-04 | |
| **Pentose phosphate pathway** | | | | | | | | |
| G6PD | G6P -> P5P + CO2 | 2.02e-02 | 0.00e+00 | 1.08e+00 | 3.64e+01 | 3.64e+01 | 3.64e+01 | 1801.98 |
| TK1 | P5P + P5P <-> S7P + GAP | -9.06e-01 | -9.28e-01 | -9.06e-01 | 1.17e+01 | 1.17e+01 | 1.17e+01 | -12.89 |

| | | DMSO[a] | | | BAY[b] | | | |
|---|---|---|---|---|---|---|---|---|
| **ID** | **Reaction** | **Flux** | **LB** | **UB** | **Flux** | **LB** | **UB** | **Ratio** |
| TA | S7P + GAP <-> F6P + E4P | -9.06e-01 | -9.28e-01 | -9.06e-01 | 1.17e+01 | 1.17e+01 | 1.17e+01 | -12.89 |
| TK2 | P5P + E4P <-> F6P + GAP | -9.06e-01 | -9.28e-01 | -9.06e-01 | 1.17e+01 | 1.17e+01 | 1.17e+01 | -12.89 |
| **Anaplerosis** | | | | | | | | |
| PYRT | PYR.c -> PYR.m | 4.99e+02 | 4.97e+02 | 4.99e+02 | 5.50e+02 | 5.50e+02 | 5.50e+02 | 1.10 |
| PC | PYR.m + CO2 -> OAC | 2.11e+01 | 2.07e+01 | 2.17e+01 | 9.05e+01 | 9.05e+01 | 9.05e+01 | 4.28 |
| PEPCK | OAC -> PEP + CO2 | 1.36e+01 | 1.36e+01 | 1.37e+01 | 8.58e+01 | 8.58e+01 | 8.58e+01 | 6.31 |
| ME2 | MAL -> PYR.m + CO2 | 1.30e+01 | 1.28e+01 | 1.37e+01 | 1.00e-07 | 0.00e+00 | 9.49e-06 | 0.00 |
| ME1 | MAL -> PYR.c + CO2 | 3.20e-03 | 0.00e+00 | 1.73e+00 | 1.00e-07 | 0.00e+00 | 2.15e-05 | |
| FAO | FAO -> AcCoA.m | 1.00e-07 | 0.00e+00 | 3.48e+00 | 1.09e-04 | 8.34e-06 | 4.14e-02 | |
| GLDH | GLU <-> AKG | 1.33e+01 | 1.31e+01 | 1.35e+01 | -2.46e-01 | -2.47e-01 | -2.46e-01 | -0.02 |
| GLS | GLN <-> GLU | 3.40e+01 | 3.35e+01 | 3.42e+01 | 1.88e+01 | 1.88e+01 | 1.88e+01 | 0.55 |
| **Tricarboxylic acid cycle** | | | | | | | | |
| PDH | PYR.m -> AcCoA.m + CO2 | 4.90e+02 | 4.90e+02 | 4.92e+02 | 4.60e+02 | 4.60e+02 | 4.60e+02 | 0.94 |
| CS | AcCoA.m + OAC -> CIT | 4.90e+02 | 4.84e+02 | 4.91e+02 | 4.60e+02 | 4.60e+02 | 4.60e+02 | 0.94 |
| IDH | CIT <-> AKG + CO2 | 2.70e+01 | 2.70e+01 | 2.76e+01 | 1.45e+01 | 1.45e+01 | 1.45e+01 | 0.54 |
| OGDH | AKG -> SUC + CO2 | 4.03e+01 | 3.99e+01 | 4.04e+01 | 1.43e+01 | 1.43e+01 | 1.43e+01 | 0.35 |
| SDH | SUC <-> FUM | 4.03e+01 | 3.99e+01 | 4.04e+01 | 1.43e+01 | 1.43e+01 | 1.43e+01 | 0.35 |
| FH | FUM <-> MAL | 4.03e+01 | 3.99e+01 | 4.04e+01 | 1.43e+01 | 1.43e+01 | 1.43e+01 | 0.35 |
| MDH | MAL <-> OAC | 4.91e+02 | 4.91e+02 | 4.92e+02 | 4.60e+02 | 4.60e+02 | 4.60e+02 | 0.94 |
| GOT | OAC <-> ASP | 7.91e+00 | 7.76e+00 | 7.98e+00 | 4.46e+00 | 4.46e+00 | 4.46e+00 | 0.56 |
| **Amino acid metabolism** | | | | | | | | |
| PST | 3PG -> SER | 4.03e-01 | 3.74e-01 | 5.04e-01 | 1.73e+02 | 1.73e+02 | 1.73e+02 | 429.83 |
| SHT | SER <-> GLY + MEETHF | 6.63e+00 | 6.59e+00 | 6.65e+00 | 2.85e+00 | 2.79e+00 | 2.93e+00 | 0.43 |
| CYST | SER <-> CYS | -3.88e+00 | -3.91e+00 | -3.87e+00 | 2.03e-01 | 2.02e-01 | 2.03e-01 | -0.05 |
| SD | SER -> PYR.c | 2.80e+00 | 2.80e+00 | 2.80e+00 | 1.70e+02 | 1.70e+02 | 1.70e+02 | 60.81 |
| GLYS | CO2 + MEETHF -> GLY | 3.63e+00 | 3.50e+00 | 3.65e+00 | 1.41e+00 | 1.30e+00 | 1.46e+00 | 0.39 |
| **Biomass** | | | | | | | | |

| | | DMSO[a] | | | BAY[b] | | | |
|---|---|---|---|---|---|---|---|---|
| ID | Reaction | Flux | LB | UB | Flux | LB | UB | Ratio |
| BIOMASS | 1216*AcCoA.c + 295.6*ALA + 232.4*ASP + 114.7*CO2 + 71.43*CYS + 57.14*DHAP + 142.4*G6P + 158.6*GLN + 190.1*GLU + 324.2*GLY + 125.6*MEETHF + 114.7*P5P + 217.2*SER -> biomass | 2.39e-02 | 2.39e-02 | 2.50e-02 | 1.14e-02 | 1.14e-02 | 1.14e-02 | 0.48 |
| ACL | CIT -> AcCoA.c + MAL | 4.63e+02 | 4.63e+02 | 4.66e+02 | 4.45e+02 | 4.45e+02 | 4.45e+02 | 0.96 |
| LIPS | AcCoA.c -> lipid | 4.34e+02 | 4.29e+02 | 4.34e+02 | 4.32e+02 | 4.32e+02 | 4.32e+02 | |
| **Mixing** | | | | | | | | |
| cPYR | 0*PYR.c -> PYR.ms | 1.00e+00 | 9.99e-01 | 1.00e+00 | 1.00e-07 | 0.00e+00 | 1.00e+00 | |
| mPYR | 0*PYR.m -> PYR.ms | 1.00e-07 | 0.00e+00 | 9.83e-04 | 1.00e+00 | 0.00e+00 | 1.00e+00 | |
| sPYR | PYR.ms -> PYR.fix | 1.00e+00 | 1.00e+00 | 1.00e+00 | 1.00e+00 | 1.00e+00 | 1.00e+00 | |
| **EXCH** | | | | | | | | |
| **Transport** | | | | | | | | |
| MCT | LAC <-> LAC.x | 6.24e-04 | 0.00e+00 | 3.56e+00 | 7.11e+02 | 7.11e+02 | 7.11e+02 | 1139423.08 |
| GLUR | GLU <-> GLU.x | 5.06e+00 | 4.82e+00 | 5.75e+00 | 3.48e+00 | 3.48e+00 | 3.48e+00 | 0.69 |
| **Glycolysis** | | | | | | | | |
| PGI | G6P <-> F6P | 1.40e+06 | 1.39e+06 | Inf | 4.31e+06 | 4.31e+06 | 4.31e+06 | |
| ALDO | FBP <-> DHAP + GAP | 2.38e+02 | 2.38e+02 | 2.38e+02 | 1.02e+03 | 1.02e+03 | 1.02e+03 | 4.28 |
| TPI | DHAP <-> GAP | 9.99e+06 | | Inf | 7.57e+03 | 7.57e+03 | 7.57e+03 | |
| GAPDH | GAP <-> 3PG | 5.81e+02 | 5.81e+02 | 7.25e+02 | 1.09e+02 | 1.07e+02 | 1.09e+02 | 0.19 |
| LDH | PYR.c <-> LAC | 2.65e+03 | 2.58e+03 | 2.65e+03 | 4.92e+01 | 4.91e+01 | 4.94e+01 | 0.02 |
| GPT1 | PYR.c <-> ALA | 1.00e-07 | 0.00e+00 | 5.60e-02 | 2.45e+03 | 2.45e+03 | 2.45e+03 | 24500000000.00 |
| GPT2 | PYR.m <-> ALA | 1.00e-07 | 0.00e+00 | 5.65e-02 | 1.00e-07 | 0.00e+00 | 1.20e-05 | |
| **Pentose phosphate pathway** | | | | | | | | |
| TK1 | P5P + P5P <-> S7P + GAP | 1.28e+06 | 9.01e+03 | Inf | 1.00e+07 | -Inf | Inf | |
| TA | S7P + GAP <-> F6P + E4P | 8.89e+00 | 8.88e+00 | 9.53e+00 | 5.10e+01 | 5.10e+01 | 5.10e+01 | 5.74 |
| TK2 | P5P + E4P <-> F6P + GAP | 6.93e+00 | 5.12e+00 | 6.98e+00 | 1.00e-07 | 0.00e+00 | 1.56e-04 | 0.00 |
| **Anaplerosis** | | | | | | | | |
| GLDH | GLU <-> AKG | 5.63e+03 | 4.43e+03 | 5.66e+03 | 1.42e+03 | 1.42e+03 | 1.42e+03 | 0.25 |
| GLS | GLN <-> GLU | 1.27e+00 | 1.20e+00 | 1.50e+00 | 5.52e-01 | 5.51e-01 | 5.55e-01 | 0.43 |

| | | DMSO[a] | | | BAY[b] | | | |
|---|---|---|---|---|---|---|---|---|
| ID | Reaction | Flux | LB | UB | Flux | LB | UB | Ratio |
| **Tricarboxylic acid cycle** | | | | | | | | |
| IDH | CIT <-> AKG + CO2 | 3.36e+00 | 3.24e+00 | 3.92e+00 | 4.66e+00 | 4.66e+00 | 4.66e+00 | 1.39 |
| SDH | SUC <-> FUM | 4.30e+02 | 4.30e+02 | 1.46e+06 | 1.04e+04 | 1.04e+04 | 1.04e+04 | |
| FH | FUM <-> MAL | 7.29e+06 | -Inf | Inf | 4.56e+06 | 4.56e+06 | 4.56e+06 | |
| MDH | MAL <-> OAC | 5.49e+02 | 5.47e+02 | 5.49e+02 | 1.00e-07 | 0.00e+00 | 6.30e-03 | 0.00 |
| GOT | OAC <-> ASP | 1.04e+02 | 1.04e+02 | 1.04e+02 | 4.76e+05 | 4.76e+05 | 4.76e+05 | 4576.92 |
| **Amino acid metabolism** | | | | | | | | |
| SHT | SER <-> GLY + MEETHF | 1.39e+00 | 1.37e+00 | 1.41e+00 | 1.86e+03 | 1.86e+03 | 1.86e+03 | 1338.13 |
| CYST | SER <-> CYS | 1.25e-07 | 0.00e+00 | 4.22e-02 | 1.33e-01 | 1.33e-01 | 1.33e-01 | 1064000.00 |

[a] SSR 393.5 [311.2-416.6] (95% CI, 362 DOF)

[b] SSR 392.4 [308.4-413.4] (95% CI, 359 DOF)

#### Table 3: PASMC fluxes in 21% and 0.5% oxygen

| | | 21%[a] | | | 0.5%[b] | | | |
|---|---|---|---|---|---|---|---|---|
| **ID** | **Reaction** | **Flux** | **LB** | **UB** | **Flux** | **LB** | **UB** | **Ratio** |
| **NET** | | | | | | | | |
| **Transport** | | | | | | | | |
| GLUT | GLC.x -> GLC | 4.28e+02 | 4.28e+02 | 4.28e+02 | 3.65e+02 | 3.65e+02 | 3.65e+02 | 0.85 |
| PYRR | PYR.x -> PYR.c | 1.04e+02 | 1.02e+02 | 1.09e+02 | 4.53e+01 | 4.31e+01 | 4.57e+01 | 0.44 |
| MCT | LAC <-> LAC.x | 8.01e+02 | 8.01e+02 | 8.04e+02 | 6.49e+02 | 6.49e+02 | 6.49e+02 | 0.81 |
| ALAR | ALA -> ALA.x | 1.43e+01 | 1.43e+01 | 1.46e+01 | 7.83e+00 | 7.83e+00 | 8.24e+00 | 0.55 |
| GLNR | GLN.x -> GLN | 7.73e+01 | 7.53e+01 | 7.73e+01 | 1.77e+02 | 1.77e+02 | 1.77e+02 | 2.29 |
| GLUR | GLU <-> GLU.x | 2.53e+01 | 2.52e+01 | 2.54e+01 | 1.19e+01 | 1.19e+01 | 1.22e+01 | 0.47 |
| ASPR | ASP -> ASP.x | 7.01e+00 | 6.99e+00 | 7.02e+00 | 6.92e+00 | 6.84e+00 | 7.00e+00 | |
| SERR | SER.x -> SER | 2.54e+00 | 2.48e+00 | 2.55e+00 | 2.57e+00 | 2.55e+00 | 2.57e+00 | 1.01 |
| CYSR | CYX.x -> CYS + CYS | 6.39e+00 | 6.34e+00 | 6.45e+00 | 3.75e+00 | 3.75e+00 | 3.75e+00 | 0.59 |
| GLYR | GLY -> GLY.x | 3.66e-01 | 3.03e-01 | 4.19e-01 | 4.06e-01 | 3.86e-01 | 4.25e-01 | |
| **Glycolysis** | | | | | | | | |
| HK | GLC -> G6P | 4.28e+02 | 4.28e+02 | 4.28e+02 | 3.65e+02 | 3.65e+02 | 3.65e+02 | 0.85 |
| PGI | G6P <-> F6P | 4.06e+02 | 4.06e+02 | 4.07e+02 | 3.62e+02 | 3.62e+02 | 3.63e+02 | 0.89 |
| PFK | F6P -> FBP | 4.17e+02 | 4.17e+02 | 4.18e+02 | 3.61e+02 | 3.60e+02 | 3.61e+02 | 0.87 |
| ALDO | FBP <-> DHAP + GAP | 4.17e+02 | 4.17e+02 | 4.18e+02 | 3.61e+02 | 3.60e+02 | 3.61e+02 | 0.87 |
| TPI | DHAP <-> GAP | 4.16e+02 | 4.16e+02 | 4.16e+02 | 3.60e+02 | 3.60e+02 | 3.60e+02 | 0.87 |
| GAPDH | GAP <-> 3PG | 8.39e+02 | 8.39e+02 | 8.41e+02 | 7.21e+02 | 7.21e+02 | 7.21e+02 | 0.86 |
| ENO | 3PG -> PEP | 8.36e+02 | 8.35e+02 | 8.53e+02 | 7.20e+02 | 7.20e+02 | 7.20e+02 | 0.86 |
| PK | PEP -> PYR.c | 9.31e+02 | 9.30e+02 | 9.31e+02 | 9.24e+02 | 9.24e+02 | 9.24e+02 | 0.99 |
| LDH | PYR.c <-> LAC | 8.01e+02 | 8.01e+02 | 8.04e+02 | 6.49e+02 | 6.49e+02 | 6.49e+02 | 0.81 |
| GPT1 | PYR.c <-> ALA | 1.64e+02 | 1.62e+02 | 1.92e+02 | -1.36e+01 | -1.39e+01 | -1.35e+01 | -0.08 |
| GPT2 | PYR.m <-> ALA | -1.43e+02 | -1.43e+02 | -1.42e+02 | 2.62e+01 | 2.51e+01 | 2.65e+01 | -0.18 |
| **Pentose phosphate pathway** | | | | | | | | |
| G6PD | G6P -> P5P + CO2 | 1.89e+01 | 1.57e+01 | 1.93e+01 | 1.16e-07 | 0.00e+00 | 1.10e-03 | 0.00 |
| TK1 | P5P + P5P <-> S7P + GAP | 5.46e+00 | 4.44e+00 | 5.96e+00 | -6.15e-01 | -6.15e-01 | -5.77e-01 | -0.11 |

| | | 21%[a] | | | 0.5%[b] | | | |
|---|---|---|---|---|---|---|---|---|
| ID | Reaction | Flux | LB | UB | Flux | LB | UB | Ratio |
| TA | S7P + GAP <-> F6P + E4P | 5.46e+00 | 4.44e+00 | 5.96e+00 | -6.15e-01 | -6.15e-01 | -5.77e-01 | -0.11 |
| TK2 | P5P + E4P <-> F6P + GAP | 5.46e+00 | 4.44e+00 | 5.96e+00 | -6.15e-01 | -6.15e-01 | -5.77e-01 | -0.11 |
| **Anaplerosis** | | | | | | | | |
| PYRT | PYR.c -> PYR.m | 7.60e+01 | 7.59e+01 | 7.66e+01 | 3.36e+02 | 3.36e+02 | 3.36e+02 | 4.42 |
| PC | PYR.m + CO2 -> OAC | 6.30e+01 | 6.29e+01 | 6.59e+01 | 2.37e+02 | 2.36e+02 | 2.37e+02 | 3.76 |
| PEPCK | OAC -> PEP + CO2 | 9.51e+01 | 9.51e+01 | 9.53e+01 | 2.03e+02 | 2.03e+02 | 2.04e+02 | 2.14 |
| ME2 | MAL -> PYR.m + CO2 | 1.20e-03 | 0.00e+00 | 5.20e-03 | 1.82e+02 | 1.81e+02 | 1.82e+02 | 151517.08 |
| ME1 | MAL -> PYR.c + CO2 | 3.29e-05 | 0.00e+00 | 1.15e+00 | 5.91e-05 | 0.00e+00 | 8.06e-02 | |
| FAO | FAO -> AcCoA.m | 1.00e-07 | 0.00e+00 | 1.32e-02 | 1.15e-04 | 0.00e+00 | 1.56e-01 | |
| GLDH | GLU <-> AKG | 4.43e+01 | 4.42e+01 | 4.45e+01 | 1.59e+02 | 1.59e+02 | 1.59e+02 | 3.60 |
| GLS | GLN <-> GLU | 7.38e+01 | 7.36e+01 | 7.38e+01 | 1.74e+02 | 1.74e+02 | 1.74e+02 | 2.36 |
| **Tricarboxylic acid cycle** | | | | | | | | |
| PDH | PYR.m -> AcCoA.m + CO2 | 1.56e+02 | 1.48e+02 | 1.66e+02 | 2.55e+02 | 2.55e+02 | 2.55e+02 | 1.63 |
| CS | AcCoA.m + OAC -> CIT | 1.56e+02 | 1.56e+02 | 1.58e+02 | 2.55e+02 | 2.55e+02 | 2.55e+02 | 1.63 |
| IDH | CIT <-> AKG + CO2 | 2.11e+01 | 2.10e+01 | 2.11e+01 | 2.16e+01 | 2.16e+01 | 2.16e+01 | 1.03 |
| OGDH | AKG -> SUC + CO2 | 6.54e+01 | 6.51e+01 | 6.59e+01 | 1.81e+02 | 1.80e+02 | 1.81e+02 | 2.77 |
| SDH | SUC <-> FUM | 6.54e+01 | 6.51e+01 | 6.59e+01 | 1.81e+02 | 1.80e+02 | 1.81e+02 | 2.77 |
| FH | FUM <-> MAL | 6.54e+01 | 6.51e+01 | 6.59e+01 | 1.81e+02 | 1.80e+02 | 1.81e+02 | 2.77 |
| MDH | MAL <-> OAC | 2.01e+02 | 2.01e+02 | 2.01e+02 | 2.32e+02 | 2.32e+02 | 2.33e+02 | 1.16 |
| GOT | OAC <-> ASP | 1.22e+01 | 1.17e+01 | 1.24e+01 | 1.07e+01 | 1.06e+01 | 1.07e+01 | 0.87 |
| **Amino acid metabolism** | | | | | | | | |
| PST | 3PG -> SER | 2.69e+00 | 2.57e+00 | 2.80e+00 | 7.12e-01 | 7.01e-01 | 7.21e-01 | 0.26 |
| SHT | SER <-> GLY + MEETHF | 5.19e+00 | 5.15e+00 | 5.20e+00 | 3.82e+00 | 3.81e+00 | 3.86e+00 | 0.74 |
| CYST | SER <-> CYS | -1.12e+01 | -1.17e+01 | -1.11e+01 | -6.35e+00 | -6.35e+00 | -6.35e+00 | 0.57 |
| SD | SER -> PYR.c | 6.39e+00 | 6.23e+00 | 6.44e+00 | 2.33e+00 | 2.33e+00 | 2.33e+00 | 0.36 |
| GLYS | CO2 + MEETHF -> GLY | 2.39e+00 | 2.36e+00 | 2.42e+00 | 1.80e+00 | 1.79e+00 | 1.81e+00 | 0.75 |
| **Biomass** | | | | | | | | |

| | | 21%[a] | | | 0.5%[b] | | | |
|---|---|---|---|---|---|---|---|---|
| ID | Reaction | Flux | LB | UB | Flux | LB | UB | Ratio |
| BIOMASS | 978*AcCoA.c + 237.8*ALA + 187*ASP + 92.3*CO2 + 57.46*CYS + 45.97*DHAP + 114.5*G6P + 127.6*GLN + 153*GLU + 260.8*GLY + 101.1*MEETHF + 92.3*P5P + 174.8*SER -> biomass | 2.77e-02 | 2.70e-02 | 2.79e-02 | 2.00e-02 | 2.00e-02 | 2.00e-02 | 0.72 |
| ACL | CIT -> AcCoA.c + MAL | 1.35e+02 | 1.34e+02 | 1.38e+02 | 2.33e+02 | 2.33e+02 | 2.33e+02 | 1.72 |
| LIPS | AcCoA.c -> lipid | 1.08e+02 | 9.99e+01 | 1.08e+02 | 2.14e+02 | 2.14e+02 | 2.14e+02 | 1.98 |
| **Mixing** | | | | | | | | |
| cPYR | 0*PYR.c -> PYR.ms | 5.77e-01 | 5.64e-01 | 5.92e-01 | 1.00e+00 | 9.96e-01 | 1.00e+00 | 1.73 |
| mPYR | 0*PYR.m -> PYR.ms | 4.23e-01 | 4.08e-01 | 4.36e-01 | 1.00e-07 | 0.00e+00 | 4.40e-03 | 0.00 |
| sPYR | PYR.ms -> PYR.fix | 1.00e+00 | 1.00e+00 | 1.00e+00 | 1.00e+00 | 1.00e+00 | 1.00e+00 | |
| **EXCH** | | | | | | | | |
| **Transport** | | | | | | | | |
| MCT | LAC <-> LAC.x | 1.00e-07 | 0.00e+00 | 1.36e+02 | 1.64e+03 | 1.63e+03 | 1.65e+03 | 16400000000.00 |
| GLUR | GLU <-> GLU.x | 1.00e-07 | 0.00e+00 | 2.27e-02 | 5.69e-05 | 0.00e+00 | 1.71e-02 | |
| **Glycolysis** | | | | | | | | |
| PGI | G6P <-> F6P | 4.88e+06 | 4.88e+06 | Inf | 9.92e+06 | 9.85e+04 | Inf | |
| ALDO | FBP <-> DHAP + GAP | 2.89e+02 | 2.80e+02 | 2.89e+02 | 2.57e+02 | 2.56e+02 | 2.57e+02 | 0.89 |
| TPI | DHAP <-> GAP | 9.86e+06 | -Inf | Inf | 1.65e+03 | 1.63e+03 | 1.68e+03 | |
| GAPDH | GAP <-> 3PG | 1.12e+03 | 0.00e+00 | 5.88e+05 | 1.00e-07 | 0.00e+00 | 2.27e-01 | |
| LDH | PYR.c <-> LAC | 1.47e+03 | 1.39e+03 | 1.47e+03 | 4.49e+02 | 4.49e+02 | 4.49e+02 | 0.31 |
| GPT1 | PYR.c <-> ALA | 2.74e+02 | 2.73e+02 | 2.77e+02 | 1.00e-07 | 0.00e+00 | 4.28e-02 | 0.00 |
| GPT2 | PYR.m <-> ALA | 1.38e+02 | 1.38e+02 | 1.49e+02 | 9.64e+01 | 0.00e+00 | 1.01e+02 | 0.70 |
| **Pentose phosphate pathway** | | | | | | | | |
| TK1 | P5P + P5P <-> S7P + GAP | 7.99e+02 | 7.97e+02 | 8.08e+02 | 3.54e+01 | 3.54e+01 | 3.55e+01 | 0.04 |
| TA | S7P + GAP <-> F6P + E4P | 1.53e-01 | 0.00e+00 | 5.82e-01 | 2.55e+00 | 2.54e+00 | 2.57e+00 | 16.67 |
| TK2 | P5P + E4P <-> F6P + GAP | 3.33e+00 | 2.62e+00 | 3.35e+00 | 1.29e+01 | 1.29e+01 | 1.29e+01 | 3.88 |
| **Anaplerosis** | | | | | | | | |
| GLDH | GLU <-> AKG | 5.36e+02 | 5.34e+02 | 8.37e+02 | 1.23e+03 | 1.23e+03 | 1.23e+03 | 2.29 |
| GLS | GLN <-> GLU | 3.20e-01 | 0.00e+00 | 2.74e+00 | 1.12e+00 | 1.07e+00 | 1.74e+00 | |
| **Tricarboxylic acid cycle** | | | | | | | | |

| | | 21%[a] | | | 0.5%[b] | | | |
|---|---|---|---|---|---|---|---|---|
| **ID** | **Reaction** | **Flux** | **LB** | **UB** | **Flux** | **LB** | **UB** | **Ratio** |
| IDH | CIT <-> AKG + CO2 | 1.04e+01 | 1.02e+01 | 1.04e+01 | 6.30e+01 | 6.30e+01 | 6.31e+01 | 6.09 |
| SDH | SUC <-> FUM | 2.78e-01 | 0.00e+00 | Inf | 3.34e+06 | 3.34e+06 | 3.34e+06 | |
| FH | FUM <-> MAL | 1.03e-04 | 0.00e+00 | 1.58e+01 | 2.18e+02 | 2.18e+02 | 2.18e+02 | 2114238.83 |
| MDH | MAL <-> OAC | 1.01e+03 | 8.27e+02 | 1.01e+03 | 3.67e+03 | 3.67e+03 | 3.69e+03 | 3.63 |
| GOT | OAC <-> ASP | 2.27e+02 | 2.27e+02 | 2.47e+02 | 1.54e+01 | 1.54e+01 | 1.55e+01 | 0.07 |
| **Amino acid metabolism** | | | | | | | | |
| SHT | SER <-> GLY + MEETHF | 3.55e+00 | 3.52e+00 | 3.59e+00 | 1.60e-01 | 1.36e-01 | 1.70e-01 | 0.05 |
| CYST | SER <-> CYS | 1.04e+03 | 1.03e+03 | 1.04e+03 | 2.00e-03 | 0.00e+00 | 2.00e-03 | 0.00 |

[a] SSR 575.6 [499.1-630.6] (95% CI, 563 DOF)

[b] SSR 521.3 [482.2-611.6] (95% CI, 545 DOF)

**FIGURE LEGENDS**

Supplementary Figure 1: **Supporting data for extracellular flux calculations.** (**A**) Cell viability as assessed by live/dead cell staining with acridine orange and propidium iodide did not differ between 21% and 0.5% oxygen culture conditions (n = 3 technical replicates). (**B**) Standard curve of lung fibroblast (LF) cell count *v.* total DNA by PicoGreen measurement used to interpolate cell numbers from DNA measurements. Data are mean ± SEM of three biological replicates. (**C**) Standard curve of PASMC cell count *v.* total DNA as in (B). (**D**) Total DNA measurements were compared to direct cell counts over the experimental time course. Cell counts and total DNA were obtained from the same sample wells. The slopes of the best-fit lines for 21% (*red*) and 0.5% (*blue*) samples were not statistically different. (**E**) Predicted well volumes were estimated from the change in culture plate mass over the experimental time course. Evaporation rates were different depending on the culture conditions and treatment. Although the mean evaporation rate is depicted, experiment-specific evaporation rates were used to calculate fluxes for each biological replicate (**F**) Metabolite accumulation (positive values) and degradation (negative values) rates. Data are mean ± SEM of 3-8 biological replicates. Rates significantly different from 0 (*) based on a probability value < 0.05 using Student's one-sample *t*-test were incorporated into flux calculations.

Supplementary Figure 2: **Effects of hypoxia on extracellular metabolite fluxes in lung fibroblasts.** (**A**) Lung fibroblasts (LFs) were cultured in 21% or 0.2% oxygen beginning 24 h prior to time 0. Samples were collected every 24 h for 72 h. (**B**) Growth curves of LFs in each experimental condition (n = 4). (**C**) Growth rates from (B) were determined by robust linear modeling of log-transformed growth curves. (**D**) Representative immunoblot of LF protein lysates cultured as in (A). (**E**) Relative change in HIF-1α protein levels from (D) normalized to 21% oxygen at time 0 (n = 4). (**F**) Relative change in GLUT1 mRNA levels normalized to 21% oxygen treatment at time 0 (n = 4). (**G**) Relative change in LDHA mRNA levels as in (F). (**H**) Relative change in LDHA protein levels as in (E). (**I**) Extracellular fluxes of glucose (GLC) and lactate (LAC) (n = 4). By convention, negative fluxes indicate metabolite consumption. (**J**) Extracellular fluxes of pyruvate (PYR) and amino acids. Data are mean ± SEM (* p < 0.05).

Supplementary Figure 3: **Effects of hypoxia on extracellular metabolite fluxes in pulmonary artery smooth muscle cells.** (**A**) Pulmonary artery smooth muscle cells (PASMCs) were cultured in 21% or 0.5% oxygen beginning 24 h prior to time 0. Samples were collected every 12 h for 48 h. (**B**) Growth curves of LFs in each experimental condition (n = 4). (**C**) Growth rates from (B) were determined by robust linear modeling of log-transformed growth curves. (**D**) Representative immunoblot of LF protein lysates cultured as in (A). (**E**) Relative change in HIF-1α protein levels from (D) normalized to 21% oxygen at time 0 (n = 4). (**F**) Relative change in GLUT1 mRNA levels normalized to 21% oxygen treatment at time 0 (n = 4). (**G**) Relative change in LDHA mRNA levels as in (F). (**H**) Relative change in LDHA protein levels as in (E). (**I**) Extracellular fluxes of glucose (GLC) and lactate (LAC) (n = 4). By convention, negative fluxes indicate metabolite consumption. (**J**) Extracellular fluxes of pyruvate (PYR) and amino acids. Data are mean ± SEM (* $p < 0.05$).

Supplementary Figure 4: **Mass isotopomer distributions after 72 h of labeling in lung fibroblasts.** Lung fibroblasts (LFs) were labeled with the indicated tracers and intracellular metabolites were analyzed by LC-MS after 72 h. Mass isotopomer distributions were adjusted for natural abundance. Data are the mean ± SEM of 4 biological replicates. Significant differences in labeling patterns between 21% and 0.5% oxygen (*), DMSO and BAY treatment (†), and 0.5% oxygen and BAY treatment (‡) for each combination of metabolite and tracer are highlighted.

Supplementary Figure 5: **Mass isotopomer distributions after 48 h of labeling in pulmonary artery smooth muscle cells.** Pulmonary artery smooth muscle cells (PASMCs) were labeled with the indicated tracers and intracellular metabolites were analyzed by LC-MS after 36 h. Mass isotopomer distributions were adjusted for natural abundance. Data are the mean ± SEM of 4 biological replicates. Significant differences in labeling patterns between 21% and 0.5% oxygen (*) for each combination of metabolite and tracer are highlighted.

Supplementary Figure 6: **Isotope incorporation over the labeling time course.** LFs were cultured in 21% or 0.5% oxygen and labeled with the indicated tracers. Intracellular metabolites were analyzed by LC-MS (FBP, fructose-bisphosphate; PYR, pyruvate; CIT, citrate; MAL, malate). Mass isotopomer distributions were calculated and adjusted for natural abundance. Data show the total amount of metabolite labeling (*i.e.,* 1 - M0 fractional abundance). Data are the mean ± SEM of 4 biological replicates.

Supplementary Figure 7: **Isotopically non-stationary metabolic flux analysis.** (**A**) Metabolic flux model of LF metabolism in 21% oxygen. Arrows are colored by $\log_{10}$(flux). (**B**) Metabolic flux model of PASMC metabolism in 21% oxygen as in (A). (**C**) Ratio of metabolic fluxes in PASMCs compared to LFs. Fluxes with non-overlapping confidence intervals are highlighted with arrows colored according to the magnitude of the change. Arrow thickness corresponds to the absolute flux measured in LFs. (**D**) Ratio of metabolic fluxes in 0.5% oxygen compared to 21% oxygen in PASMCs. (**E**) LF fluxes were normalized to cell growth rate. Graph depicts the ratio of normalized metabolic fluxes in LFs cultured in 0.5% oxygen compared to 21% oxygen control. Fluxes with non-overlapping confidence intervals are highlighted to indicate significant changes.

Supplementary Figure 8: **Metabolomic profiling of hypoxia and BAY treated lung fibroblasts.** (**A**) Volcano plot of differentially regulated metabolites following 0.5% oxygen culture. (**B**) Volcano plot of differentially regulated metabolites by BAY treatment in 21% oxygen. (**C**) Venn diagram illustrating the overlap among metabolites differentially regulated by hypoxia (*blue*) or BAY treatment (*purple*). (**D**) Metabolite enrichment in KEGG pathways following hypoxia. (**E**) Metabolite enrichment in KEGG pathways following BAY. Only significantly enriched metabolite sets with $p < 0.05$ are shown. NES, normalized enrichment score.

Supplementary Figure 9: **Transcriptomic profiling of hypoxia and BAY treated lung fibroblasts.** (**A**) Volcano plot of differentially expressed genes following 0.5% oxygen culture. (**B**) Volcano plot of differentially expressed genes following BAY treatment. (**C**) Venn diagram illustrating the number of differentially expressed transcripts following hypoxia (*blue*) or BAY treatment (*purple*). (**D**) Venn diagram illustrating the number of differentially enriched Hallmark gene sets with hypoxia (*blue*) or BAY treatment (*purple*). (**E**) Gene set enrichment results of hypoxia-treated cells. (**F**) Gene set enrichment results of BAY-treated cells. All gene sets listed were significantly enriched at FDR < 0.05. NES, normalized enrichment score. (**G**) Volcano plot illustrating the results of a transcription factor enrichment analysis in hypoxia-treated cells. (**H**) Volcano plot illustrating the results of a transcription factor enrichment analysis in BAY-treated cells. (**I**) Venn diagram illustrating the overlap among enriched transcription factors following hypoxia or BAY treatment.

#### FIGURES

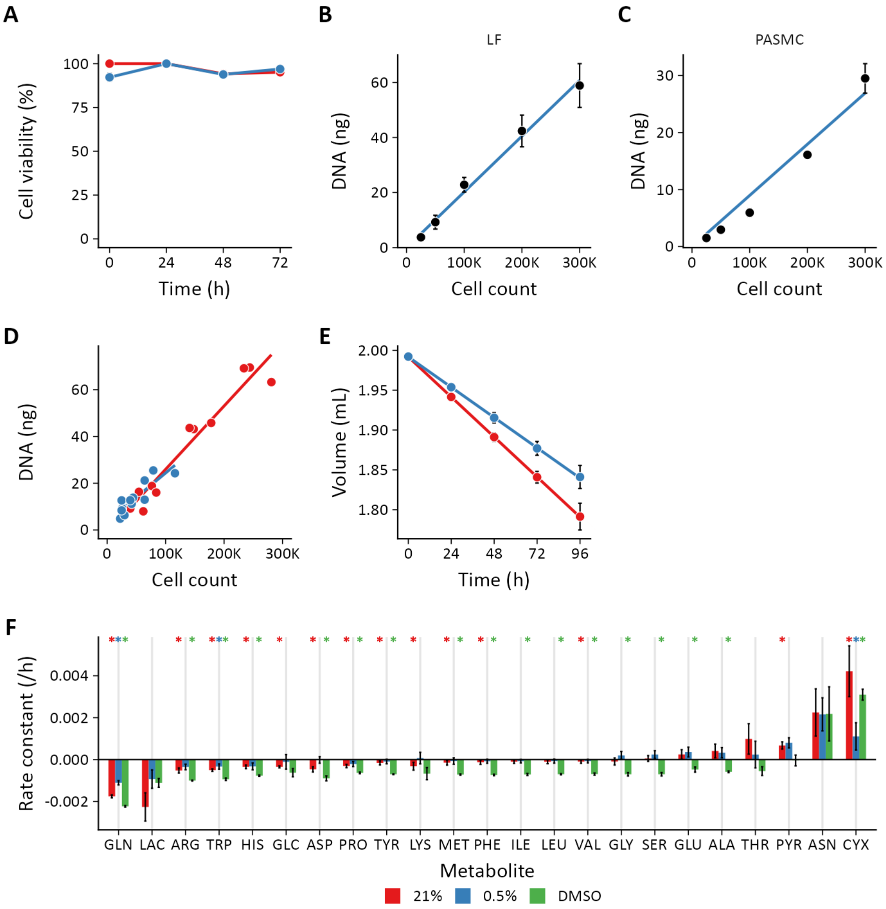


Figure 1: **Supporting data for extracellular flux calculations.** (**A**) Cell viability as assessed by live/dead cell staining with acridine orange and propidium iodide did not differ between 21% and 0.5% oxygen culture conditions (n = 3 technical replicates). (**B**) Standard curve of lung fibroblast (LF) cell count *v.* total DNA by PicoGreen measurement used to interpolate cell numbers from DNA measurements. Data are mean ± SEM of three biological replicates. (**C**) Standard curve of PASMC cell count *v.* total DNA as in (B). (**D**) Total DNA measurements were compared to direct cell counts over the experimental time course. Cell counts and total DNA were obtained from the same sample wells. The slopes of the best-fit lines for 21% (*red*) and 0.5% (*blue*) samples were not statistically different. (**E**) Predicted well volumes were estimated from the change in culture plate mass over the experimental time course. Evaporation rates were different depending on the culture conditions and treatment. Although the mean evaporation rate is depicted, experiment-specific evaporation rates were used to calculate fluxes for each biological replicate (**F**) Metabolite accumulation (positive values) and degradation (negative values) rates. Data are mean ± SEM of 3-8 biological replicates. Rates significantly different from 0 (*) based on a probability value < 0.05 using Student's one-sample *t*-test were incorporated into flux calculations.

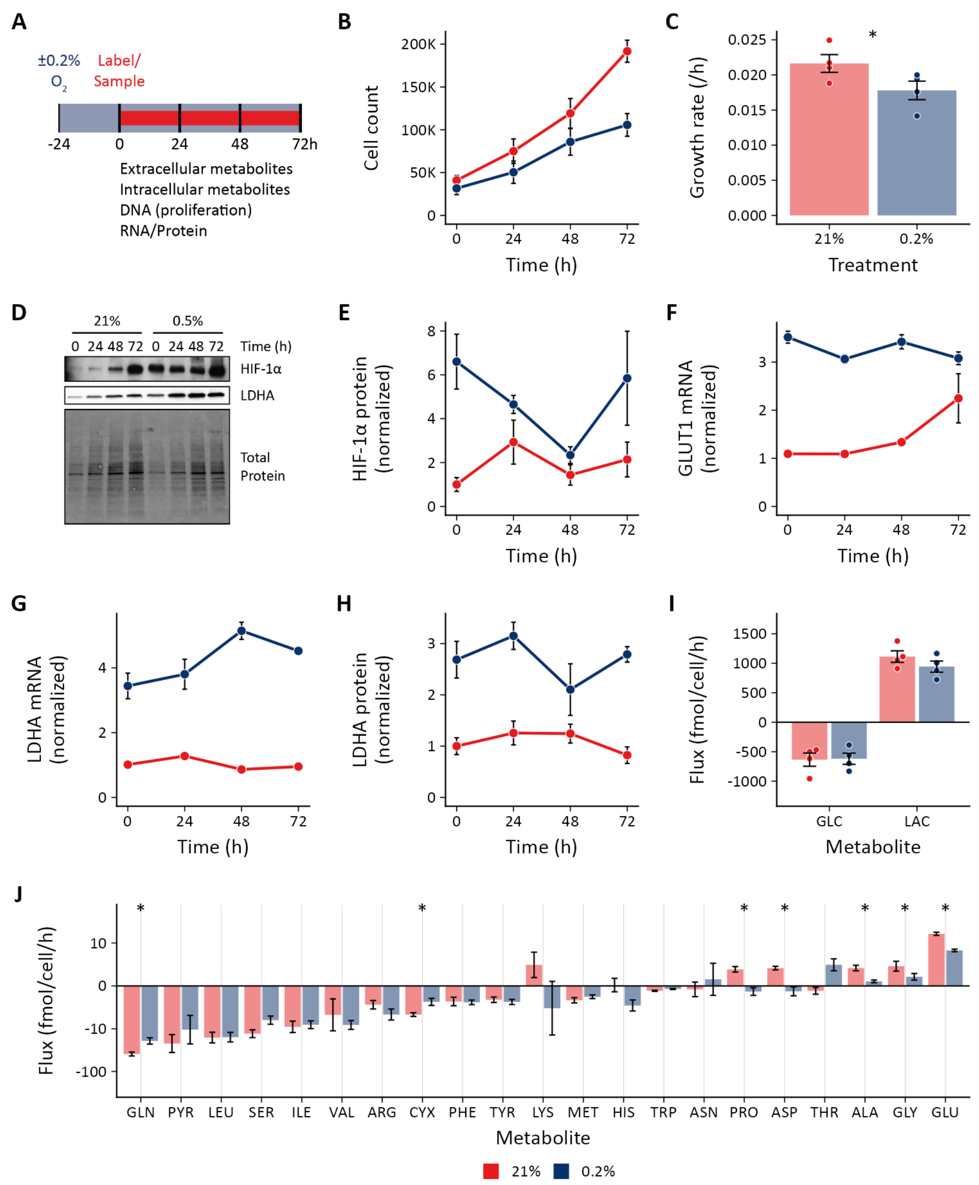


Figure 2: **Effects of hypoxia on extracellular metabolite fluxes in lung fibroblasts.** (**A**) Lung fibroblasts (LFs) were cultured in 21% or 0.2% oxygen beginning 24 h prior to time 0. Samples were collected every 24 h for 72 h. (**B**) Growth curves of LFs in each experimental condition (n = 4). (**C**) Growth rates from (B) were determined by robust linear modeling of log-transformed growth curves. (**D**) Representative immunoblot of LF protein lysates cultured as in (A). (**E**) Relative change in HIF-1α protein levels from (D) normalized to 21% oxygen at time 0 (n = 4). (**F**) Relative change in GLUT1 mRNA levels normalized to 21% oxygen treatment at time 0 (n = 4). (**G**) Relative change in LDHA mRNA levels as in (F). (**H**) Relative change in LDHA protein levels as in (E). (**I**) Extracellular fluxes of glucose (GLC) and lactate (LAC) (n = 4). By convention, negative fluxes indicate metabolite consumption. (**J**) Extracellular fluxes of pyruvate (PYR) and amino acids. Data are mean ± SEM (* $p < 0.05$).

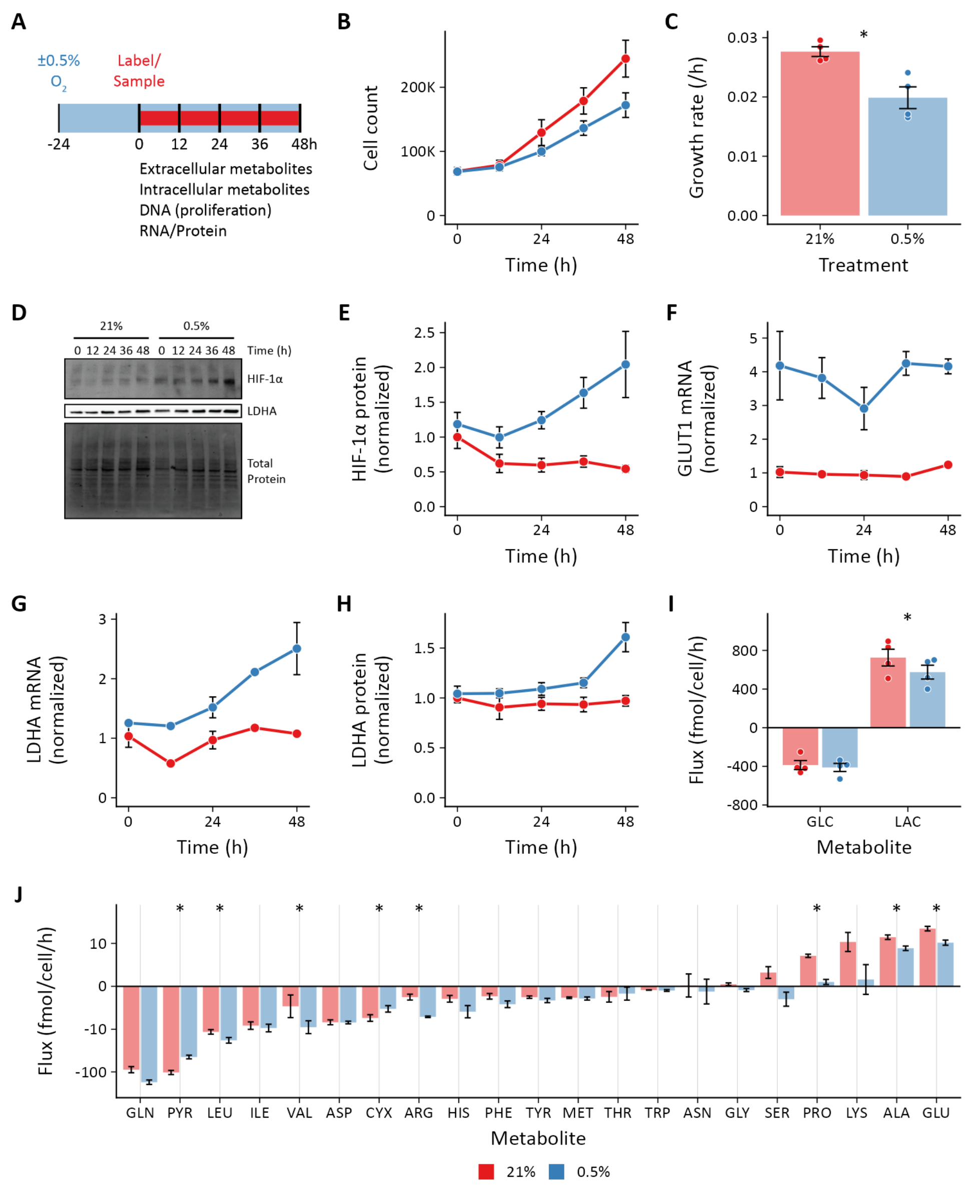


Figure 3: **Effects of hypoxia on extracellular metabolite fluxes in pulmonary artery smooth muscle cells.** (**A**) Pulmonary artery smooth muscle cells (PASMCs) were cultured in 21% or 0.5% oxygen beginning 24 h prior to time 0. Samples were collected every 12 h for 48 h. (**B**) Growth curves of LFs in each experimental condition (n = 4). (**C**) Growth rates from (B) were determined by robust linear modeling of log-transformed growth curves. (**D**) Representative immunoblot of LF protein lysates cultured as in (A). (**E**) Relative change in HIF-1α protein levels from (D) normalized to 21% oxygen at time 0 (n = 4). (**F**) Relative change in GLUT1 mRNA levels normalized to 21% oxygen treatment at time 0 (n = 4). (**G**) Relative change in LDHA mRNA levels as in (F). (**H**) Relative change in LDHA protein levels as in (E). (**I**) Extracellular fluxes of glucose (GLC) and lactate (LAC) (n = 4). By convention, negative fluxes indicate metabolite consumption. (**J**) Extracellular fluxes of pyruvate (PYR) and amino acids. Data are mean ± SEM (* $p < 0.05$).

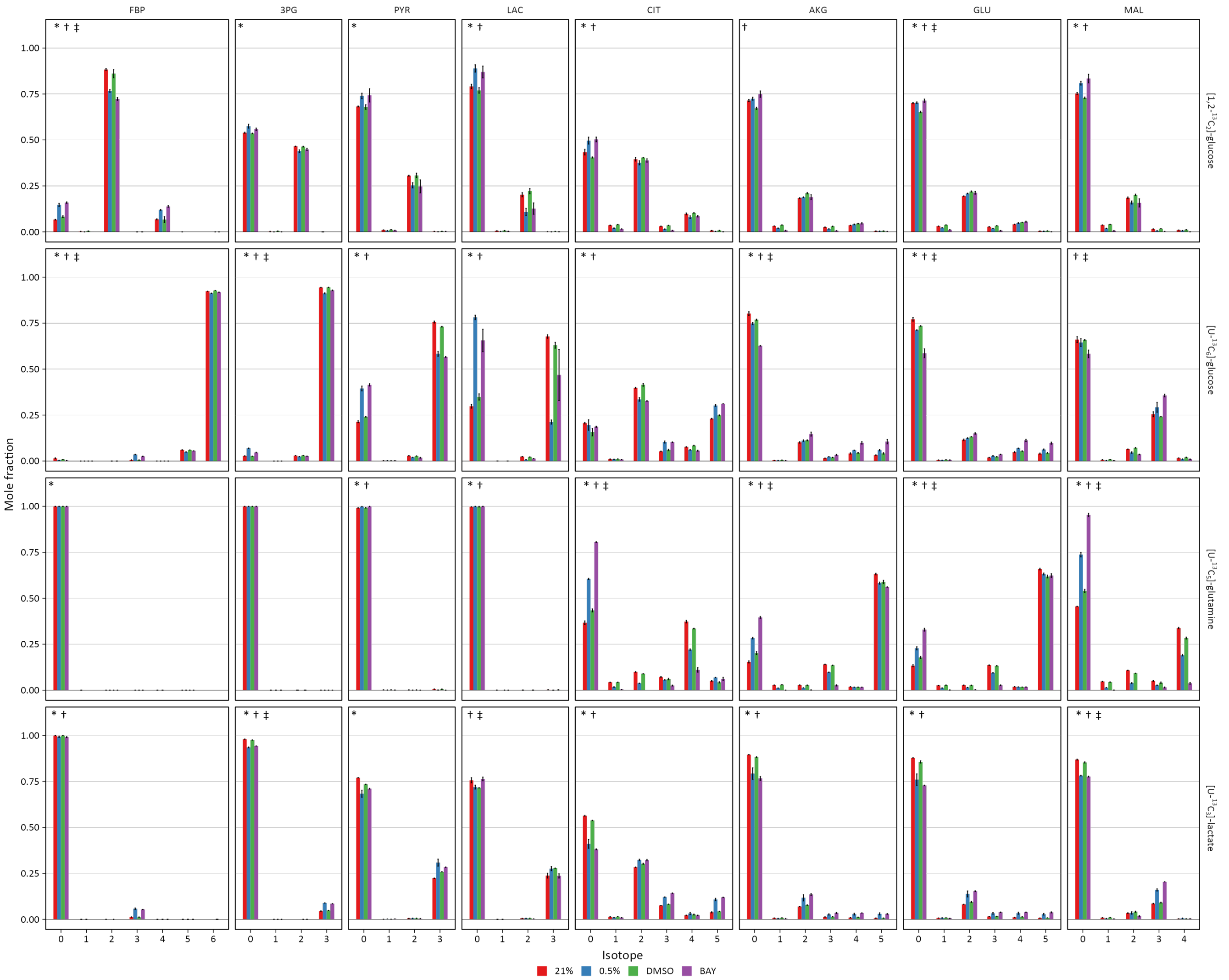


Figure 4: **Mass isotopomer distributions after 72 h of labeling in lung fibroblasts.** Lung fibroblasts (LFs) were labeled with the indicated tracers and intracellular metabolites were analyzed by LC-MS after 72 h. Mass isotopomer distributions were adjusted for natural abundance. Data are the mean ± SEM of 4 biological replicates. Significant differences in labeling patterns between 21% and 0.5% oxygen (*), DMSO and BAY treatment (†), and 0.5% oxygen and BAY treatment (‡) for each combination of metabolite and tracer are highlighted.

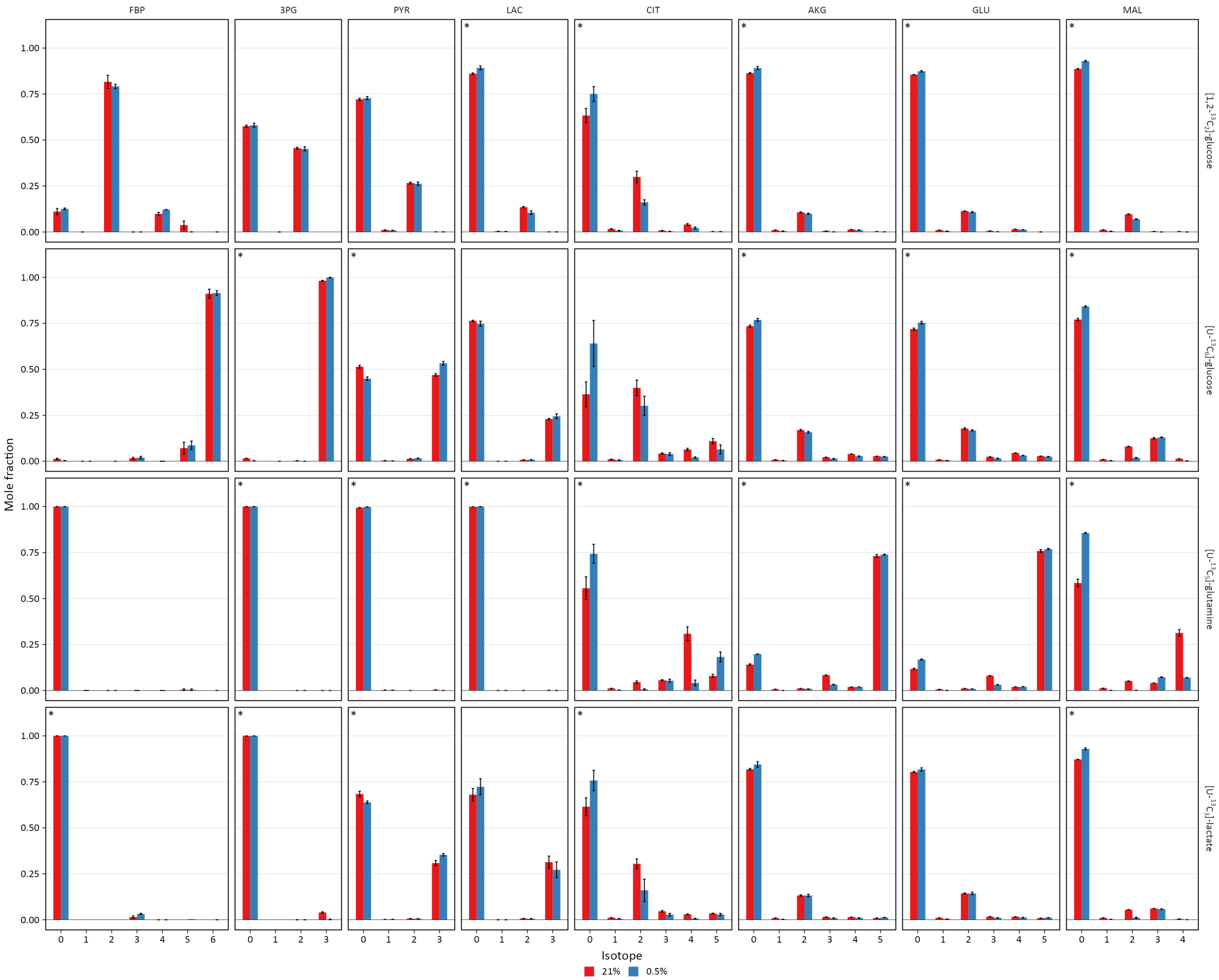


Figure 5: **Mass isotopomer distributions after 48 h of labeling in pulmonary artery smooth muscle cells.** Pulmonary artery smooth muscle cells (PASMCs) were labeled with the indicated tracers and intracellular metabolites were analyzed by LC-MS after 36 h. Mass isotopomer distributions were adjusted for natural abundance. Data are the mean ± SEM of 4 biological replicates. Significant differences in labeling patterns between 21% and 0.5% oxygen (*) for each combination of metabolite and tracer are highlighted.

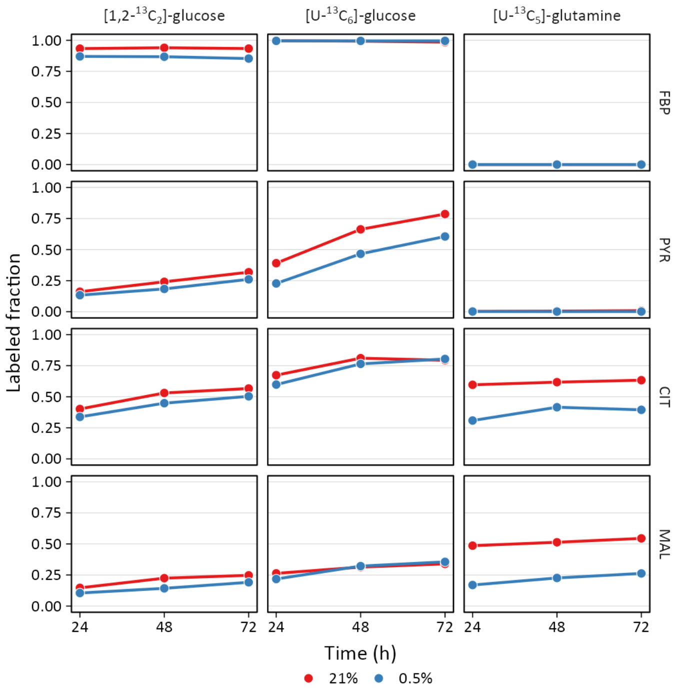


Figure 6: **Isotope incorporation over the labeling time course.** LFs were cultured in 21% or 0.5% oxygen and labeled with the indicated tracers. Intracellular metabolites were analyzed by LC-MS (FBP, fructose-bisphosphate; PYR, pyruvate; CIT, citrate; MAL, malate). Mass isotopomer distributions were calculated and adjusted for natural abundance. Data show the total amount of metabolite labeling (*i.e.,* 1 - M0 fractional abundance). Data are the mean ± SEM of 4 biological replicates.

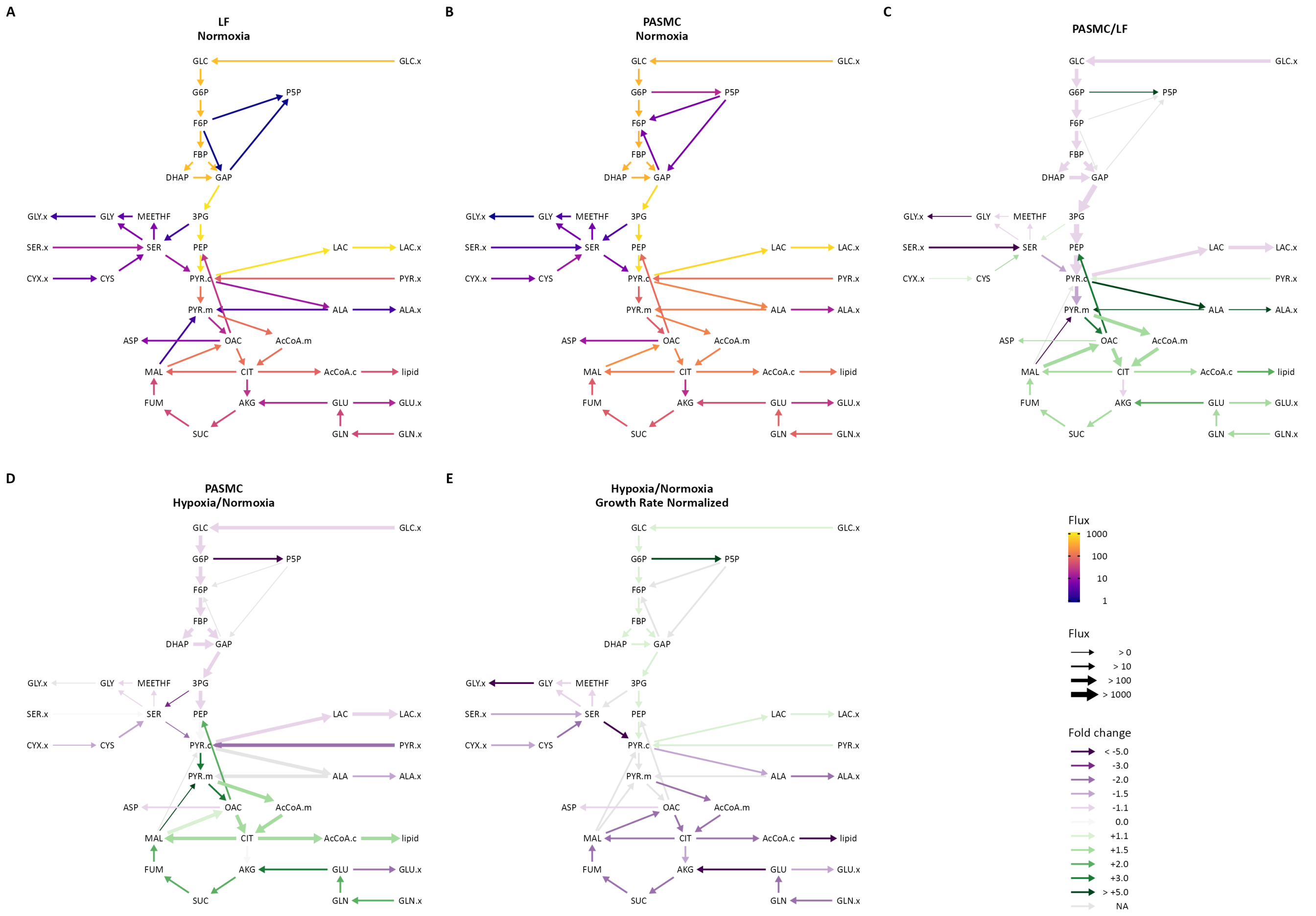


Figure 7: **Isotopically non-stationary metabolic flux analysis.** (**A**) Metabolic flux model of LF metabolism in 21% oxygen. Arrows are colored by $\log_{10}$(flux). (**B**) Metabolic flux model of PASMC metabolism in 21% oxygen as in (A). (**C**) Ratio of metabolic fluxes in PASMCs compared to LFs. Fluxes with non-overlapping confidence intervals are highlighted with arrows colored according to the magnitude of the change. Arrow thickness corresponds to the absolute flux measured in LFs. (**D**) Ratio of metabolic fluxes in 0.5% oxygen compared to 21% oxygen in PASMCs. (**E**) LF fluxes were normalized to cell growth rate. Graph depicts the ratio of normalized metabolic fluxes in LFs cultured in 0.5% oxygen compared to 21% oxygen control. Fluxes with non-overlapping confidence intervals are highlighted to indicate significant changes.

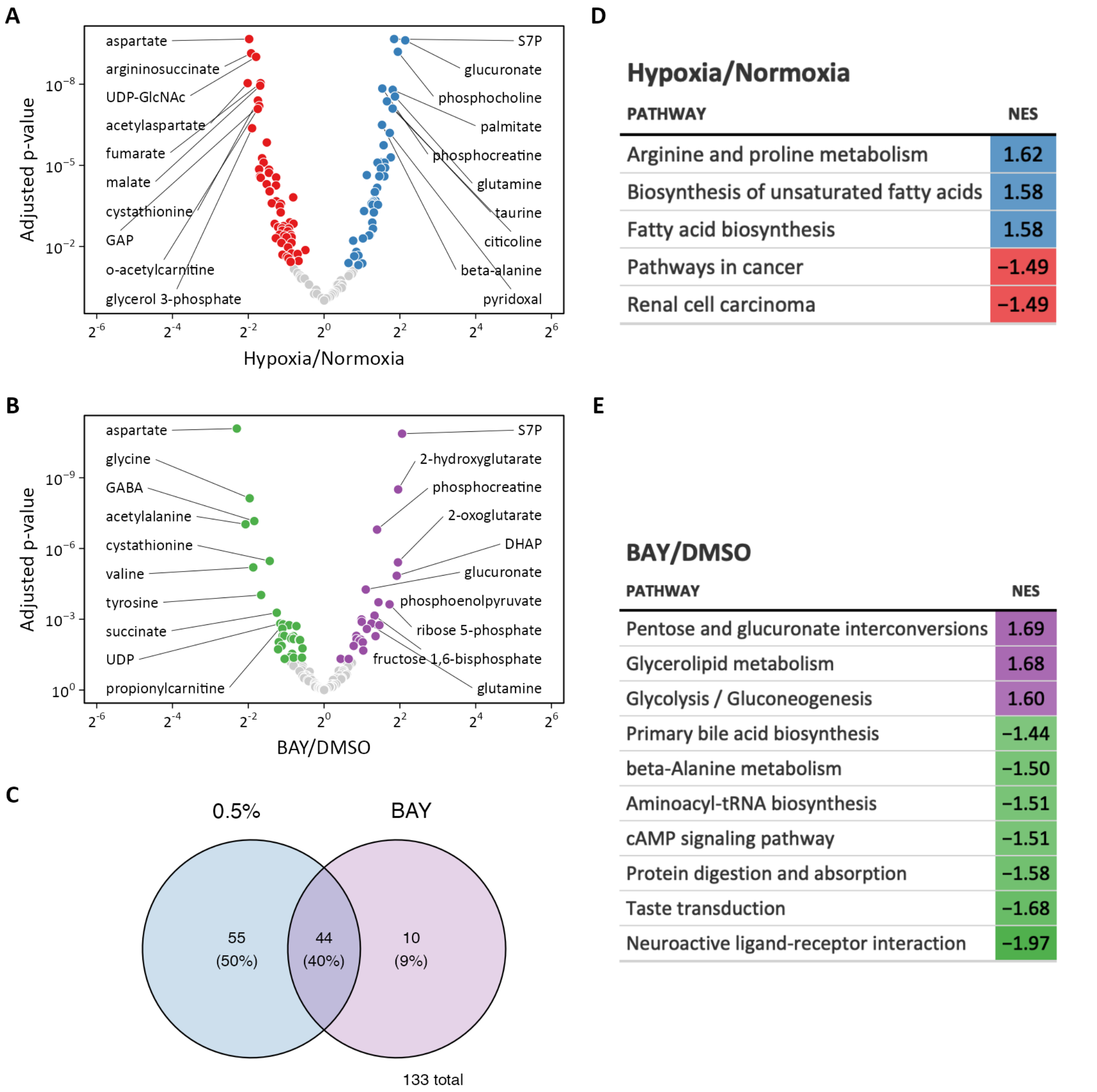


Figure 8: **Metabolomic profiling of hypoxia and BAY treated lung fibroblasts.** (**A**) Volcano plot of differentially regulated metabolites following 0.5% oxygen culture. (**B**) Volcano plot of differentially regulated metabolites by BAY treatment in 21% oxygen. (**C**) Venn diagram illustrating the overlap among metabolites differentially regulated by hypoxia (*blue*) or BAY treatment (*purple*). (**D**) Metabolite enrichment in KEGG pathways following hypoxia. (**E**) Metabolite enrichment in KEGG pathways following BAY. Only significantly enriched metabolite sets with $p < 0.05$ are shown. NES, normalized enrichment score.

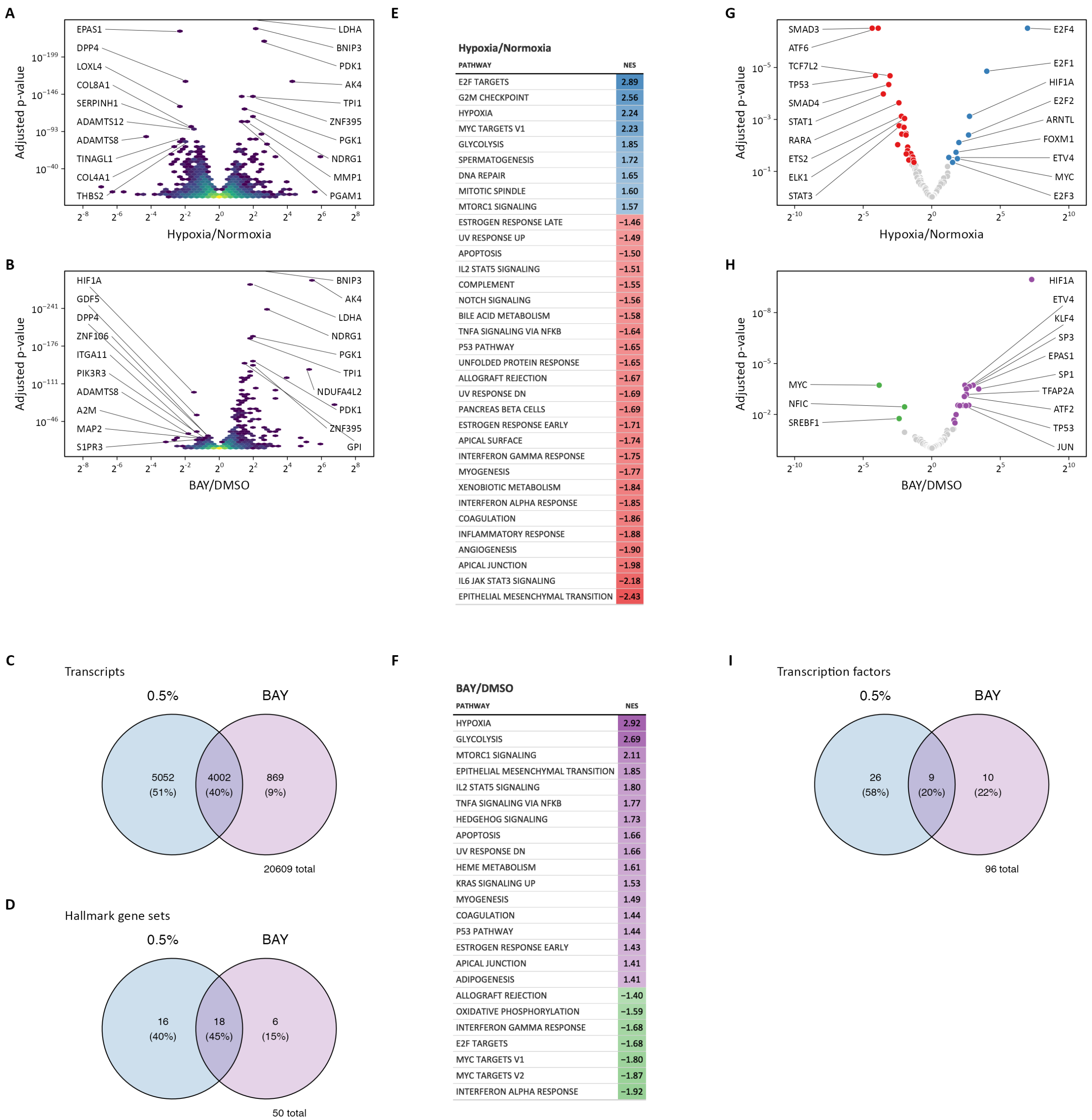


Figure 9: **Transcriptomic profiling of hypoxia and BAY treated lung fibroblasts.** (**A**) Volcano plot of differentially expressed genes following 0.5% oxygen culture. (**B**) Volcano plot of differentially expressed genes following BAY treatment. (**C**) Venn diagram illustrating the number of differentially expressed transcripts following hypoxia (*blue*) or BAY treatment (*purple*). (**D**) Venn diagram illustrating the number of differentially enriched Hallmark gene sets with hypoxia (*blue*) or BAY treatment (*purple*). (**E**) Gene set enrichment results of hypoxia-treated cells. (**F**) Gene set enrichment results of BAY-treated cells. All gene sets listed were significantly enriched at FDR < 0.05. NES, normalized enrichment score. (**G**) Volcano plot illustrating the results of a transcription factor enrichment analysis in hypoxia-treated cells. (**H**) Volcano plot illustrating the results of a transcription factor enrichment analysis in BAY-treated cells. (**I**) Venn diagram illustrating the overlap among enriched transcription factors following hypoxia or BAY treatment.

#### REFERENCES

##### Software


Allaire J, Xie Y, McPherson J, Luraschi J, Ushey K, Atkins A, Wickham H, Cheng J, Chang W & Iannone R (2022) Rmarkdown: Dynamic documents for r

Bache SM & Wickham H (2022) Magrittr: A forward-pipe operator for r

Bates D, Mächler M, Bolker B & Walker S (2015) Fitting linear mixed-effects models using lme4. *Journal of Statistical Software* 67: 1–48

Bates D, Maechler M, Bolker B & Walker S (2022) lme4: Linear mixed-effects models using eigen and S4

Garnier S (2021) Viridis: Colorblind-friendly color maps for r

Grolemund G & Wickham H (2011) Dates and times made easy with lubridate. *Journal of Statistical Software* 40: 1–25

Halekoh U & Højsgaard S (2014) A kenward-roger approximation and parametric bootstrap methods for tests in linear mixed models – the R package pbkrtest. *Journal of Statistical Software* 59: 1–30

Halekoh U & Højsgaard S (2021) Pbkrtest: Parametric bootstrap, kenward-roger and satterthwaite based methods for test in mixed models

Henry L & Wickham H (2020) Purrr: Functional programming tools

Henry L & Wickham H (2022) Rlang: Functions for base types and core r and tidyverse features

Kuznetsova A, Brockhoff PB & Christensen RHB (2017) lmerTest package: Tests in linear mixed effects models. *Journal of Statistical Software* 82: 1–26

Kuznetsova A, Bruun Brockhoff P & Haubo Bojesen Christensen R (2020) lmerTest: Tests in linear mixed effects models

Lenth RV (2022) Emmeans: Estimated marginal means, aka least-squares means

Müller K & Wickham H (2022) Tibble: Simple data frames

Neuwirth E (2022) RColorBrewer: ColorBrewer palettes

Oldham W (2022) Wmo: Personal utility functions

Ooms J (2021) Magick: Advanced graphics and image-processing in r

Pedersen TL (2020) Patchwork: The composer of plots

R Core Team (2022) R: A language and environment for statistical computing Vienna, Austria: R Foundation for Statistical Computing

Ripley B (2022) MASS: Support functions and datasets for venables and ripley's MASS

Spinu V, Grolemund G & Wickham H (2021) Lubridate: Make dealing with dates a little easier

Ushey K (2022) Renv: Project environments

Venables WN & Ripley BD (2002) Modern applied statistics with s Fourth. New York: Springer

Wickham H (2016) ggplot2: Elegant graphics for data analysis Springer-Verlag New York

Wickham H (2019) Stringr: Simple, consistent wrappers for common string operations

Wickham H (2021a) Forcats: Tools for working with categorical variables (factors)

Wickham H (2021b) Tidyverse: Easily install and load the tidyverse

Wickham H, Averick M, Bryan J, Chang W, McGowan LD, François R, Grolemund G, Hayes A, Henry L, Hester J, *et al* (2019) Welcome to the tidyverse. *Journal of Open Source Software* 4: 1686

Wickham H & Bryan J (2022) Readxl: Read excel files

Wickham H, Bryan J & Barrett M (2022a) Usethis: Automate package and project setup

Wickham H, Chang W, Henry L, Pedersen TL, Takahashi K, Wilke C, Woo K, Yutani H & Dunnington D (2022b) ggplot2: Create elegant data visualisations using the grammar of graphics

Wickham H, Danenberg P, Csárdi G & Eugster M (2022c) roxygen2: In-line documentation for r

Wickham H, François R, Henry L & Müller K (2022d) Dplyr: A grammar of data manipulation

Wickham H & Girlich M (2022) Tidyr: Tidy messy data

Wickham H, Hester J & Bryan J (2022e) Readr: Read rectangular text data

Wickham H, Hester J, Chang W & Bryan J (2021) Devtools: Tools to make developing r packages easier

Xie Y (2015) Dynamic documents with R and knitr 2nd ed. Boca Raton, Florida: Chapman; Hall/CRC

Xie Y (2016) Bookdown: Authoring books and technical documents with R markdown Boca Raton, Florida: Chapman; Hall/CRC

Xie Y (2022a) Bookdown: Authoring books and technical documents with r markdown

Xie Y (2022b) Knitr: A general-purpose package for dynamic report generation in r

Xie Y (2022c) Tinytex: Helper functions to install and maintain TeX live, and compile LaTeX documents

Xie Y, Allaire JJ & Grolemund G (2018) R markdown: The definitive guide Boca Raton, Florida: Chapman; Hall/CRC

Zhu H (2021) kableExtra: Construct complex table with kable and pipe syntax
